# Proteostasis dysregulation in p.A53T-α-Synuclein iPSC-derived astrocytes exacerbates neurodegeneration in a Parkinson’s disease model with Lewy-like pathology

**DOI:** 10.1101/2024.11.06.621638

**Authors:** Christina Paschou, Olympia Apokotou, Anastasios Kollias, Konstantina Charmpi, Sofia Dede, Martina Samiotaki, Francesca Palese, Konstantina Dimoula, Evangelia Emmanouilidou, Era Taoufik, Chiara Zurzolo, Rebecca Matsas, Florentia Papastefanaki

## Abstract

Alpha-Synuclein (αSyn) plays a central role in Parkinson’s disease (PD) and the p.A53T mutation causes an early-onset familial form of PD with severe manifestations. The pathological effects of the p.A53T-αSyn mutation have been extensively investigated in neurons, yet the consequences on astrocytes and astrocytic contribution to PD pathology are understudied. Here, we differentiated induced pluripotent stem cells from PD patients carrying the p.A53T-αSyn mutation to astrocytes, which uncovered cell-intrinsic phenotypes, including calcium dyshomeostasis and accumulation of protein aggregates. Proteomic profiling and functional analyses revealed perturbed protein catabolic processes, involving the proteasome and autophagy, associated with lysosomal malfunction. Dopamine neurons co-cultured with p.A53T-αSyn astrocytes displayed exacerbated neurodegeneration with hallmark Lewy-like pathologies, reversed by control astrocytes at least due to their ability to resolve neuronal αSyn aggregates by endocytic clearance. Our findings underscore a critical impact of p.A53T-αSyn on astrocytic protein quality control mechanisms, positioning astrocytes as important contributors to PD neuropathology.

**Highlights:**

- iPSC-derived astrocytes from PD patients with the p.A53T-αSyn mutation display cell-autonomous pathological phenotypes and toxic αSyn accumulation
- Proteome and functional analyses reveal failure of major proteostasis mechanisms in p.A53T-αSyn astrocytes, largely related to lysosomal malfunction
- Inherent Lewy-like pathological features are identified in p.A53T-αSyn astrocyte-neuron co-cultures
- p.A53T-αSyn astrocytes induce PD-relevant neuropathology to healthy neurons
- Control but not p.A53T-αSyn astrocytes alleviate neuropathology in co-cultured neurons

## Introduction

Synucleinopathies, including Parkinson’s disease (PD), Lewy body dementia (LBD), multiple system atrophy, and REM sleep behavior disorder, comprise a spectrum of neurodegenerative disorders defined by intracellular accumulation and aggregation of alpha-synuclein (αSyn) in neurons and glia^1^. Although the physiological function of αSyn is not fully understood, its primary suggested role involves regulation of vesicle trafficking and neurotransmitter recycling at presynaptic terminals, where it is mainly located^2–4^. In the brain, αSyn is predominantly expressed in neurons and to a far lesser extent in glial cells^5^, with its levels depending on the balance between synthesis, aggregation, clearance, and secretion^4^. Misfolded αSyn forms pathological aggregates that propagate among brain regions and cell populations, in a prion-like manner^6,7^. Post-mortem PD and LBD brain specimens typically exhibit intraneuronal protein inclusions rich in aggregated αSyn, termed Lewy bodies (LB) and Lewy neurites (LN)^8–10^. Indeed, the *SNCA* gene encoding for αSyn was the first gene associated with both sporadic and familial PD, with the autosomal dominant G209A mutation (encoding for the pathological p.A53T-αSyn protein) rendering αSyn more prone to aggregation and leading to early-onset and severe disease^11^.

Accumulation of neurotoxic αSyn-rich inclusions results in gradual loss of dopaminergic neurons (DAn) projecting to the striatum from the substantia nigra pars compacta^12^. Therefore, research in PD has traditionally focused on neuron-intrinsic deficits. However, there is now emerging interest in the contribution of glial cells to neurodegenerative diseases, including PD, given their critical roles in maintaining brain homeostasis and supporting physiological neuronal functions. Astrocytes in particular, the most abundant glial cells in human brain, are essential for clearance of neuronal-derived damaged or aggregated proteins, such as αSyn. Hence, astrocyte dysfunctions could participate to the development and progression of PD pathology^13,14^. Notably, intracellular deposits of aggregated αSyn have been detected within astrocytes^15^. Astrocytic αSyn-immunoreactive inclusions were reported in sporadic PD autopsies, exhibiting cortical^5^ and nigral^16^ distribution, but also in other brain regions, displaying a unique signature of post-translational modifications^17^ and morphology distinct from neuronal Lewy pathology^15^. Alpha-synuclein in astrocytes may originate from endogenous expression, uptake of neuronal αSyn, or both. Alpha-synuclein transfer has been reported *in vitro* from neurons to astrocytes^18–21^ and may trigger an inflammatory response, impair the protein degradation systems, and cause mitochondrial and endoplasmic reticulum (ER) dysfunctions^22^.

Several PD-related genes are expressed in astrocytes and their risk variants, aside from affecting neurons, alter astrocytic function^14^. Cell reprogramming has enabled the generation of human-relevant *in vitro* systems through differentiation of induced pluripotent stem cells (iPSC) from patients with sporadic or familial PD^23–31^. Previous studies, including ours, have shown disease-associated phenotypes in iPSC-derived neurons^23,28,30,32^. More recently, iPSC-derived astrocytes from PD patients, harboring mutations in Leucine-rich repeat kinase 2 (*LRRK2*), Glucosylceramidase beta 1 (*GBA1*), or ATPase 13A2 (*ATP13A2*) genes, have been shown to display alterations in metabolic pathways, clearance mechanisms, and neuroinflammatory responses^14,21,29,33–38^. These observations suggest an astrocyte-originating, non-cell autonomous component of PD neuropathology, requiring further investigation.

Here we sought to investigate whether p.A53T-αSyn (G209A *SNCA* mutation) causes cell-autonomous dysfunctions in astrocytes that may contribute to aggravation of neuronal pathology, in a human early-onset PD model. We differentiated iPSC lines obtained from PD patients carrying the p.A53T-αSyn mutation and healthy controls^30^, as well as an isogenic pair of p.A53T-αSyn patient and gene-corrected iPSC, toward ventral midbrain (vm) astrocytes and characterized them for disease-relevant phenotypes, along with their proteome profiling. Our analysis revealed toxic αSyn accumulation and cell-intrinsic pathology affecting proteostasis. Major clearance mechanisms, including the proteasome, autophagy and endo-lysosomal pathways were perturbed in astrocytes. In a co-culture system of iPSC-derived control or PD dopaminergic neurons with PD astrocytes, we observed exacerbated neurodegeneration with Lewy-like pathology, mirroring the histopathological hallmarks identified in post-mortem PD brains. These phenotypes were rescued by control astrocytes, underscoring their critical involvement in preserving neuronal integrity.

## Results

### Differentiation of iPSCs into ventral midbrain astrocytes reveals p.A53T-αSyn-related functional aberrations

Ventral midbrain (vm)-regionalized astrocytes were generated from previously characterized iPSCs from two patients bearing the p.A53T-αSyn mutation^30^, using established protocols^39,40^ (Fig. 1a). Age- and sex-matched control iPSCs from two healthy donors (non-PD), as well as an isogenic pair of p.A53T-αSyn patient-derived iPSCs and its gene-corrected control were also included^25^ (Supplementary Table 1). Following iPSC neuralization via dual SMAD inhibition, vm-patterned neural precursor cells (NPCs) were obtained (Fig. 1a and Supplementary Fig. 1). After 28 days of astrocytic differentiation, practically all iPSC-derived cells displayed expression of astroglial markers, including CD44, Vimentin, Aldehyde Dehydrogenase 1 Family Member L1 (ALDH1L1), and S100β, with no apparent differences between genotypes (healthy and p.A53T-αSyn iAstrocytes, hereafter designated Ha and PDa, respectively) (Supplementary Fig. 2a). More mature astrocytic markers, including aquaporin 4 (AQP4), Excitatory amino acid transporter 1 (EAAT1), and CD49f, were enriched after 2-w maturation treatment with bone morphogenetic protein-4 (BMP-4) and ciliary neurotrophic factor (CNTF) (Fig. 1a, Supplementary Fig. 2b). Whole proteome profiling of Ha and PDa (Supplementary Table 2) followed by gene set enrichment analysis in the MSig C8 database^41^ revealed at least two astrocyte-related gene sets significantly enriched in highly expressed proteins while none of the brain neuron-related nor oligodendrocyte-related gene sets were significantly enriched (Supplementary Fig. 3). Moreover, by immunofluorescence for beta-3-tubulin or MAP2 we could not detect neuronal cells in the astrocyte cultures.

**Figure 1.**
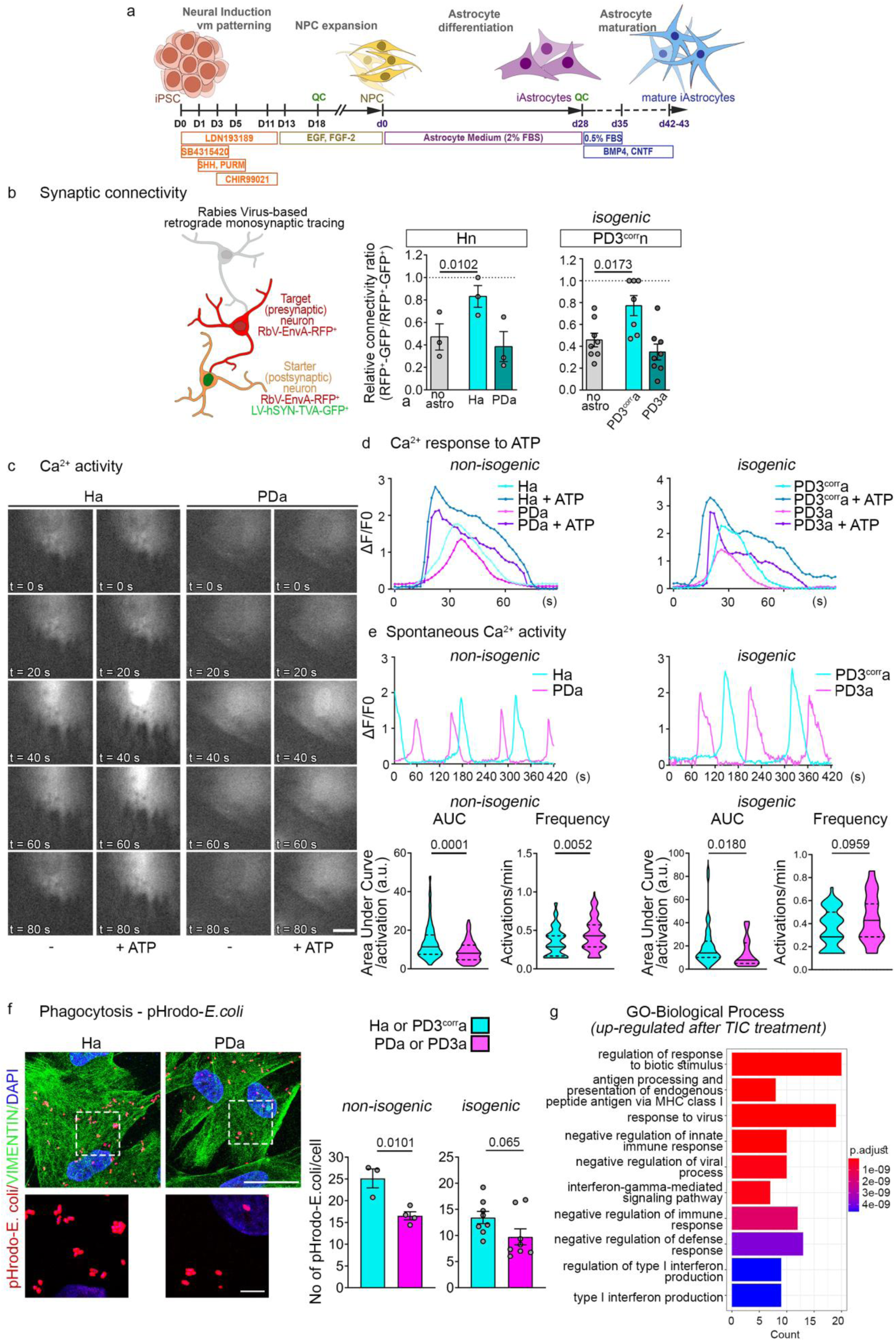
Functional characterization of healthy and p.A53T-αSyn iPSC-derived astrocytes. **a.** Schematic of the differentiation protocol of iPSC into ventral midbrain astrocytes. **b** (Left) Drawing of the rabies virus-based retrograde monosynaptic tracing system. Target (presynaptic) neurons are RbV-ENVA-RFP^+^/LV-TVA-GFP^-^ and starting (postsynaptic) neurons are RbV-ENVA-RFP^+^/LV-TVA-GFP^+^. (Right) Bar plots showing the relative connectivity ratio in differentiation day 40 healthy neurons (Hn) alone or co-cultured with Ha or PDa (left) and the isogenic pair PD3^corr^a and PD3a (right). **c** Time series of spontaneous intracellular Ca^2+^ responses visualized using Fluo4AM in live Ha and PDa, and after 50-μM ATP stimulation. **d** Line plots of responses (ΔF/F0 over time [s]) of one representative Ha and one PDa cell before and after ATP treatment. **e** Line plots of spontaneous Ca^2+^ responses (ΔF/F0 over time [s]) of one representative Ha and one PDa cell (left, non-isogenic; right, isogenic). Violin plots show the distribution of the AUCs and frequencies of spontaneous Ca^2+^ responses normalized to average basal response after background subtraction in Ha and PDa or the isogenic pair. **f** Representative confocal images of Ha and PDa treated with pHRodo-*E. coli* to monitor phagocytosis, immunostained for Vimentin, with DAPI. Scale bar: 30 μm and 5 μm for insets. Bar plots show the average number of pHRodo^+^ puncta per cell in Ha and PDa (left) or the isogenic pair (right). **g** Bar plot showing the top 10 most significant up-regulated GOBP terms after TIC treatment. The length of each bar corresponds to the number of up-regulated proteins associated with that term, while the color intensity reflects the p-value. Data are presented as mean ± SEM (c, g) or median with interquartile range (f). One-way ANOVA with Greenhouse-Geisser correction and Dunnett’s multiple comparisons test was applied in b, and unpaired two-tailed t-tests were used for comparisons in f; Mann-Whitney tests were used in e. Sample sizes: b, n = 3 Hn lines, n = 8 FOV in PD3^corr^n, 7 FOV in PD3^corr^n-PD3^corr^a, and 8 FOV in PD3^corr^n-PD3a co-cultures; e, n = 135 Ha and 106 PDa, across 3 Ha and 3 PDa non-isogenic lines, in separate experiments; n = 45 PD3^corr^a and 32 PD3a; f, n = 3 Ha and 4 PDa non-isogenic lines; n = 8 FOV in PD3^corr^a and 8 FOV in PD3a. *AUC, Area Under Curve; FOV, Field of View; GOBP, Gene Ontology Biological Process; LV, lentiviral vector; RbV, Rabies Virus; SEM, standard error of the mean*.

Investigation of the properties of iAstrocytes verified the functional competence of Ha and revealed cell-intrinsic dysfunctional phenotypes in PDa. Using a rabies virus-based retrograde monosynaptic tracing system^42^ to assess the relative neuronal connectivity index, which reflects the ratio of target neurons (RFP^+^/GFP^-^) retrogradely labeled through monosynaptic connections with starter neurons, over starter neurons (RFP^+^/GFP^+^), we noted a significant increase in synaptic connectivity of iPSC-derived healthy neurons (iNeurons, Hn) co-cultured with Ha – but not with PDa – as compared to Hn cultured without astrocytes for the same duration (Fig. 1b). This indicates an essential supportive function of healthy as opposed to PD astrocytes on the ability of healthy neurons to form synaptic connections.

The capacity of iAstrocytes to elicit spontaneous and ATP-evoked Ca^2+^ transients was captured by live cell imaging using Fluo4AM (Fig. 1c-e). PDa exhibited significantly smaller area under the curve (AUC) in spontaneous Ca^2+^ activity compared to Ha, while the frequency of responses showed a significant increase. The aberrations in Ca^2+^ activity are indicative of disruption in Ca^2+^ homeostasis and potential dysfunction in cellular compartments, including the ER, mitochondria, and the endo-lysosomal pathway, that have been implicated in neurodegeneration^43^.

The phagocytic activity of iAstrocytes was tested by exposure for 6 h to pHRodo-*E. coli* bioparticles that fluoresce in the acidic environment of intracellular compartments (Fig. 1f). Both Ha and PDa displayed internalization ability, yet significantly fewer pHRodo^+^ puncta were detected in the cytoplasm of PDa, compared to Ha, indicating defective endocytosis (Fig. 1f).

Last, we examined the responsiveness of iAstrocytes to pro-inflammatory signals, such as the “TIC” cytokine cocktail, comprising tumor necrosis factor-alpha (TNFα), interleukin-1 alpha (IL-1α), and complement component C1q^44,45^. Both Ha and PDa responded to TIC-treatment, as revealed by proteomic analyses. Principal component analysis (PCA) demonstrated clustering of samples by disease on the first principal component and by treatment on the second (Supplementary Fig. 4). Gene Ontology Biological Process (GOBP) enrichment based on the 113 most up-regulated proteins identified in TIC-treated samples, revealed that the top up-regulated terms were involved in immune response and inflammation processes (Fig. 1g), consistent with previous observations^45^.

Together, our data confirm the generation of highly pure and functional astrocytic populations, and provide evidence for cell-intrinsic defects in PDa, causally related to the p.A53T mutation as evidenced by concurrent analysis of the isogenic pair PD3a and its gene corrected control PD3^corr^a, which gave similar results with the non-isogenic lines PDa and Ha, respectively (Fig. 1 and throughout this study).

### Upregulation of αSyn and protein aggregation in p.A53T-αSyn iAstrocytes

The origin of αSyn inclusions in astrocytes detected in post-mortem PD brains lacks consensus, with some publications asserting absence^21,46^ and others demonstrating existence^29,34^ of inherent astrocytic αSyn expression. Here, by whole proteome profiling of Ha and PDa (Supplementary Table 2 and below), we detected SNCA among the proteins with considerable abundance (mean Log2 intensity: 20.566; ranking: 2317/7607) and levels significantly increased in PDa (Log_2_FC 0.902; p = 0.0018), compared to Ha (Fig. 2a). This increase was evident by immunofluorescence in both the non-isogenic and isogenic lines (Fig. 2b, Supplementary Fig. 5a) and quantified by Western blot at the protein level (Fig. 2c). At the transcriptional level, no significant difference was detected by RT-qPCR (Fig. 2d).

**Figure 2.**
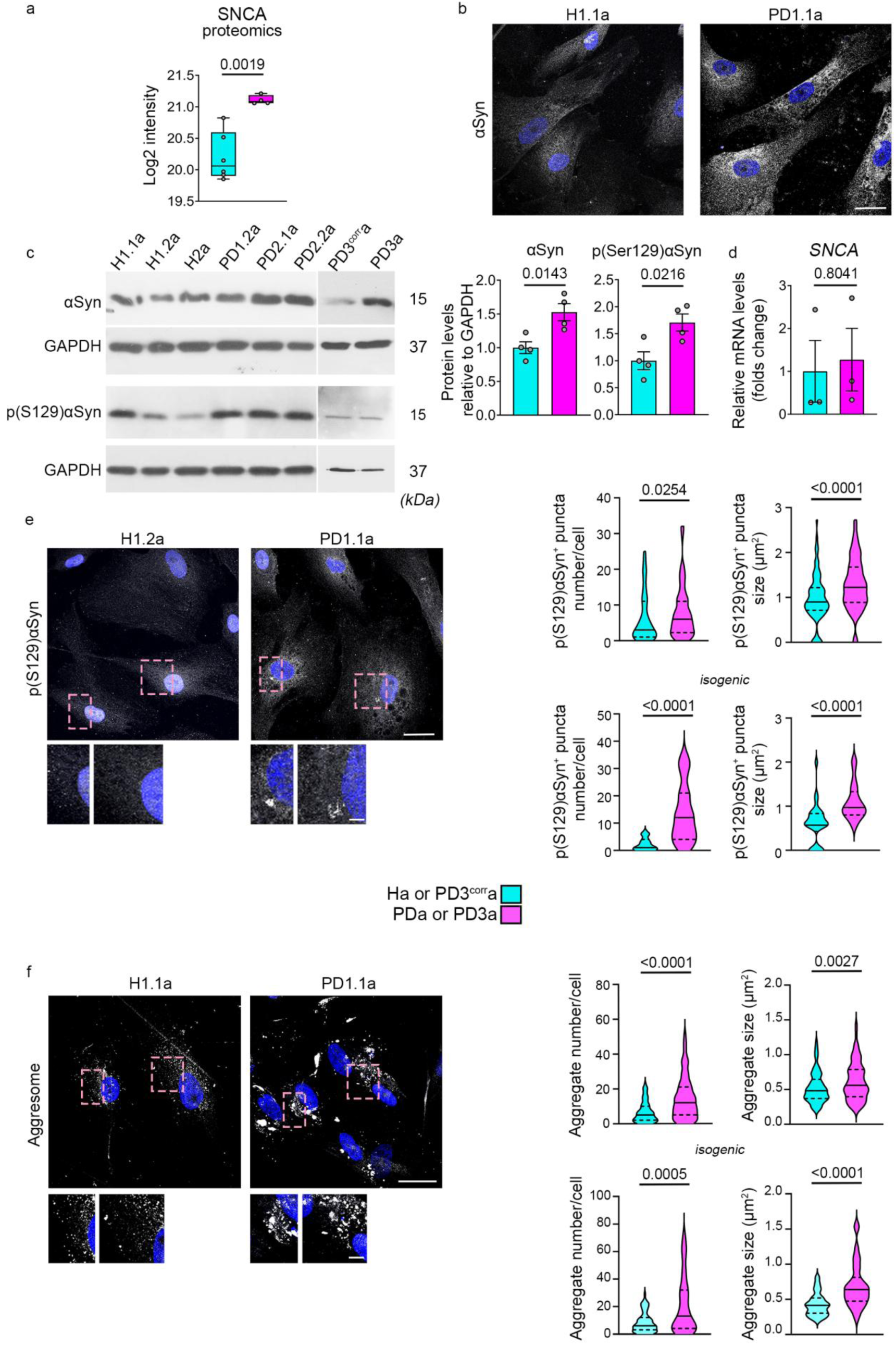
Increased αSyn, p(Ser129)αSyn, and protein aggregation in p.A53T-αSyn astrocytes. **a** Box plot showing Log2 intensity of SNCA in Ha and PDa, as detected in the proteomic analysis. **b** Representative confocal images of Ha and PDa immunostained for αSyn with DAPI. Scale bar: 30 μm. **c** Western blot analysis of αSyn, p(Ser129)αSyn, and GAPDH (loading control) in 3 Ha and 3 PDa non-isogenic lines and the isogenic pair. Bar plots represent quantification of αSyn and p(Ser129)αSyn levels normalized to GAPDH. **d** Bar plot representing mRNA levels of *SNCA* relative to *GAPDH*, in Ha and PDa. **e** Representative confocal images of Ha and PDa immunostained for p(Ser129)αSyn with DAPI. Insets show magnified views of boxed regions. Scale bars: 30 μm and 5 μm for insets. Violin plots represent the distributions of the number of p(Ser129)αSyn^+^ puncta per cell (left) and size (right) in Ha and PDa (top) and the isogenic pair (bottom). **f** Representative confocal images of Aggresome staining with DAPI in Ha and PDa. Insets show magnified views of boxed regions. Scale bar: 30 μm and 5 μm for insets. Violin plots show the distributions of the number of Aggresome^+^ puncta per cell (left) and size (right) in Ha and PDa (top) and the isogenic pair (bottom). Data are presented as median with min and max values (a), or mean ± SEM (c, d), or median with interquartile range (e, f). Unpaired two-tailed t-tests were used for comparisons in a, c, and d; Mann-Whitney tests were used in e and f. Sample sizes: a, n = 6 Ha and n = 4 PDa; c, n = 4 Ha and 4 PDa lines (including the isogenic pair); d, n = 3 Ha and 3 PDa non-isogenic lines; e, puncta number, n =81 cells across 3 non-isogenic Ha lines and 96 cells across 4 non-isogenic PDa lines. For puncta size, n = 90 cells across 3 Ha and 120 cells across 3 PDa non-isogenic lines; puncta number, n 27 cells in PD3^corr^a and 27 cells in PD3a; puncta size, n = 30 cells in PD3^corr^a and n = 28 cells in PD3a; f, aggregate number, n =180 cells across 3 Ha and 151 cells across 4 PDa non-isogenic lines; for aggregate size, n =182 cells across 3 Ha and 150 cells across 4 PDa non-isogenic lines; isogenic, aggregate number, n = 76 cells in PD3^corr^a and 71 cells in PD3a; aggregate size, n =73 cells in PD3^corr^a and 74 cells in PD3a. *αSyn, alpha-synuclein; p(Ser129)αSyn, phosphorylated alpha-synuclein at serine 129; Ha, healthy astrocytes; PDa, p.A53T-αSyn astrocytes; PD3a, p.A53T-αSyn astrocytes; PD3^corr^a, corrected isogenic astrocytes; SEM, standard error of the mean; SNCA, α-synuclein gene*.

The p.A53T mutation has been correlated with increased αSyn aggregation^47–49^ and phosphorylation at Ser129^50,51^, a post-translational modification upregulated in PD and related to neurotoxicity^51,52^. In PDa, we noted a significant increase of p(Ser129)αSyn by Western blot (Fig. 2c), as well as in the number and size of p(Ser129)αSyn^+^ puncta detected by immunofluorescence, compared to Ha (Fig. 2e, Supplementary Fig. 5b). These findings provide evidence that, in the absence of neurons, αSyn in iAstrocytes derives from intrinsic expression and that the p.A53T mutation triggers accumulation of toxic αSyn species. Accordingly, PDa possessed more and larger cytoplasmic protein aggregates compared to Ha, as identified using the Proteostat kit for aggresome detection (Fig. 2f, Supplementary Fig. 5c), suggesting dysregulation in proteostasis.

### Neuronal viability and growth are compromised in co-culture with p.A53T-αSyn astrocytes, rescued by control astrocytes

To address if the cell-intrinsic phenotypes identified in p.A53T-αSyn astrocytes affect neuronal health and contribute to PD pathology, we generated ventral midbrain dopaminergic iNeurons from the same populations of iPSC-derived vm-NPCs, as those used for astrocytic differentiation^39,40^ (Supplementary Fig. 6a). After 21 days of neuronal differentiation of vm-NPCs, iNeurons expressed essential markers of the dopaminergic fate, such as *TH* (tyrosine hydroxylase), *DAT* (dopamine transporter), *VMAT2* (vesicular monoamine transporter), and *NURR1* (Nuclear receptor-related 1 protein), as well as *GIRK2* (G-protein-regulated inward-rectifier potassium channel 2), typical marker of A9-like DAn of the substantia nigra, relevant to *VGLUT1* (Vesicular glutamate transporter 1; glutamatergic neurons) and *VGAT* (vesicular GABA transporter; GABAergic neurons), as evidenced by RT-qPCR (Supplementary Fig. 6b). At 28 days of neuronal differentiation, 72 ± 4.95% (mean ± standard error of mean; SEM) of microtubule-associated protein 2^+^ (MAP2^+^) healthy neurons (Hn) were TH^+^ by immunofluorescence, whereas PD neurons (PDn) comprised 52.40 ± 3.50% TH^+^ DAn (Supplementary Fig. 6c). We have previously characterized in detail the disease-associated phenotypes of PDn from p.A53T-αSyn iPSCs versus Hn^30,31,53^. In accordance with our past observations, the rabies virus-based assay revealed lower synaptic connectivity in PDn, compared to Hn (Supplementary Fig. 6d), and increased frequency and peak amplitude in Ca^2+^ activity (Supplementary Fig. 6e).

Subsequently, we established neuron-astrocyte co-cultures by combining Hn or PDn with Ha or PDa, at all possible arrangements (Fig. 3a). iNeurons at day 21 of neuronal differentiation were seeded onto replated day-28 iAstrocytes and homogeneous neuronal densities were verified across co-culture pairs by counting MAP2^+^ cells 12 h after seeding (Supplementary Fig. 7a). At later stages, significant differences in neuronal morphology were noted depending on whether iNeurons were co-cultured with Ha or PDa. At two days, a decrease in total neurite length and a concomitant increase in the number of primary neurites were observed when Hn were co-cultured on PDa, compared to Ha. Conversely, co-culture of PDn on Ha improved neuritic outgrowth and branching, compared to PDa (Fig. 3b). At the same time-point, an effect on neuronal viability and/or differentiation was also observed. Hn displayed similar densities whether cultured on Ha or PDa, however the number of PDn was significantly higher when cultured on Ha, as compared to PDa (Fig. 3c). This beneficial effect was more pronounced after 14 days in co-culture when the viability/differentiation of total MAP2^+^ and, to a greater extent, of TH^+^ PDn were considerably rescued by Ha. At this time-point, the adverse effects of PDa became prominent also in Hn, since both MAP2^+^ and TH^+^ vmDAn were reduced on PDa, compared to Ha. This indicates that PDa exert a neurotoxic effect even on Hn (Fig. 4a).

**Figure 3.**
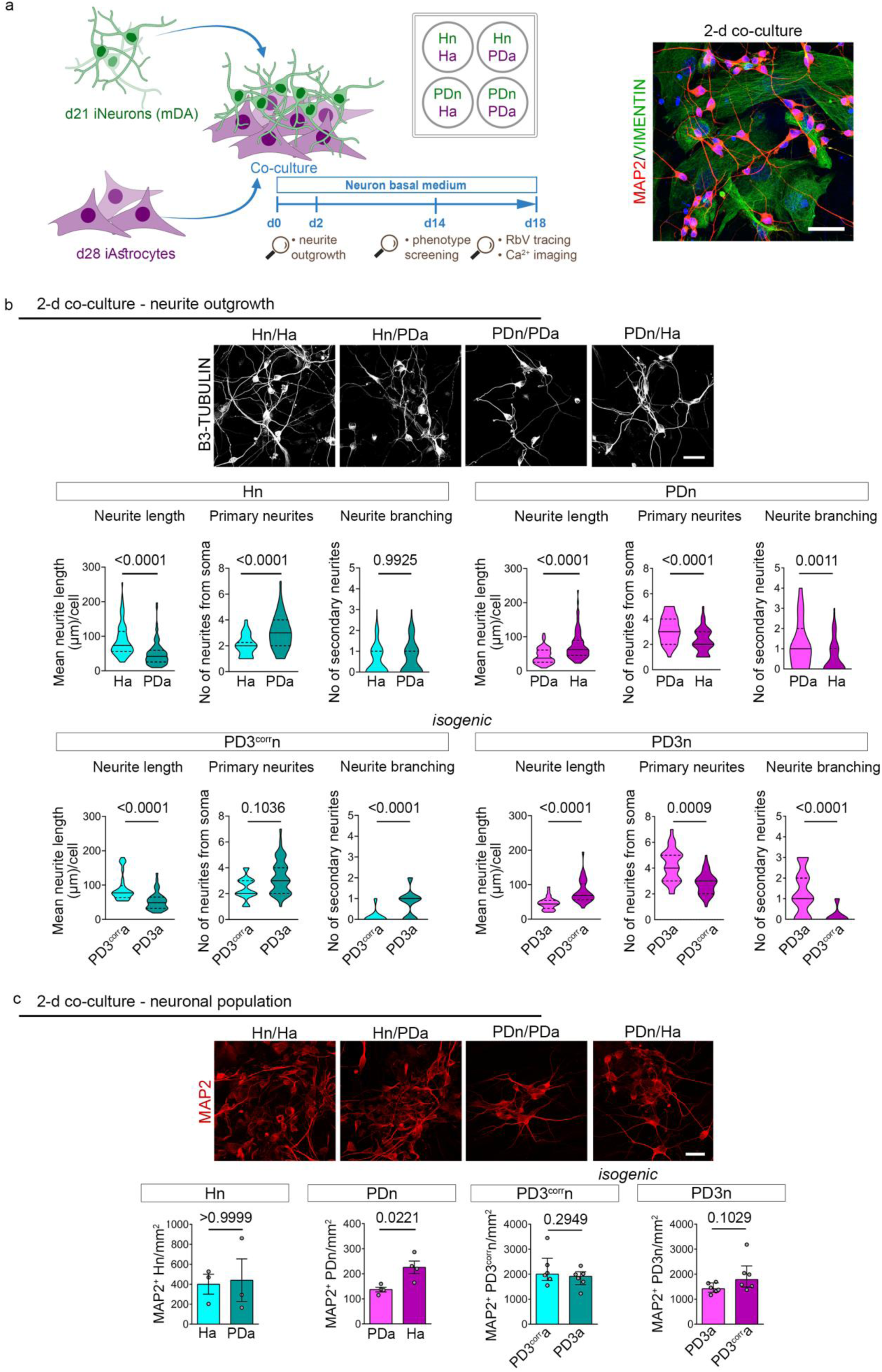
Effects of p.A53T-αSyn astrocytes on neurite outgrowth and neuronal numbers after 2 days in co-culture. **a** Schematic overview showing the co-culture setup of iPSC-derived vmDAns and iAstrocytes in all combinations and representative confocal image of a 2-day co-culture, immunostained for the neuronal marker MAP2, the astrocytic marker Vimentin and DAPI nuclear counterstain. Scale bar = 30 μm. **b** Representative confocal images of 2-day co-cultures of Hn, on Ha and PDa, and PDn, on PDa and Ha, showing neurite outgrowth after B3-tubulin immunostaining. Scale bar: 30 μm. Violin plots show the distributions of neurite length, primary neurites, and neurite branching across all combinations (non-isogenic, top; isogenic, bottom). **c** Representative confocal images of 2-day co-cultures showing MAP2^+^ populations of Hn and PDn with Ha and PDa. Scale bar: 30 μm. Quantification is provided in the accompanying bar plots (left, non-isogenic; right, isogenic). Data are presented as median with interquartile range (b) or mean ± SEM (c). (b) Mann-Whitney test was used for comparisons of neurite lengths; Kolmogorov Smirnov test was used to compare one-dimensional probability distributions for number of neurites and branching; (c) Ratio-paired t-test was used for comparisons in non-isogenic and unpaired t-test for isogenic experiments. Sample sizes: b, n = 98 neurons (Hn/Ha), 96 (Hn/PDa), 86 (PDn/PDa), 89 (PDn/Ha) from 3 Hn and 3 PDn non-isogenic lines; isogenic, n = 83 neurons (PD3^corr^n/PD3^corr^a), 78 (PD3^corr^n/PD3a), 86 (PD3n/PD3a), 89 (PD3n/PD3^corr^a); c, n = 3 Hn and n = 4 PDn non-isogenic lines; n = 83 FOV (PD3^corr^n/PD3^corr^a), 79 (PD3^corr^n/ PD3a), 29 (PD3n/PD3a), 36 (PD3n/PD3^corr^a). *Ha, healthy astrocytes; MAP2, Microtubule-associated protein 2; PDn, p.A53T-αSyn neurons; PDa, p.A53T-αSyn astrocytes; Ha, healthy astrocytes; Hn, healthy neurons; PD3a, p.A53T-αSyn astrocytes; PD3^corr^a, Corrected isogenic astrocytes; PD3n, p.A53T-*α*Syn neurons; PD3^corr^n, Corrected isogenic neurons; SEM, standard error of the mean, vmDAn, ventral midbrain dopaminergic neurons*.

**Figure 4.**
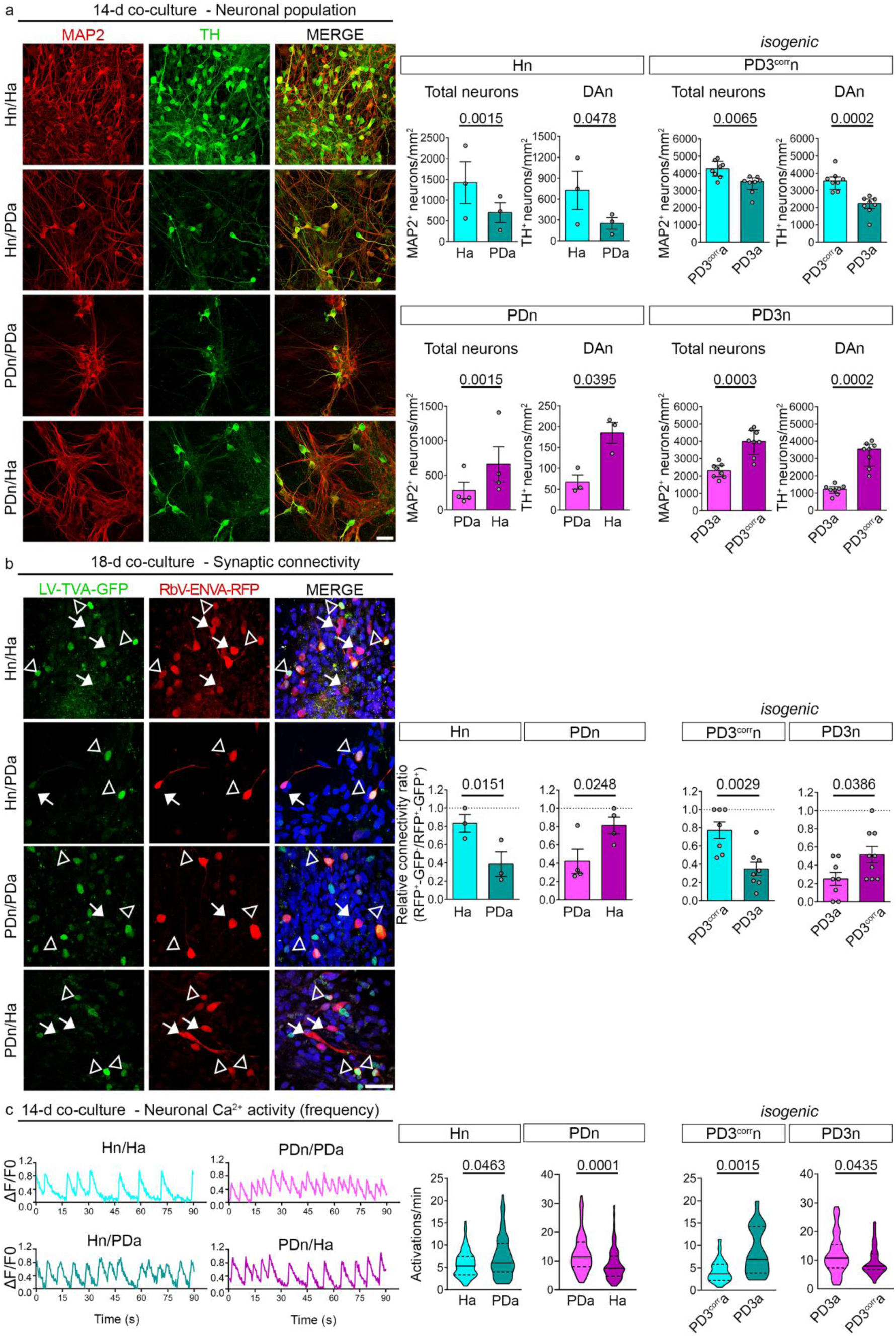
Impact of p.A53T-αSyn astrocytes on neuronal numbers, synaptic connectivity and neuronal calcium activity after 14 days in co-culture. **a** Representative confocal images of 14-day co-cultures of Hn on Ha or PDa, and of PDn on PDa or Ha, after immunostaining for MAP2 and TH. Scale bar: 30 μm. Accompanying bar plots show the quantification of total MAP2^+^ neurons and TH^+^ DAn in healthy and PD cultures across all combinations (left panel, non-isogenic; right panel, isogenic). **b** Representative confocal images of rabies virus-based retrograde monosynaptic tracing in 18-day co-cultures of Hn on Ha or PDa, and PDn on PDa or Ha. Target presynaptic neurons are RbV-ENVA-RFP^+^/LV-TVA-GFP^-^ (solid arrows) and starting postsynaptic neurons are RbV-ENVA-RFP^+^/LV-TVA-GFP^+^ (empty arrowheads). Scale bar, 30 µm. Bar plots represent the quantification of the relative connectivity ratio in Hn and PDn, across all combinations (left panel, non-isogenic; right panel, isogenic). **c** Representative Fluo4AM traces showing spontaneous Ca^2+^ activity in Hn co-cultured with Ha or PDa, and in PDn co-cultured with PDa or Ha, after 14 days. Violin plots show the distribution of Ca^2+^ oscillation frequencies in Hn and PDn, across all combinations normalized to their average basal response after background subtraction (left panel, non-isogenic; right panel, isogenic). Data are presented as mean ± SEM (a, b) or median with interquartile range (c). Ratio-paired two-tailed t-tests were used for comparisons in non-isogenic lines, unpaired t-test for isogenic in (**a**) and paired t-test for non-isogenic lines, unpaired t-test for isogenic in (**b**); Mann-Whitney test was used in (**c**). Sample sizes: a, b, n = 3 Hn and 4 PDn non-isogenic lines, from separate experiments; a, isogenic n = 8 FOV in each combination; b, isogenic n = 7 FOV (PD3^corr^n/PD3^corr^a); 8 FOV (PD3^corr^n/ PD3a); 8 FOV (PD3n/PD3a); 9 FOV (PD3n/PD3^corr^a) [Note: Samples Hn/Ha, Hn/PDa, PD3^corr^n/PD3^corr^a, and PD3^corr^n/ PD3a have been reused in Fig. 1c, in a different context]; c, n = 90 neurons (Hn/Ha); 93 (Hn/PDa); 80 (PDn/PDa), 93 (PDn/Ha), across 2 Hn and 3 PDn non-isogenic lines, in separate experiments; n = 24 neurons (PD3^corr^n/PD3^corr^a), 21 (PD3^corr^n/ PD3a), 50 (PD3n/PD3a); 43 (PD3n/PD3^corr^a). *DAn, dopaminergic neurons; Hn, healthy neurons; MAP2, Microtubule-associated protein 2; PDn, p.A53T-*α*Syn neurons; PDa, p.A53T-*α*Syn astrocytes; PD3a, p.A53T-*α*Syn astrocytes; PD3^corr^a, Corrected isogenic astrocytes; PD3n, p.A53T-*α*Syn neurons; PD3^corr^n, Corrected isogenic neurons; RbV, Rabies Virus; SEM, standard error of the mean; TH, tyrosine hydroxylase*.

Using the rabies virus-based retrograde monosynaptic tracing system as a proxy for structural network integrity, likely affecting functionality, we demonstrated that PDa had negative impact on the synaptic connectivity of Hn, compared to Ha. Moreover, the synaptic connectivity of PDn was improved when co-cultured with Ha (Fig. 4b). Further, in live Ca^2+^ imaging experiments, Hn displayed higher frequency on PDa, as compared to their counterparts on Ha, whereas PDn in the presence of Ha displayed a significant reversal (Fig. 4c). As Ca^2+^ dynamics regulate cell survival and/or differentiation, including neuritic growth, and synaptic connectivity with astrocytic input^54^, our data suggest that the observed alterations in Ca^2+^ signaling could explain, at least in part, the observed phenotypes.

Together, our data reveal that PD iAstrocytes impair neuronal integrity and function of both PD and healthy neurons, while healthy iAstrocytes have neuroprotective properties.

### PD-associated neuropathology including Lewy-like structures in neurons co-cultured with p.A53T-αSyn astrocytes is alleviated by control astrocytes

Inclusions of αSyn in neuronal perikarya, known as LB, as well as in neurites, referred to as LN, are typical in PD and collectively constitute Lewy-related αSyn pathologies^55,56^. By p(Ser129)αSyn immunostaining of PDn-PDa co-cultures at 14 d, we identified intracellular LB-like inclusions, as well as small-(< 4 μm) and large-(> 4 μm) diameter extra-cytoplasmic inclusions, and Lewy-like neurites (Fig. 5a, b), that corresponded to histopathological findings typically identified in post-mortem PD brains^57,58^. Additional neuropathological features, also seen in human brain histopathological analyses, included swollen axonal varicosities (both positive and negative for beta-3 tubulin), bulbous/dystrophic axonal endings, likely representing retracting axons, and neuronal cell bodies that were highly positive for p(Ser129)αSyn and presented a necrotic (condensed) nucleus, corresponding to degenerating neurons (Fig. 5a, c). The LB-like inclusions were positive for p62, ubiquitin, p(Thr212)Tau, αSyn aggregates as identified by the MJFR-14-6-4-2 antibody, and Thioflavin S (Fig. 5b), typical markers of LB^59^. Each of these pathological features was quantified in all co-culture combinations, demonstrating significantly different values between genotypes (Supplementary Fig. 8, 9). In particular, Hn performed worse in every aspect when cultured on PDa, while the pathological features of PDn were rescued on Ha. Subsequently, individual values were integrated in a relative readout, the relative PD-neuropathology index (Fig. 5d, see Methods). This approach demonstrated that PD-associated neuropathology was induced in Hn when co-cultured with PDa, as compared to Ha. Conversely, PD-related neuropathology of PDn was alleviated when co-cultured with Ha (Fig. 5c, d). To the best of our knowledge this is the first time that a systematic and comprehensive characterization of PD-related pathologies is performed in an iPSC-based human PD model comprising neurons and astrocytes, manifesting authentic-like histopathological hallmarks.

**Figure 5.**
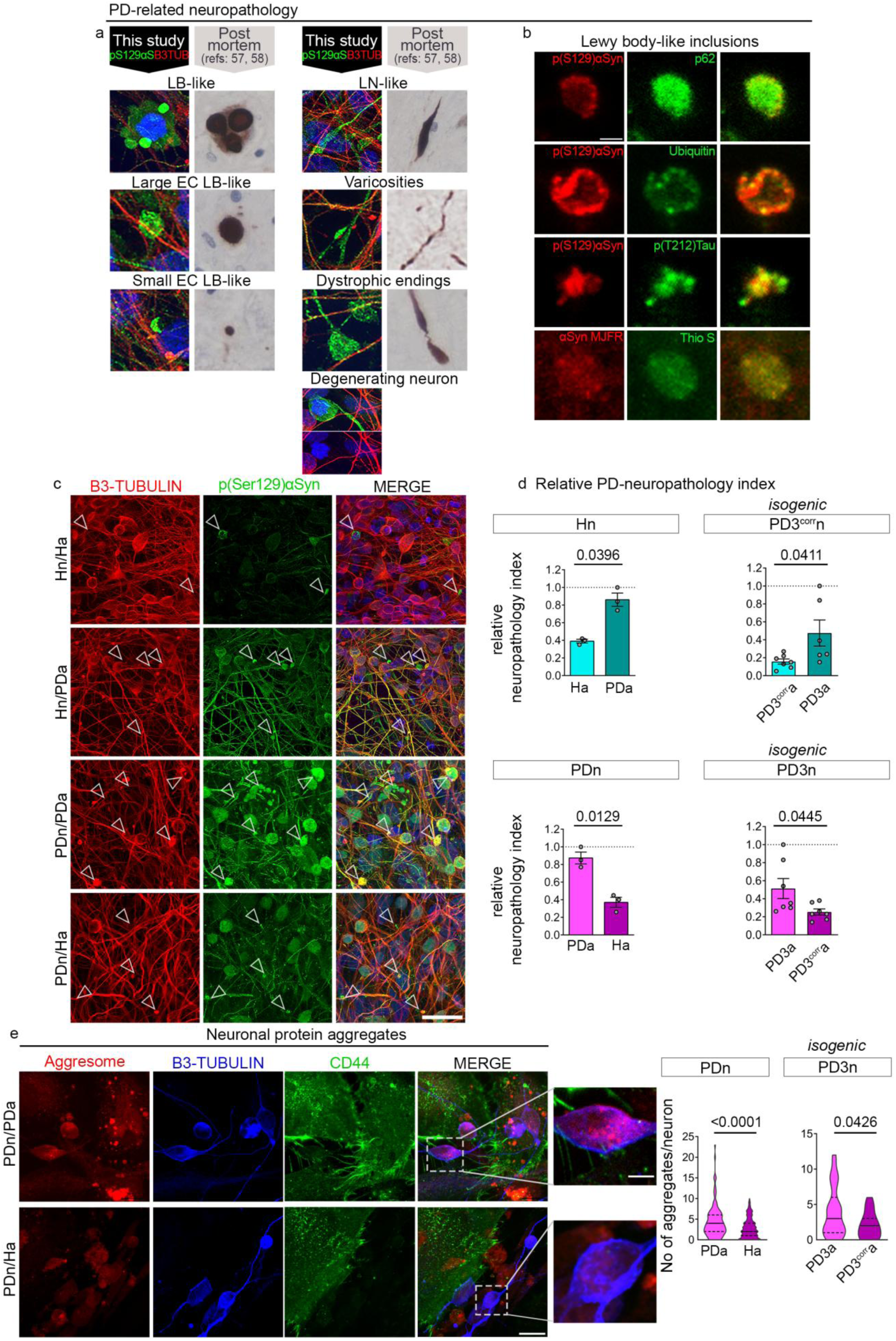
PD-relevant neuropathological features including Lewy-like pathology in p.A53T-αSyn iNeuron-iAstrocyte co-cultures. **a** Typical neuropathological hallmarks of PD identified by confocal analysis in PDn co-cultured with PDa, after immunostaining for pSer(129)αSyn and B3-tubulin; here benchmarked against relevant histopathological observations from post-mortem PD brains, as previously reported^57,58^ (permitted use). **b** Representative confocal images of LB-like inclusions detected in 14-day PDn/PDa co-cultures, double-stained for typical LB markers, in colocalization pairs: pSer(129)αSyn with p62, pSer(129)αSyn with ubiquitin, pSer(129)αSyn with pThr(212)Tau, and aggregated αSyn with Thioflavin S. Scale bar: 1 μm. **c** Representative confocal images of 14-day co-cultures of Hn on Ha or PDa, and PDn on PDa or Ha, immunostained for pSer(129)αSyn and B3-tubulin. Arrowheads indicate identified neuropathological features. Scale bar: 30 μm. **d** Bar plots showing the relative neuropathology index, calculated by integrating measurements of all individual neuropathological features in co-cultures of Hn or PDn, across all combinations (non-isogenic, left; isogenic, right). **e** Representative confocal images and magnified insets from 14-day co-cultures of PDn on PDa or Ha, showing Aggresome staining and immunostaining for the neuronal and astrocytic markers B3-tubulin and CD44, respectively. Scale bar, 30 μm and 5 μm for inset. The violin plots represent the distributions of the numbers of aggregates per neuron in PDn co-cultured on PDa or Ha (non-isogenic, left; isogenic, right). Data are presented as mean ± SEM (d) or median with interquartile range (e); Paired t-test was used for group comparisons of non-isogenic lines and unpaired t-test for isogenic in d and Mann-Whitney test was used in e; Sample sizes: d, n = 3 Hn and 3 PDn non-isogenic lines, from separate experiments; isogenic, n = 7 FOV (PD3^corr^n/PD3^corr^a); 6 (PD3^corr^n/ PD3a), 7 (PD3n/PD3a); 7 FOV (PD3n/PD3^corr^a); e, n = 137 (PDn/PDa) and 215 (PDn/Ha) neurons, across 3 PDn non-isogenic lines from separate experiments; n = 30 (PD3n/PD3^corr^a) and 29 (PD3n/PD3a) neurons. *Ha, healthy astrocytes; Hn, healthy neurons; LB, Lewy body; MAP2, Microtubule-associated protein 2; p(Ser129) αSyn, phosphorylated α-synuclein at Serine 129; PDn, p.A53T-αSyn neurons; PDa, p.A53T-αSyn astrocytes; PD3a, p.A53T-αSyn astrocytes; PD3^corr^a, Corrected isogenic astrocytes; PD3n, p.A53T-αSyn neurons; PD3^corr^n, Corrected isogenic neurons*.

Additionally, using the Proteostat aggresome detection kit, we detected a significant reduction in the number of protein aggregates in the perikarya of PDn when they were co-cultured for 14 days with Ha, as compared to being co-cultured with PDa (Fig. 5e).

Overall, these experiments document the role of PD astrocytes in exacerbating neurodegeneration and highlight the capacity of healthy astrocytes to rescue PD neurons, probably by supporting the clearance, or preventing the accumulation, of neuronal protein aggregates.

### Deficient endocytic clearance of exogenous αSyn cargo in p.A53T-αSyn astrocytes

We next explored if the poor capacity of p.A53T-αSyn mutant astrocytes to alleviate protein aggregates from co-cultured PD neurons, stems from their compromised ability to internalize and clear neuronal αSyn species^23,32,60^. We incubated Ha and PDa with recombinant αSyn pre-formed fibrils (αSyn-PFFs) conjugated with AlexaFluor568 and observed the kinetics of αSyn-PFF endocytosis at time intervals up to 16 h (Fig. 6a, b). During the first 2 h, no significant difference was detected between αSyn-PFFs internalized by Ha and PDa, whereas at 4 and 6 h, αSyn-PFFs started to accumulate in the cytoplasm of PDa, compared to Ha, and reached significance at 16 h. To uncouple cumulative uptake from degradation during the 16-h period, the same experiment was repeated, but this time excess αSyn-PFFs were washed away after 2 h. Again at 16h, significantly more αSyn-PFFs had accumulated in PDa compared to Ha, verifying the clearance lag (Fig. 6c).

**Figure 6.**
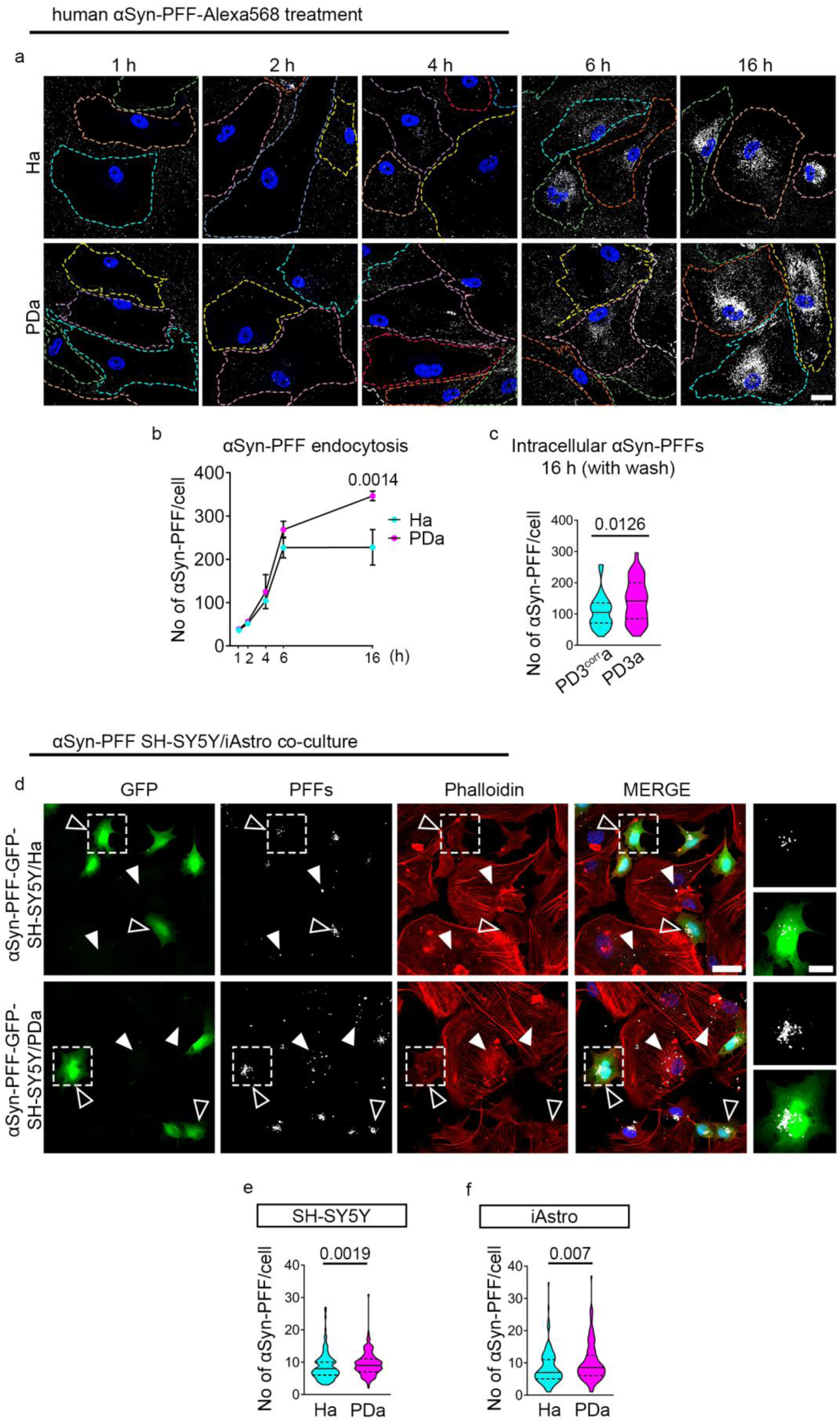
Impaired endocytosis of αSyn by p.A53T-αSyn Astrocytes. **a** Representative confocal images of Ha and PDa at 1, 2, 4, 6, and 16 h after treatment with αSyn-PFF-AlexaFluor568 (white). Randomly-colored dashed lines demarcate cell borders. Scale bar: 30 μm. **b** Line graph illustrating the time course of intracellular processing of uptaken αSyn-PFF-Alexa568 in Ha and PDa. **c** Quantification of αSyn-PFF-Alexa568 detected intracellularly in PD3^corr^a and PD3a at 16 h after a 2-h PFF treatment followed by washout. **d** Representative confocal images of Ha and PDa co-cultured for 24 h with GFP-SH-SY5Y human neuroblastoma cells loaded with αSyn-PFF-AlexaFluor568, and labeled with AlexaFluor647 Phalloidin. Magnifications of the boxed areas are shown in insets. Empty arrowheads indicate αSyn-PFFs in GFP-SH-SY5Y cells; solid arrowheads indicate αSyn-PFFs in iAstrocytes. Scale bar: 30 μm. **e** Violin plot showing the distribution of the numbers of αSyn-PFFs per cell in GFP-SH-SY5Y cells, after 24 h in co-culture with either Ha or PDa. **f** Violin plot showing the distribution of the numbers of αSyn-PFFs per cell in Ha and PDa after 24 h in co-culture with αSyn-PFF-loaded GFP-SH-SY5Y cells. Data are presented as mean ± SEM (b) or as median with interquartile range (c, e, f). Two-way ANOVA with Sidak post-hoc analysis for multiple comparisons was used in b, and Mann-Whitney tests were used for comparisons in c, e, f. Sample sizes: b, n = 3 Ha or PDa lines (including the isogenic pair) from separate experiments; c, n = 36 PD3^corr^a and 43 PD3a cells, from 3 wells in each case; e, n = 154 GFP-SH-SY5Y cells (co-cultured with 3 independent Ha lines) and 154 GFP-SH-SY5Y cells (co-cultured with 3 independent PDa lines) from separate experiments (including the isogenic pair); f, n = 145 cells (Ha) and 147 cells (PDa), across 3 Ha and 3 PDa lines (including the isogenic pair) from separate experiments. *αSyn-PFF, α-synuclein pre-formed fibrils; Ha, healthy astrocytes; PDa, p.A53T-αSyn astrocytes; PD3^corr^a, isogenic control; PD3a, p.A53T-αSyn astrocytes. Ha, healthy astrocytes; PDa, p.A53T-αSyn astrocytes; SEM, standard error of the mean*.

To further investigate the clearance capacity of iAstrocytes, GFP-tagged SH-SY5Y human neuroblastoma cells were first exposed to αSyn-PFFs for 16 h and, after washing, dissociation and mixing with either Ha or PDa, cells were chased for an additional 24 h (Fig. 6d). SH-SY5Y cells co-cultured with PDa retained significantly more intracellular αSyn-PFFs, compared to those co-cultured with Ha, confirming that Ha, but not PDa, can relieve neuronal αSyn aggregation (Fig. 6d, e). At the same time, PDa co-cultured with αSyn-PFF-loaded SH-SY5Y cells contained significantly more αSyn-PFFs, as compared to Ha, in line with their compromised clearance capacity (Fig. 6d, f). Altogether, these findings suggest that p.A53T-αSyn astrocytes have inherent deficiencies in endocytic protein aggregate degradation, making them less capable to relieve neuronal cells from accumulated αSyn.

### Proteomic analysis reveals aberrations in proteostasis in p.A53T-αSyn astrocytes

To gain mechanistic insight into the molecular pathways perturbed in p.A53T astrocytes, we performed comparative proteomic profiling of PDa and Ha, at steady state. Of 5,162 proteins represented across all samples (Supplementary Table 2), 454 proteins were differentially expressed (DE; p < 0.01); 148 up-regulated and 306 down-regulated in PDa (Supplementary Table 3). Among the most up-regulated proteins (Fig. 7a), were those participating in the inositol metabolic pathway and Ca^2+^ homeostasis (ITPKB, ITPR2, TPCN1), the mTORC1 pathway (CASTOR1), lipid transport and degradation (VLDLR, GALC), and extracellular vesicle secretion (CD82). The most down-regulated included proteins involved in phagocytosis (ANXA2, GULP1), autophagy (ANXA2, LRRC59, TBC1D2) and ubiquitination (UBE2L6), the neuroprotective astrocyte marker S100A10, and the TGF-β1 signal transducer SMAD3 (Fig. 7a).

**Figure 7.**
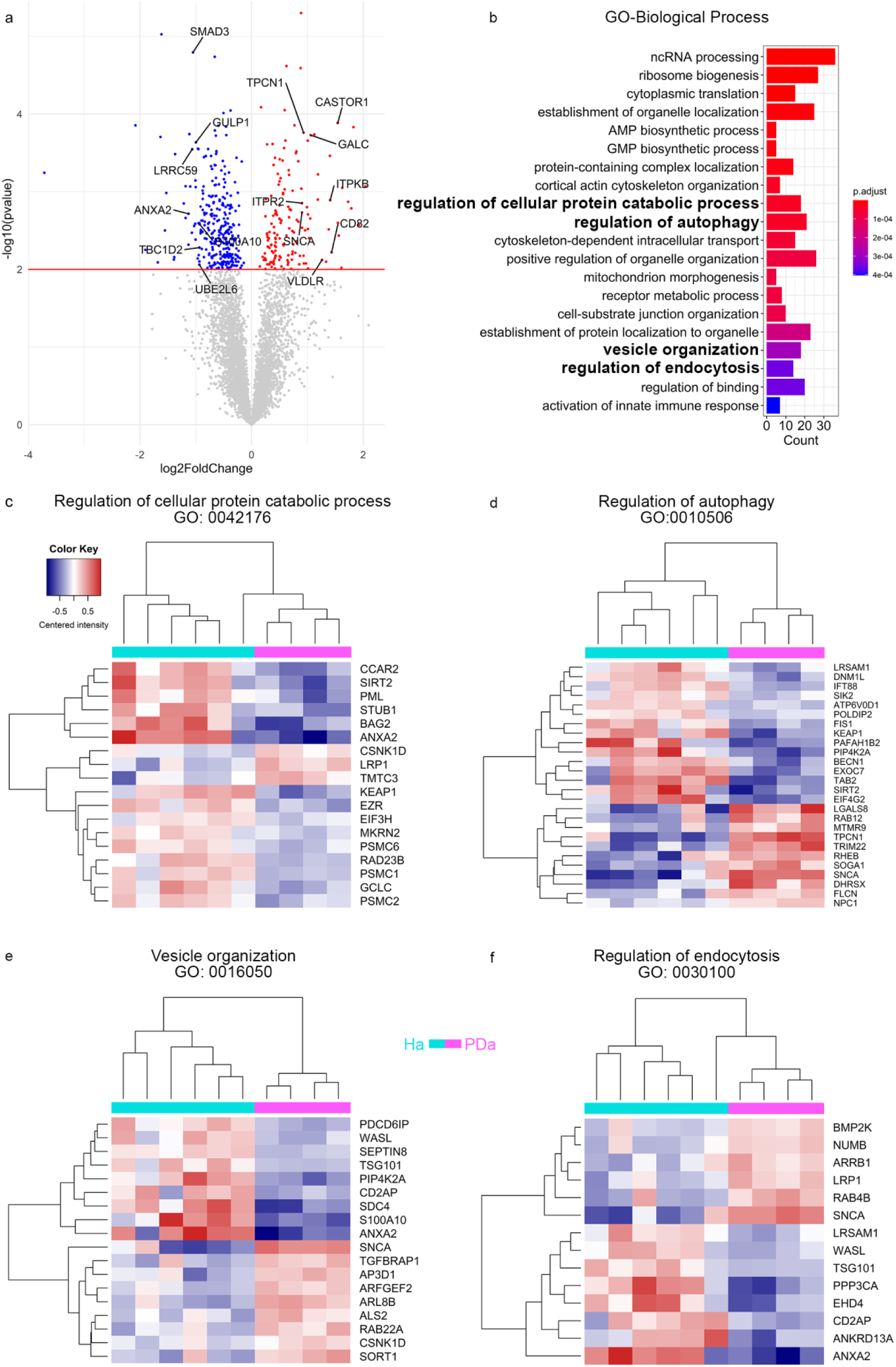
Proteome analysis of p.A53T-αSyn astrocytes - GOBP. **a** Volcano plot displaying -log10 p-values of proteins plotted against their corresponding log2 FC. Proteins are color-coded based on their DE: gray indicates no differential expression, red represents up-regulation in PDa and blue signifies down-regulation. Proteins of interest that are discussed in the text are named. **b** Bar plot showing the top 20 most significant GOBP terms. The length of each bar corresponds to the number of DE proteins associated with that term, while the color intensity reflects the term’s p-value. Terms that are further analyzed in heatmaps and discussed in the text are highlighted with bold fonts. **c-f** Heatmaps representing selected GOBP terms: ‘Regulation of cellular protein catabolic process’ **(c)**, ‘Regulation of autophagy’ **(d)**, ‘Vesicle organization’ **(e)**, and ‘Regulation of endocytosis’ **(f)**. Each heatmap shows the intensities of DE proteins (rows) across the samples (columns). For ‘Regulation of autophagy’ term, manually curated proteins from the list of DE proteins have been added to those identified in the term. The centered intensity (mean value across samples subtracted) for each protein is depicted. Hierarchical clustering has been applied to both rows and columns. *DE, differential expression; GOBP, Gene Ontology Biological Process; PDa, p.A53T-αSyn astrocytes; FC, fold change*.

Gene ontology (GO) enrichment analysis in the aspects Biological Process (GOBP), Molecular Function (GOMF), and Cellular Component (GOCC), highlighted dysregulated core pathways in p.A53T-αSyn astrocytes (Fig. 7 and 8). The top 20 most enriched GOBP terms comprised pathways related to proteostasis (Fig. 7b-f). Most of the proteins in *‘Regulation of cellular protein catabolic process’* were down-regulated in PDa, including several involved in the ubiquitin-proteasome system (UPS) (STUB1, TMTC3, KEAP1, MKRN2, PSMC1, PSMC2, PSMC6, RAD23B) (Fig. 7c). Of relevance, the GOMF term *‘Ubiquitin protein ligase activity’* was among the top 20 most enriched, and at large down-regulated (Fig. 8a, b). In *‘Regulation of autophagy’* (Fig. 7d), key down-regulated proteins included ATP6V0D1, PIP4K2A, and BECN1, while up-regulated proteins comprised RAB12, RHEB (activator of the mammalian target of rapamycin complex 1 [mTORC1]^61^), and FLCN, a GTPase-activating protein involved in the AMP-activated protein kinase (AMPK) and mTORC1 pathways^62^. Within *‘Vesicle organization’* (Fig. 7e) equally up- and down-regulated proteins included PDCD6IP, TSG101, SDC4, AP3D1, ARFGEF2, SORT1 and RAB22A, all involved in the regulation of vesicular trafficking process, endocytosis, multivesicular body biogenesis^63,64^ and Golgi vesicular transport^65–67^. Among the DE proteins in ‘*Regulation of endocytosis’* (Fig. 7f), LRP1 is a receptor involved in endocytosis and spreading of Tau^68^, PPP3CA is the catalytic subunit of calcineurin, and trafficking regulator EHD4 is an ATP- and membrane-binding protein that controls membrane reorganization/tubulation upon ATP hydrolysis^69^. *‘ATP hydrolysis activity’* came up among the most enriched GOMF terms, with proteins also involved in the UPS (Fig. 8a, c). Within GOCC (Fig. 8d-f), the enriched term *‘Early endosome’* (Fig. 8e) comprised non-redundant proteins such as WDFY2, FIG4, WASHC2C, SNX1, and PPP1R21 with roles in the processes of endosomal sorting, transport, and maturation^70–73^. Moreover, the protein set *‘Lysosomal membrane’* (Fig. 8f), comprised unique targets including ARL8B, as well as LAMTOR3 and LAMTOR2, both participating in the Ragulator, a lysosome-anchored protein complex involved in the mTORC1 pathway^74^, and MIOS, indirectly activating mTORC1 through inhibition of the GATOR1 subcomplex^75^. In all, the proteomic analysis revealed that the p.A53T mutation in astrocytes leads to altered levels of proteins implicated in inositol and lipid metabolism, Ca^2+^ homeostasis, and the mTORC1 pathway. Among pathways related to proteostasis, endocytosis, the UPS, and the autophagy-lysosome pathway (ALP) were significantly affected.

**Figure 8.**
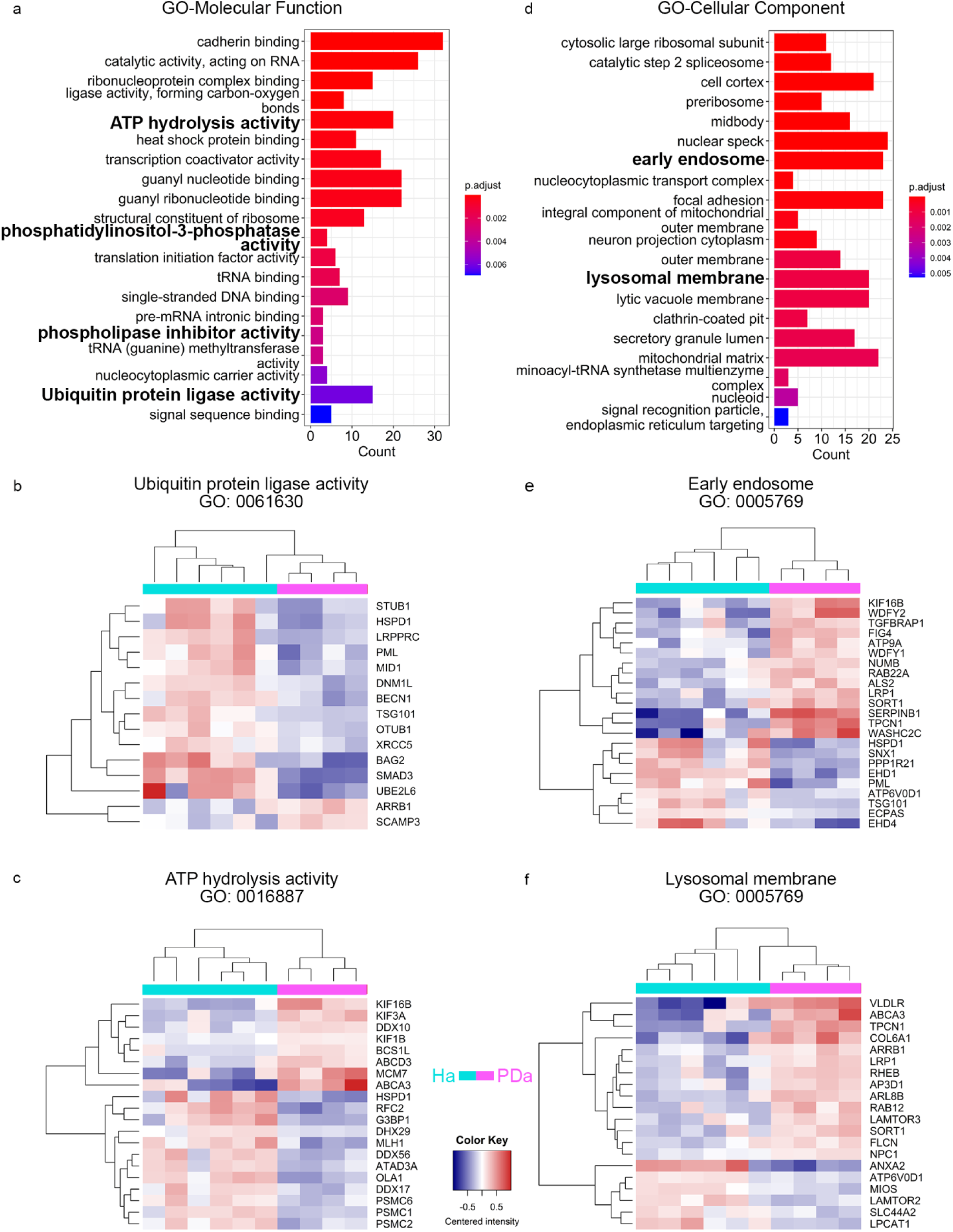
Proteome analysis of p.A53T-αSyn astrocytes (GOMF, GOCC). **a** Barplot showing the top 20 most significant GOMF terms. **b, c** Heatmaps for two selected terms: ‘ATP hydrolysis activity’ **(b)** and ‘Ubiquitin protein ligase activity’ **(c)**. **d** Bar plot showing the top 20 most significant GOCC terms. **e, f** Heatmaps for two selected terms: ‘Early endosome’ **(e)** and ‘Lysosomal membrane’ **(f)**. Heatmaps display the intensities of the DE proteins belonging to the term (rows) across samples (columns). In the bar plots, the length of each bar corresponds to the number of DE proteins belonging to that term, and its color is based on the p-value. The terms discussed in the text and analyzed in heatmaps are marked in bold. In the heatmaps, for each protein, the centered intensity (with its mean value subtracted) is shown. Hierarchical clustering was applied to both rows and columns. *GOMF, Gene Ontology Molecular Function; DE, differentially expressed; GOCC, Gene Ontology Cellular Component*.

### p.A53T-αSyn iAstrocytes display disturbances at the major protein clearance pathways

The UPS and the ALP are the two most important mechanisms for intracellular protein quality control. Failure of each of the two systems to degrade misfolded or aggregated proteins has been correlated with PD. In essence, αSyn is degraded by both the UPS and ALP, but it can also inhibit their function under pathological conditions^76^. Considering the accumulation of p(Ser129)αSyn and protein aggregation observed in PDa, and based on the proteomic data, we first examined the chymotrypsin-like proteasome activity in iAstrocytes. In agreement with the noted down-regulation of critical relevant proteins (STUB1, TMTC3, KEAP1, MKRN2, PSMC1, PSMC2, PSMC6, RAD23B), we found that the proteasome activity is significantly reduced in PDa (Fig. 9a), accompanied by accumulation of high-MW ubiquitinated proteins, as evidenced by Western Blot (Fig. 9b).

**Figure 9.**
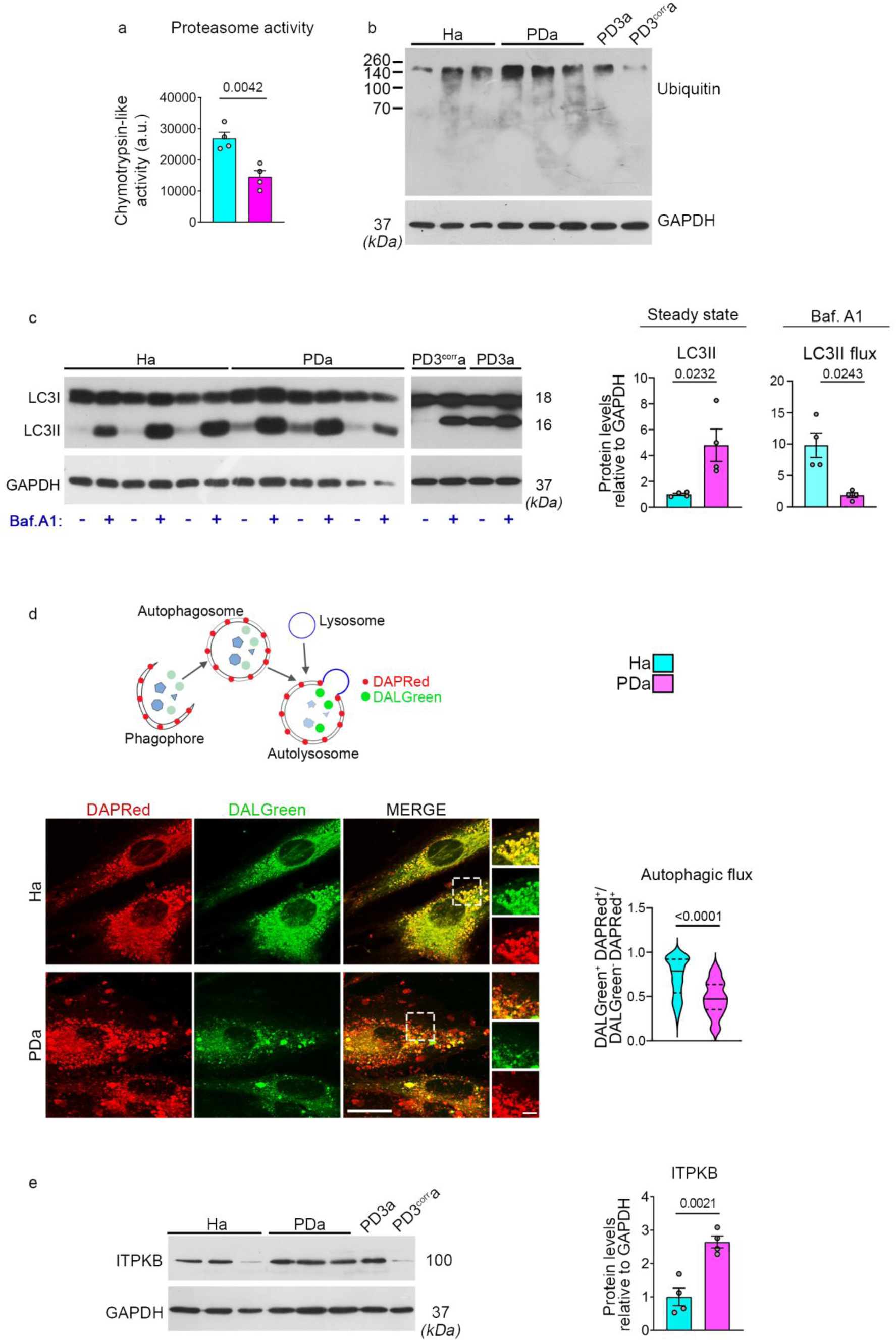
p.A53T-αSyn astrocytes present defects in the proteasome and autophagy protein quality control mechanisms. **a** Bar plot representing the quantification of chymotrypsin-like proteasomal activity in Ha and PDa. **b** Western blot analysis of ubiquitinated proteins, and GAPDH (loading control) in 3 Ha and 3 non-isogenic PDa lines, and in PD3a and PD3^corr^a isogenic lines. **c** Western blot analysis of LC3B (LC3I and LC3II) and GAPDH in 3 Ha and 3 PDa non-isogenic lines, and in PD3^corr^a and PD3a isogenic lines, before and after treatment with Bafilomycin A1. The bar plots represent quantification of steady-state LC3II levels normalized to GAPDH, and the LC3II flux (difference between Baf.A1-treated and untreated samples, relative to untreated and GAPDH); **d** (Top) Schematic of the DAPRed and DALGreen-based fluorescent detection system for monitoring the autophagic flux. (Bottom left) Representative confocal images of Ha and PDa after live incubation with DAPRed and DALGreen and insets with magnifications of the boxed areas. Scale bar = 30 μm. (Bottom right) Violin plot showing the quantification of the ratio (per cell) of DALGreen^+^-DAPRed^+^ autolysosomes to DALGreen^-^-DAPRed^+^ autophagosomes. **e** Western blot analysis of ITPKB and GAPDH (loading control) and quantification in 3 Ha and 3 PDa non-isogenic lines, and in PD3a, and PD3^corr^a isogenic lines. Data are presented as mean ± SEM (a, c, e). Unpaired two-tailed t-test was used for comparisons in a, c, e; Mann-Whitney was used in d. Sample sizes: a, c, e, n = 4 Ha and 4 PDa lines (including the isogenic pair); d, n = 91 Ha and 74 PDa across 3 Ha and 3 PDa non-isogenic lines, respectively. *Baf. A1, Bafilomycin A1; Ha, healthy astrocytes; LC3, microtubule-associated protein 1 light chain 3; Ha, healthy astrocytes; PDa, p.A53T-αSyn astrocytes; PD3^corr^a, corrected isogenic astrocytes; PD3a, p.A53T-αSyn astrocytes. Ha, healthy astrocytes; SEM, standard error of the mean*.

We next investigated the autophagic capacity of PDa, recognizing ALP as a primary degradation mechanism for aggregated αSyn^77^. Western blot showed that at steady state, the levels of LC3-II, the lipidated form of the autophagy facilitator LC3 (microtubule-associated protein 1A/1B-light chain 3) that is associated with the autophagosomal membrane, were significantly increased in PDa (Fig. 9c). We then asked if the elevated levels of LC3II in PDa reflect accumulating autophagosomes due to reduced autophagic flux. Bafilomycin A1 (Baf.A1) blocks the autophagic flux by inhibiting autophagosome-lysosome fusion, resulting in accumulation of LC3II^+^ autophagosomes^78^. After 4 h of Baf.A1 treatment, LC3II flux (calculated as the normalized difference between Baf.A1-treated and untreated samples) was significantly lower in PDa, as compared to Ha (Fig. 9c), indicating reduced autophagic flux.

We further employed DALGreen and DAPRed sensors, specialized fluorescent probes for monitoring the autophagic flux by selectively labeling autophagosomes and autolysosomes. DALGreen fluorescence, enhanced at low pH, traces autolysosomes while DAPRed labels both autophagosomes and autolysosomes^79,80^ (Fig. 9d). After 6 h of simultaneous DALGreen and DAPRed treatment, the ratio of autolysosomes (DALGreen^+^/DAPRed^+^) over autophagosomes (DALGreen^-^/DAPRed^+^) was significantly lower in PDa compared to Ha, confirming a hampered autophagic flux and disruption of the autophagosome-lysosome fusion process (Fig. 9d).

The observed Ca^2+^ dyshomeostasis in PDa as depicted by live cell imaging and in the proteomic analysis, could be linked to impaired autophagy. Hence, we examined the levels of ITPKB that was among the most up-regulated targets in proteomics. ITPKB is a lipid kinase that modulates Ca^2+^ dynamics and autophagy, while a positive correlation between αSyn (wild type, A30P, and A53T) and increased ITPKB levels were detected in post-mortem cortices from PD patients and in αSyn overexpression cellular systems^81^. By Western blot, we verified that ITPKB is significantly increased in PDa, relative to Ha (Fig. 9e).

We conclude that the p.A53T-αSyn mutation in astrocytes hinders crucial protein quality control mechanisms, including the proteasome system and autophagy, correlating with their observed contribution to PD pathology.

### The p.A53T-αSyn mutation affects lysosomal physiology in astrocytes

The proteomic data implicating lysosomal proteins, along with the observed phenotypes, led us to examine lysosomal integrity in PDa. LAMP1 (Lysosomal-associated membrane protein 1), is a transmembrane glycoprotein located at the lysosomal membrane with crucial roles in maintaining lysosomal integrity and function^82^. Western blot demonstrated that the levels of LAMP1 were significantly reduced in PDa, as compared to Ha (Fig. 10a). On the other hand, using Lysotracker DR labeling in live cells we detected increased number of lysosomes (Fig. 10b) that may reflect reduced consumption of lysosomes due to impaired autophagy or a compensatory response to lysosomal dysfunction^83^. Of relevance, the ATP6V0D1 subunit of V-type H^+^-ATPase on the lysosomal membrane, was down-regulated in PDa, as noted in the proteomics, and loss of V-ATPase subunits has been shown to cause accumulation of non-functional lysosomes and a block in the autophagic flux^84^. For the lysosomal enzymes to function properly the acidification of lysosomal lumen is tightly regulated by the V-type H^+^-ATPase and ion channels on the organelle membrane. Notably, defective lysosomal acidification has been implicated as a key driving factor in the pathogenesis of neurodegenerative diseases, including AD and PD^85^. Using the pH-sensitive fluorescent dye Lysosensor, we confirmed the significant decrease of lysosomal acidity in PDa (Fig. 10c) also verified in the lower Lysotracker DR fluorescence intensity in PDa (Fig. 10b). Subsequently, assessment of the pH-dependent activity of the lysosomal enzyme Cathepsin B, using the Magic Red kit, revealed a significant reduction in PDa as compared to Ha (Fig. 10d). Moreover, by LAMP2 immunofluorescence we detected significantly altered lysosomal positioning; in Ha, lysosomes were largely restricted to the perinuclear area while in PDa they appeared dispersed within the cytoplasm (Fig. 10e, also visible by Lysotracker DR staining in Fig. 10b). Proper lysosomal positioning is critical for lysosomal functions and regarding autophagy, perinuclear lysosomal positioning is important for autophagosome-lysosome fusion^86,87^. In relevance, lysosomal positioning regulates mTORC1 signaling, which in turn influences autophagosome formation^88^. As discussed above and evidenced by STRING analysis of the proteomic data, a number of mTORC1-related differentially expressed proteins, including CASTOR, RHEB, PDE6D, FLCN, LAMTOR3, LAMTOR2, MIOS, and ATP6V0D1, form a network component significantly perturbed in PDa (Fig. 10f). mTOR regulates autophagy through the direct phosphorylation of Unc-51-like autophagy activating kinase 1 ULK1. High mTOR activity prevents ULK1 activation by phosphorylation at Ser757, due to disruption of ULK1 and AMPK interaction which promotes autophagy^89^. In accordance, the ratio of p(Ser757)ULK1 over total ULK1 was significantly increased in PDa when compared to Ha, as assessed by Western blot (Fig. 10g). The above results demonstrate that defective lysosomal positioning and function in PDa result in dysfunctional autophagy that causes accumulation of protein aggregates, at least via disrupted mTORC1 signaling.

**Figure 10.**
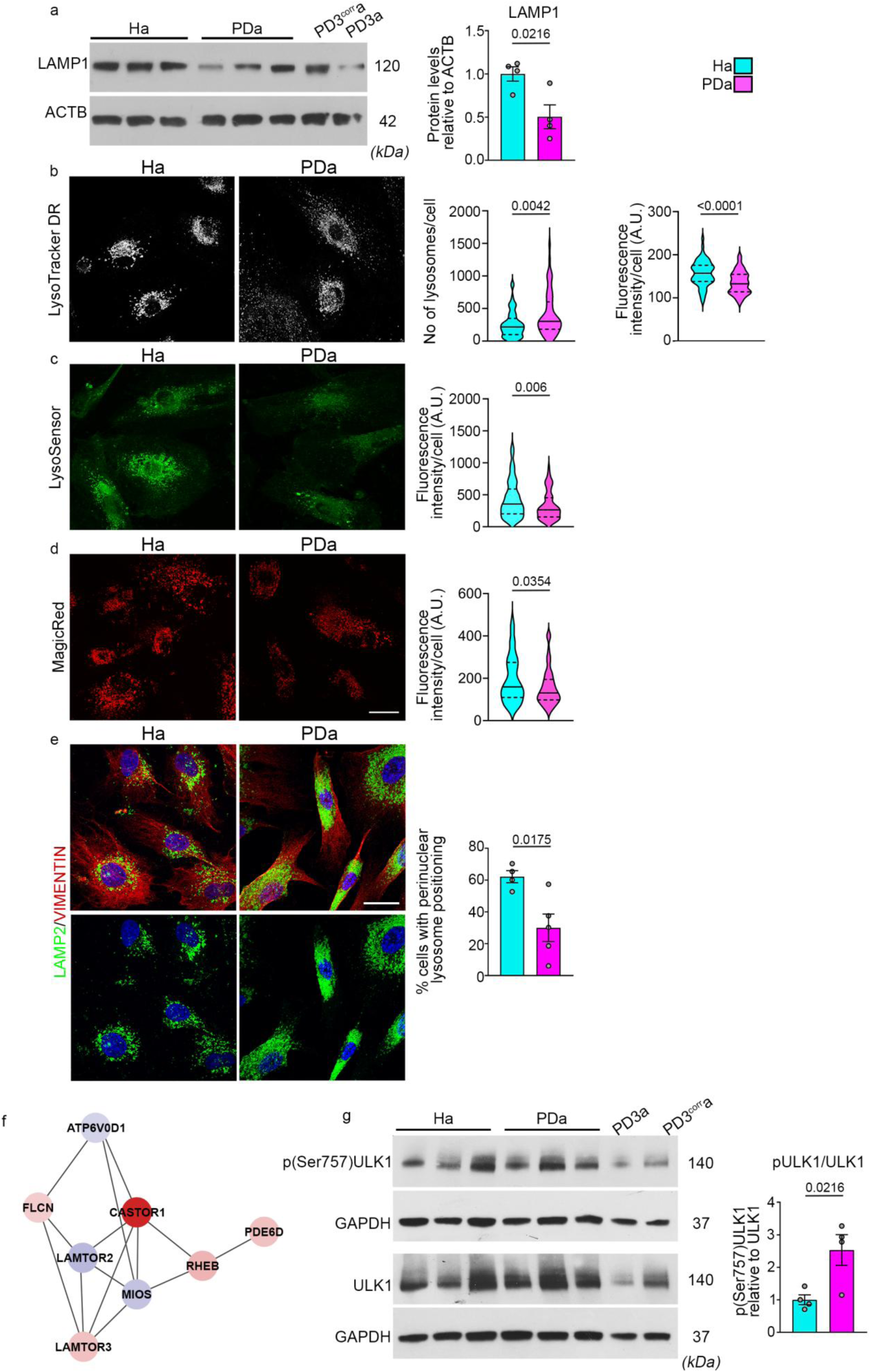
Disturbed lysosomes in iAstrocytes with the p.A53T-αSyn mutation. **a** Western blot analysis of LAMP1 and beta-actin (ACTB, loading control) in 3 Ha and 3 PDa non-isogenic lines, and the isogenic lines PD3^corr^a, and PD3a. The bar plot quantifies LAMP1 levels normalized to ACTB. **b** Representative images and quantification of live Ha and PDa after labeling with LysoTracker DR. Violin plots show the distributions of lysosome numbers per cell (left) and of the fluorescent intensities per cell (right). **c, d** Representative images and quantification of live Ha and PDa after labeling with the pH-sensitive fluorescent dye LysoSensor (c) or Magic Red indicating Cathepsin B activity (d); violin plots show the distribution of fluorescence intensities per cell. **e** Representative confocal images of Ha and PDa, after immunostaining for LAMP2 and Vimentin with DAPI. Bar plot shows the percentage of cells with perinuclear lysosomes relative to total cells. Scale bars b, c, d, in d, and e: 30 μm. **f** STRING network analysis visualizing interactions among DE proteins identified in PDa compared to Ha. Proteins up-regulated in PDa are highlighted in red, while down-regulated proteins are in blue. Connecting lines represent protein-protein associations. **g** Western blot analysis of ULK1, its phosphorylated form p(Ser757)ULK1, and GAPDH (loading control) in 3 Ha, 3 PDa non-isogenic lines, and PD3a and PD3^corr^a. The bar plot represents the levels of p(Ser757)ULK1 relative to ULK1. Data are presented as mean ± SEM in a, c-g, or as median and interquartile range in violin plots in b, c, d. Unpaired two-tailed t-tests were used for comparison in a, and e-g; Mann-Whitney test was used for the comparisons in b, c, d. Sample sizes: a and g, n = 4 Ha and 4 PDa lines; b, n = 60 cells across 3 Ha lines and 60 cells across 3 PDa lines; c, n = 112 cells across 3 Ha and 105 cells across 3 PDa lines; d, n = 74 cells across 3 Ha and 81 cells across 3 PDa lines; e, n = 4 Ha and 5 PDa lines (the isogenic pair is included in a-e, and g). *ACTB, Beta-actin; LAMP1, Lysosome-associated membrane protein 1; mTORC1, mammalian Target of Rapamycin Complex 1; Ha, healthy astrocytes; PDa, p.A53T-αSyn astrocytes; PD3^corr^a, corrected isogenic astrocytes; PD3a, p.A53T-αSyn astrocytes; SEM, standard error of the mean; ULK1, Unc-51-like kinase 1*.

## Discussion

By leveraging PD patient-derived iPSCs, this study demonstrates that the p.A53T-αSyn mutation in astrocytes induces critical cell-intrinsic defects associated with proteostasis, which in turn aggravate neuronal pathology. Proteomic profiling and functional analyses of PDa versus Ha revealed major affected pathways, involving the UPS, autophagy, endocytosis, and lysosomal function. Our data deepen and extend the role of astrocytes as essential regulators of neuronal physiology and document a non-cell autonomous contribution to PD neuropathology. In a co-culture system of human-derived iNeurons and iAstrocytes that reproduced major PD histopathological hallmarks, including Lewy-like pathology, our analysis captured exacerbation of PD-related neuronal phenotypes by PDa, and a considerable rescue by Ha. Moreover, PDa proved less competent than Ha in clearing aggregated αSyn, which resulted in accumulation of toxic αSyn species both in PDa and PDn, aggravating the neurodegenerative phenotype.

Human genetic iPSC-derived models have so far strived to faithfully replicate Lewy-related pathologies, without exogenous αSyn overexpression or the addition of preformed αSyn fibrils^90–92^. In the PDn-PDa co-cultures, we detected inherent, authentic-like PD neuropathology comprising intraneuronal and extracellular LB-like formations, positive for p(Ser129)αSyn, and typical LB constituents such as p62, ubiquitin, and p(Thr212)Tau, as well as LN-like structures and other features, reflecting human post-mortem PD brain neuropathological hallmarks, substantially resolved by Ha. This validates our setup as a reliable model for mechanistic studies, and a valuable platform for drug discovery displaying disease-relevant readouts. Further, the beneficial effects of Ha identified herein, support their potential therapeutic value, as previously alluded^38,93^, while the adverse effects of PDa on neuronal morphology and function indicate prospective consequences on disease progression. The observed pathological phenotypes may be attributed to the reduced phagocytic clearance capacity of PDa, as shown by their impaired ability to uptake and degrade toxic αSyn species. However, on present evidence we cannot rule out an additional paracrine astrocyte-to-neuron effect, either neurotoxic from PDa or neuroprotective from Ha^38,93^ that requires further investigation. Our initial observations suggest such a paracrine effect of iAstrocyte-conditioned media on iNeurons (unpublished data).

Astrocytic αSyn has been mainly considered of neuronal origin^5,15^. Here we show by immunofluorescence, Western blot and proteomics that αSyn is detected in astrocytes, in the absence of neurons. In rodent and immortalized astrocytes, overexpression of p.A53T-αSyn causes inhibition of autophagy, mitochondrial changes, ER stress, and apoptosis^94,95^, as well as lysosomal dysfunction^96^. Astrocyte-specific expression of p.A53T-αSyn in a transgenic mouse led to αSyn accumulation, blood brain barrier disruption, and neuroinflammation, ultimately resulting in neurodegeneration^97^, while transgenic mice with global p.A53T-αSyn expression displayed altered astrocytic Ca^2+^ activity^98^. In our human-relevant patient-derived genetic model, PDa displayed altered Ca^2+^ dynamics, and accumulation of intracellular αSyn species and protein aggregates. In line, intrinsic PDa defects were revealed in proteasome activity and autophagic function, consistent with the proteome analysis data.

As previously shown primarily in neuronal systems, under normal conditions αSyn is degraded through the UPS and chaperone-mediated autophagy (CMA)^99^. ALP is recruited when the UPS cannot resolve an overload of cellular αSyn, or is activated to compensate CMA impairment^100^. PD-linked genes, carrying causal mutations (*LRRK2*, *ATP13A2*) or non-causal, risk-associated variants (*GBA1*) have been implicated in endo-lysosomal processes in neuronal cells^77^ and more recently in astrocytes^14,21,29,46,101^. Moreover, previous studies have shown that αSyn aggregates inhibit the ALP by a direct effect on lysosomal components^99,102–104^, or by inhibition of vesicular trafficking^27,105–107^. Additionally, the αSyn pathogenic mutations p.A53T and p.G51D directly impair its lysosomal degradation and increase cellular protein concentration by extending the protein’s half-life^108^. Evidently, accumulation of aggregated αSyn is related with impairment of clearance mechanisms^109,110^.

Second to microglia, astrocytes are proficient scavengers of αSyn species released by neurons in the extracellular space, efficiently degrading αSyn^18,20,21,111,112^. However, upon extensive αSyn internalization, overcharge of the endo-lysosomal pathway results to large intracellular deposits in astrocytes, suggesting incomplete digestion^113^, indicative of impaired phagocytic clearance^114^. In this context, among the most down-regulated proteins in PDa, according to the proteome analysis, was ANXA2, a Ca^2+^-regulated protein that contributes to phagocytosis^115^, endosome stabilization^116^, autophagy^117,118^, and astrocytic clearance of extracellular αSyn aggregates. Loss of ANXA2 function leads to reduced αSyn internalization in astrocytes^119^. GULP1, required for efficient astrocytic phagocytosis^120^ was also among the most down-regulated proteins. Accordingly, PDa displayed impaired phagocytic clearance.

Our observations point to dysregulation of the lysosomal hub in the less-acknowledged astrocytic component, aligning with previous studies that suggest a major role of lysosomal disruption in PD pathogenesis^109^. In support, our data revealed disturbances in anatomical and functional lysosomal features, while the proteomic analysis highlighted differential expression of proteins implicated in modulation of lysosomal function. Down-regulated targets included ATP6V0D1, subunit d1 of V-type H^+^-ATPase in the integral membrane V0 complex, responsible for acidifying intracellular compartments including the lysosomes^121^ and target of mTORC1^122^, as well as PIP4K2A required for autophagosome-lysosome fusion^123^. Up-regulated targets comprised GALC, galactosylceraminidase, a lysosomal enzyme involved in glycosphingolipid degradation with pathogenic gene variants linked with Krabbe disease, an infantile neurodegenerative lysosomal storage disorder. Meta-analyses of genome-wide association studies (GWAS) have identified a correlation of *GALC* locus variants with PD, driven by increased galactosylceraminidase expression and activity^124^, with enrichment in glial cells^125^. RAB12, a Ras-related small GTPase found in vesicular compartments related to endosome function with critical involvement in PD^126^, reported as regulator of LRRK2 activity in response to lysosomal damage^127^, and ARL8B, a small GTPase regulating lysosome positioning^128^, also modulating late endosome-lysosome fusion^129^, were also found up-regulated. Given that lysosome positioning is a determinant of intralumenal pH, ARL8B plays a critical role in lysosomal function^130^. The mTORC1 pathway that regulates autophagy via phosphorylation of ULK1 at Ser757, having an inhibitory role^89^, is also modulated by lysosomal integrity. STRING analysis revealed an mTORC1-related hub, mostly up-regulated, while we also confirmed increased levels of p(Ser757)ULK1 in PDa. These results provide evidence that lysosomal dysfunction in PDa is linked to mTORC1 dysregulation, which may be responsible, at least in part, for the observed disturbances in autophagy.

Our data further implicate Ca^2+^ dyshomeostasis and inositol metabolism in lysosomal dysfunction, based on (i) the abnormal Ca^2+^ intracellular activity recorded in PDa, as compared to Ha, (ii) a considerable number of relevant DE proteins in proteomics and (iii) enrichment of the GOMF terms *‘Phosphatidylinositol-3-phosphatase activity’* and *‘Phospholipase inhibitor activity’*. Among the most up-regulated proteins, the two-pore Ca^2+^ channel TPCN1, critically participating in regulation of Ca^2+^ within endosomes and lysosomes, is abundantly expressed in astrocytes, and involved in intracellular Ca^2+^ signaling, endo-lysosomal trafficking, lysosomal function, regulation of mTORC1 activity^131^, and autophagy, while its dysregulation has been implicated in neurodegenerative diseases and lysosomal storage disorders^131,132^. Other up-regulated proteins included: ITPR2 that mediates Ca^2+^ release from the ER and is the predominant inositol-trisphosphate (IP3) receptor in astrocytes, while its ablation strongly attenuates astroglial Ca^2+^ signals^133–135^. ITPKB, verified as up-regulated in PDa, was previously found increased in PD^81^, while GWAS studies have linked its gene variants with sporadic PD^136^. Interestingly, ITPKB has been shown to modulate αSyn aggregation by inhibiting ER-to-mitochondria Ca^2+^ transport and affecting autophagy^137,138^. Among down-regulated proteins, PPP3CA is the catalytic subunit alpha of calcineurin, a phosphatase controlled by lysosomal Ca^2+^ release to dephosphorylate transcription factor EB (TFEB), master regulator of lysosomal and autophagic function^139^, and promote its nuclear translocation. Down-regulated BECN1 is key regulator of autophagy initiation, participating in the formation of the PIP3KC3 complex that generates PI3P, recruiting proteins essential for autophagy progression; PIP4K2A^123^ and FIG4^140^, that participates in the PIKfyeve complex, mediating synthesis and turnover of lysosomal PI(3,5)P, are both indispensable for the fusion between autophagosomes and lysosomes.

In conclusion, the p.A53T-αSyn mutation drives astrocytic pathology and leads to malfunctioning of cellular clearance mechanisms, rendering PD astrocytes incompetent to maintain their own proteostasis, support neuronal fitness, or rescue PD neurons. For the first time in a patient-derived cellular system of neurons and astrocytes, exhibiting Lewy-related pathology, it is demonstrated that healthy astrocytes can mitigate neurodegeneration, providing insights for therapeutic manipulation. Conversely, mutant PD astrocytes emerge as critical contributors to PD neuropathology, while the pathways impaired by p.A53T-αSyn converge on dysfunctions of the astrocytic lysosomal hub, and suggest novel mechanistic targets.

## Methods

### Ethics statement

This study involves previously characterized induced pluripotent stem cell (iPSC) lines^25,30^ derived in accordance with established ethical guidelines and distributed under Material Transfer Agreements to ensure compliance with all legal and ethical standards.

### Culture of human induced pluripotent stem cells (iPSCs)

The iPSCs used included lines from two male PD patients (PD1 and PD2) carrying the G209A (p.A53T) SNCA mutation and one age- and sex-matched non-PD subject (H1) (two different clones from each subject), previously generated and characterized in our lab^30^; an age- and sex-matched non-PD control line (H2) obtained from the New York Stem Cell Foundation; a female PD iPSC line (PD3) carrying the G209A SNCA mutation and its gene-corrected isogenic control (PD3^corr^) provided by Dr Rudolf Jaenisch^25^ (Supplementary Table 1). iPSCs were cultured either on mouse embryonic fibroblasts (Thermo Fisher Scientific, Waltham, MA, USA) or under feeder-free conditions on Matrigel (BD Biosciences, Franklin Lakes, NJ, USA), following standard procedures. iPSCs were quality checked according to the ISSCR standards^141^ for karyotype integrity, mycoplasma contamination, undifferentiated state and pluripotency, and presence of the disease mutation. Unless otherwise stated, differentiations from independent non-isogenic iPSC clones were considered biological replicates (H1.1, H1.2, H2 for the control [non-PD] condition and PD1.1, PD1.2, PD2.1, PD2.2 for the PD condition) and the outcome was separately validated in the isogenic pair (PD3 vs PD3^corr^).

### Differentiation of iPSCs into midbrain-patterned neural progenitor cells

To generate ventral midbrain-patterned neural precursor cells (vmNPC) we used previously published methods^39,40^ on iPSCs dissociated to single cells with accutase (Thermo Fisher Scientific). On day 11, the cells were replated at 700,000 cells/cm^2^ on Geltrex and the culture medium was switched to NPC medium [Neurobasal, 2%, B27 supplement (minus vitamin A), (Thermo Fisher Scientific); 20 ng/ml fibroblast growth factor-2 (FGF-2; Peprotech, Cranbury, NJ, USA), and 20 ng/ml epidermal growth factor (EGF; Peprotech)] supplemented with 3 μM CHIR99021 (Stemgent, Cambridge, MA, USA), maintained thereafter in NPC medium. All reagents used in cell cultures are listed in Supplementary Table 4. Upon passage 1 or 2, NPCs were quality checked for regional identity (Supplementary Fig. 1). Only those NPCs that passed the quality control were used for differentiations.

### Differentiation of neural progenitor cells into astrocytes

vmNPCs were differentiated to astrocytes, as previously described^39^. After at least 4 w of expansion, vmNPCs were plated at 15,000 cells/cm^2^, continuously maintained in astrocyte differentiation medium (AM, Sciencell Research Laboratories, Carlsbad, CA, USA), and passaged weekly, until day 28. For maturation, iAstrocytes were plated at 30,000 cells/cm^2^ on Geltrex and on day 0, AM was switched to Astro-Maturation Medium [Advanced DMEM/F-12, 2% B27 (minus vitamin A), 1% Glutamax, 1% N2, 1% non-essential amino-acids, BMP4, and CNTF both 20 ng/μL; Peprotech], for at least 2 w. For the first 7 days of maturation the cells were treated with 0.5% FBS, and then FBS was fully withdrawn. Phenotypic analyses were performed between 28-42 days of differentiation and maturation.

### Differentiation of neural precursor cells into dopaminergic neurons

To prepare DAn, vmNPCs were harvested using accutase (Thermo Fisher Scientific) and plated at 600,000 cells/cm^2^ on Geltrex. Upon 80% confluency, vmNPCs were switched to neuronal differentiation medium [Neurobasal, 2% B27 supplement Plus, 1% N2 supplement, 1% GlutaMAX, 0.5 mM dibutyryl cyclic AMP (Sigma Aldrich, St. Louis, MO, USA), 10 μM DAPT (Tocris, Bristol, UK), 0.2 mM ascorbic acid (Sigma Aldrich), 20 ng/mL brain-derived growth factor (BDNF; Peprotech), 20 ng/mL glial-derived growth factor (GDNF; Peprotech), 1 ng/mL transforming growth factor-β3 (TGF-β3; Peprotech)]^39,40^, replenished every 2 d, for 21 d. Following differentiation, neurons were dissociated with accutase and plated on poly-L-ornithine (PLO; Sigma Aldrich)/Laminin (Sigma Aldrich)-coated plates (300.000 cells/cm^2^), or on iAstrocytes.

### Cytokine treatment of iAstrocytes

Serum was withdrawn from the culture medium for at least 48 h, before astrocytes were stimulated for 24 h with human recombinant TNFα (30 ng/mL, Peprotech), IL-1α (4 ng/mL, Peprotech), and C1q (400 ng/mL, Gentaur MyBiosource, San Diego, CA, USA), defined as “TIC-treatment”^45^.

### Phagocytosis assay (pHrodo-*E. coli*)

iAstrocytes were plated at 30,000 cells/cm^2^ on Geltrex coverslips and treated for 6 h with pHrodo-*E. coli* (2 μg/mL, Thermo Fischer Scientific). Cells were then fixed with 1% paraformaldehyde (Sigma Aldrich)/PBS for 15 min, at 22-23 °C. Images were acquired in TCS SP8P or TCS-SP5II confocal microscopes (LEICA Microsystems, Wetzlar, Germany).

### Treatment of iAstrocytes with αSyn preformed fibrils (PFFs)

Human αSyn preformed fibrils (PFFs) conjugated with AlexaFluor568 were prepared as described previously^142,143^. Differentiation day-38 iAstrocytes were plated at 20,000 cells/cm^2^ on Geltrex in AΜ. Forty-eight h later, αSyn-PFFs were diluted in AM (500 nM), sonicated, and added to astrocytes for 16 h, or added and washed after 2 h, while iAstrocytes remained in culture for 16 h. Cells were imaged fixed under a confocal spinning-disk microscope (Nikon Eclipse Ti2, Tokyo, Japan).

### Bafilomycin A1 treatment of iAstrocytes

iAstrocytes were plated at 40,000 cells/cm^2^ on Geltrex. Twenty-four h later, FBS was withdrawn and after 24 h, medium was changed to Astro-Maturation Medium with Bafilomycin A1 (100 nM, Cayman Chemicals, Ann Arbor, MI, USA) for 4 h, before lysis for Western blot. LC3II flux was calculated as the difference between Baf.A1-treated and untreated samples, relative to untreated and GAPDH.

### Lysosensor, Lysotracker DR, Magic Red assays

iAstrocytes were plated at 15,000 cells/Geltrex-coated 35-mm μ-dishes (Ibidi, Martinsried, Germany), in AM. Forty-eight h later, LysoSensor™ Green (1 μM, Thermo Fisher Scientific) or LysoTracker™ Deep Red (DR, 50 nM, Thermo Fisher Scientific) were added to the medium for 30 min, at 37 °C. Magic Red® Fluorescent Cathepsin B assay kit (ImmunoChemistry Technologies, Davis, CA, USA) was added for 15 min. Subsequently, cells were washed and imaged live in a confocal spinning-disk microscope (Nikon Eclipse Ti2) at 37°C. Images were acquired with identical settings across samples, in each experiment.

### Co-culture of iPSC-derived neurons and iAstrocytes

Differentiation day-28 astrocytes were plated at a density of 20,000-30,000 cells/cm^2^ on Geltrex in AM. When 70% confluent, FBS was withdrawn. The next day, iPSC-derived DAn (21 day of differentiation) were seeded on top of iAstrocytes at a density of 80,000 cells/cm^2^. The co-cultures were maintained in Neuron Basal medium, half replenished every 2-3 d, for 2 w.

### iAstrocyte-SH-SY5Y co-culture (SH-SY5Y loaded with PFFs)

Human neuroblastoma SH-SY5Y GFP^+^ cells were plated at 40,000 cells/cm^2^ in RPMI1640. Twenty-four h later, human αSyn-PFFs-were added for 16 h. PFF-loaded SH-SY5Y cells were trypsinized, mixed with either Ha or PDa at a 1:1 ratio, and sub-confluently seeded on Geltrex-coated coverslips in AM. After fixation, cells were stained for F-Actin with Phalloidin-AlexaFluor647 (1:300, Thermo Fisher Scientific), for 30 min, at 22-23 °C. All images were acquired under a confocal spinning-disk microscope (Nikon Eclipse Ti2).

### Calcium imaging

For calcium imaging, the Celldiscoverer 7 imaging system (Zeiss, Oberkochen, Germany) or ΙX81 Cell-R microscopes (Olympus, Tokyo, Japan) were used. Differentiation day-28 iAstrocytes were plated at densities 30,000-60,000/well in Geltrex-coated glass bottom 4-well plates (Ibidi) for maturation. Fourteen days later, Ha and PDa with similar densities were selected and incubated with 6 μM Fluo4-AM (Biotium, Fremont, CA, USA) in HBSS for 30 min, then washed twice and equilibrated with culture medium for 20 min in the microscope’s environmental chamber (37 °C, 5% CO_2_, >70% humidity). Cells were imaged at 488 nm. Sequential image series were acquired over 7 min in 3-4 fields, at a frequency of 1 frame/2 s. ATP (50 μM, Sigma Aldrich) was gently pipetted dropwise in the recording well to assess evoked activity in the same astrocytes. iNeurons in monocultures or co-cultured over 14 days with iAstrocytes were incubated in 2 μM Fluo4-AM with the adaptation process described above. Image series over 90 s, at 4 frames/s frequency were recorded in 3-4 fields. All recordings were performed using 20× lens, with 200 ms exposure for astrocytes and 100 ms for neurons, and 2×2 binning.

### Rabies-based monosynaptic retrograde tracing

To assess the synaptic connectivity of neurons in monocultures, we used an established rabies virus-based (RBV) retrograde monosynaptic tracing system^144^. Transduction of iNeurons in co-culture was done at least 5 days after replating onto iAstrocytes, using the lentivirus LV-hSyn-TVA-GFP (2.5 × 10^4^ IU/ml) for expression of TVA-receptor under control of the human synapsin promoter (starter neurons). Three days later, infection followed with the deletion-mutant virus RBV-ΔG-EnvA-RFP (3 × 10^4^ IU/ml) that entered starter neurons via the TVA-receptor. Then the RBV virus spread the infection once in presynaptic target neurons. Images were acquired after 10 days in a TCS SP8P confocal microscope (LEICA Microsystems). The relative connectivity index is calculated as the ratio of target neurons (RFP^+^/GFP^-^) retrogradely labeled through monosynaptic connections with starter neurons, over starter neurons (RFP^+^/GFP^+^).

### Immunofluorescence

Cells on glass coverslips were fixed with 4% paraformaldehyde (Sigma Aldrich) in PBS for 15 min at 22-23°C, blocked for 30 min with 5% normal donkey serum in PBS, containing 0.1-0.3% Triton X-100 (Sigma-Aldrich) for permeabilization (for intracellular epitopes) and subsequently incubated with primary antibodies (Supplementary Table 5) at 4 °C for 16-18 h, followed by incubation with the appropriate combination of secondary antibodies (Thermo Fisher Scientific) conjugated with AlexaFluor488, −546, or −647 for 2 h at 22-23 °C. Coverslips were mounted with StayBrite Hardset Mounting Medium with DAPI (Biotium).

### Thioflavin S staining

Following immunofluorescence staining, neuron-astrocyte co-cultures on glass coverslips were incubated with Thioflavin S (Sigma Aldrich, 0.05% w/v in 50% ethanol/water) for 15 min at 22-23 °C, and then washed twice with 50% ethanol/water, once with 80% ethanol/water, and once with 0.3% Triton/PBS.

### Protein aggregate detection

The presence of protein aggregates in iAstrocyte cultures and in co-cultures was examined using the PROTEOSTAT Aggresome Detection Kit (Enzo Life Sciences, Farmingdale, NY, USA), according to manufacturer’s instructions.

### Proteasome activity assay

iAstrocytes were plated on Geltrex-coated 12-well plates at 150.000 cells/well. The next day FBS was withdrawn for 24 h. Cells were collected in PBS, centrifuged (3800 × g, 5 min), resuspended in lysis buffer (25 mM Tris-HCl pH 7.6, 5 mM DTT, 2 mM ATP, 2 mM MgCl_2_ and 0.1 mM EDTA), and lysed by sonication followed by 20-min incubation on ice and centrifugation (16,000 × g, 5 min, 4 °C). Protein lysate (0.5 μg) was mixed with 100 μΜ fluorogenic substrate SUC-AMC (Abcam, Cambridge, UK) in 50 mM Tris-HCl pH 7.6, 5 mM DTT, 10 mM ATP, and 50 mM MgCl_2_, in dark 96-well microplates, and shaked (700 rpm, 10 min, 37 °C). Reactions were terminated by adding 100 μL of 5% SDS. Fluorescence was measured at 356 nm (excitation)/449 nm (emission) using a Biotek Synergy H1 multimode reader (Agilent, Santa Clara, CA, USA).

### Autophagic flux assay

iAstrocyte cultures were treated with 0.5 μΜ DALGreen and 0.1 μΜ DAPRed (Dojindo Molecular Technologies, Kumamoto, Japan) in Astro-Maturation Medium for 6 h. Cells were fixed and mounted with StayBrite Hardset Mounting Medium with DAPI (Biotium), before proceeding with image aquisition in TCS SP8P confocal microscope (LEICA Microsystems).

### Western blot

iAstrocytes were lysed for 15 min in ice-cold RIPA buffer containing PhosSTOP phosphatase inhibitors and protease inhibitors (Roche Life Science, Basel, Switzerland), sonicated, and centrifuged (20,000 × g). Total protein lysates (10-20 μg) were separated on 12% SDS-PAGE and transferred onto nitrocellulose (Porablot NCP, Macherey Nagel, Düren, Germany) or PVDF membrane (Porablot PVDF, Macherey Nagel) (for ULK1 and ITPKB). For p(Ser129)αSyn, the membrane was heated in PBS (65 °C, 16-18 h). Membranes were blocked with 3-5% non-fat milk, or 3% bovine serum albumin (BSA) in TBS/Tween-20, and then incubated (16-18 h, 4 °C) with primary antibodies (Supplementary Table 5), followed by the corresponding HRP-conjugated secondary antibodies (1:5000; Millipore Sigma, Burlington, MA, USA) for 1 h, and visualized using the Immobilon Crescendo Western HRP substrate (Millipore Sigma). Densitometric analysis was performed in FIJI (NIH)^145^.

### RNA isolation, cDNA Synthesis and RT-qPCR

Total RNA was extracted from cultured cells using the TRIzol Reagent (Thermo Fisher Scientific). Samples were processed with RQ1 DNAse (Promega Corporation, Madison, WI, USA) and further cleaned-up and concentrated using the RNA Clean and Concentrator kit (Zymo Research, Irvine, CA, USA). ProtoScript II Reverse Transcriptase (New England BioLabs, Ipswich, MA, USA) was used for first strand cDNA synthesis. PCR was performed in a StepOnePlus™ Real-Time PCR detection system (Applied Biosystems, Foster City, CA, USA) using the SYBR™ Select Master Mix (Applied Biosystems). Transcript expression was determined in triplicate reactions, and normalized to *GAPDH*. In quality control RT-qPCR of NPCs and iNeurons, relative mRNA levels were calculated for each gene using the 1/ΔC_T_ method. For *SNCA* expression, fold changes between conditions were calculated by the 2^-ΔΔCT^ method. Primers are listed in Supplementary Table 6.

### Preparation of cell lysates for proteomics

Ha and PDa clones analyzed by proteomics are shown in Supplementary Table 2. Day-28 iAstrocytes were plated in Geltrex-coated 6-well plates at 300,000 cells/well. Cells were maintained in AM and upon 60-70% confluency, serum was withdrawn for 72 h. iAstrocytes, untreated or TIC-treated, were lysed in 0.25 M Tris-HCl pH 8, 150 mM NaCl, 4% SDS, 0.1 M DTT, boiled for 4 min, sonicated (2 x 5 s), and centrifuged (17,000 × g, 15 min). The protein lysates were processed according to the Sp3 protocol^146^.

### LC-MS/MS analysis

Samples were run on a liquid chromatography tandem mass spectrometry (LC-MS/MS) setup consisting of a Dionex UltimateRSLC online with a Thermo Q Exactive HF-X Orbitrap mass spectrometer Thermo Fisher Scientific), as previously described^147^. At least two technical replicates were acquired per sample. The raw data were analyzed by the software DIA-NN against the canonical Uniprot human database.

### Data deposition

The mass spectrometry proteomics data have been deposited to the ProteomeXchange Consortium via the PRIDE partner repository with the dataset identifier PXD057491.

### Proteomic data analysis

For all analyses of the proteomics data, the statistical language R has been used (R Core Team (2022). [R: A language and environment for statistical computing. R Foundation for Statistical Computing, Vienna, Austria. URL https://www.R-project.org/]

### Data preprocessing

Samples with more than 1800 missing values were removed from the original dataset, yielding a protein data matrix of 7,653 proteins across 46 samples, 24 without and 22 with TIC-treatment. Data were log2 transformed and quantile normalization was subsequently applied. For batch correction, for each protein and batch, the average intensity across that batch was subtracted from its values while afterwards adding back the average intensity across all replicates. Then, for each protein its average value across the technical replicates corresponding to the same sample was computed. Principal component analysis (PCA) revealed that cell cycle had the largest effect on the dataset. For cell cycle correction, each protein was regressed against the first principal component. After corrections, PCA revealed clustering of samples by disease on the first principal component and by treatment on the second (Supplementary Fig. 4). Τhe top 200 proteins contributing most to the separation on the second principal component were identified, with 113 up- and 87 down-regulated after TIC treatment. Gene set enrichment analysis was subsequently applied to the up-regulated proteins, as described below. For Ha vs PDa analyses of samples without TIC treatment, data preprocessing was applied as described above. For comparisons between Ha and PDa only proteins without any missing values were considered, comprising 5,162 proteins across 24 untreated samples (Supplementary Table 2).

### Quality control of astrocyte differentiation

To check astrocytic differentiation, gene set enrichment analysis was performed. The MSig database C8^41^ – cell type signature gene sets – was tested against the list of proteins sorted in decreasing order of mean intensity across all samples using the one-sided Weighted Kolmogorov Smirnov test^148^. Of interest were the astrocyte-, (brain) neuron- and oligodendrocyte-related gene sets. P-values were obtained and mountain plots for selected gene sets were generated (all proteins were used).

### Differential expression analysis

For each protein a two-sided t-test comparing the two groups was applied (PDa vs. Ha, p-value threshold set to 0.01).

### Gene set enrichment analysis

To identify enriched gene sets, the DE proteins were tested against the three aspects of the Gene Ontology (GO) terms^149^: GO Biological Process (GOBP) database, GO Cellular Component (GOCC) and GO Molecular Function (GOMF). For that, the function ‘enrichGO’ from the Bioconductor package clusterProfiler^150^ was used (p-value threshold 0.01 without further adjustment). To remove redundancy between the enriched GO terms in each of the three databases, the function ‘simplify’ from the same package was subsequently used (similarity cutoff set to 0.6).

### STRING network

For network analysis, the STRING network for Homo sapiens (version 11.5)^151^ was used. For high confidence interactions, edges with a combined score not above 900 were filtered out, yielding 11,764 nodes and 115,262 edges. To identify dysregulated network regions in PDa, the DE proteins were mapped onto the sub-network and only the corresponding nodes and edges among them were kept. Of particular interest was the mTOR-related dysregulated network region.

### Image analysis

All image analyses were performed in FIJI (NIH)^145^. CD44^+^, VIMENTIN^+^, ALDH1L1^+^, and S100β^+^ iAstrocytes were counted manually and the percentage was calculated relative to total cells (DAPI). The number and size of p(Ser129)αSyn^+^ puncta were counted using the “Analyze Particles” plugin, keeping constant threshold settings, considering each cell area as region of interest (ROI). The number and size of protein aggregates (Aggresome) in iAstrocytes were counted using the “Aggrecount” plugin with constant threshold settings, and cell areas as ROIs. To assess astrocytic phagocytosis, the number and shape descriptors of pHrodo^+^ puncta were recorded using “Analyze Particles” with Auto Threshold, after denoising. In Ca^2+^ imaging experiment, each ROI (manually drawn in the cytoplasm in the case of astrocytes) was presented as relative fluorescence changes (ΔF/F0) after background subtraction. Area Under Curve (AUC), peak amplitude, and frequency were calculated. In co-cultures, MAP2^+^ and TH^+^ neurons per area (mm^2^) were counted manually. The number of neurites extending from the soma and branching of beta-3 tubulin^+^ neurons was manually determined. Neurite length was estimated by manually tracing all neurites on beta-3 tubulin^+^ neurons using the NeuronJ FIJI plugin^152^. Analysis of neuronal degeneration was performed in co-cultures, using previously published post-mortem findings as reference^58^. The number of p(Ser129)αSyn^+^ Lewy-like neurites, Lewy-like bodies, extracellular Lewy-body like formations, bulbous/dystrophic endings, axonal varicosities, and degenerating neurons were counted manually. The sum of total pathological features normalized with the sum of total counted neurons was defined as relative neuropathology index. In rabies-based monosynaptic retrograde tracing experiments, RBV-ΔG-EnvA-RFP^+^/LV-hSyn-TVA-GFP^-^ and RBV-ΔG-EnvA-RFP^+^/LV-hSyn-TVA-GFP^+^ neurons were manually counted. Relative connectivity was calculated as the ratio of RBV-ΔG-EnvA-RFP^+^/LV-hSyn-TVA-GFP^-^ over RBV-ΔG-EnvA-RFP^+^/LV-hSyn-TVA-GFP^+^ neurons. Neuronal aggresomes (Proteostat) were manually counted. For the analysis of uptaken αSyn-PFFs or SH-SY5Y-to-astrocyte transfer of αSyn-PFFs, the numbers of intracellular αSyn-PFFs in iAstrocytes and SH-SY5Y cells were counted per cell, using “Analyze Particles” with Auto Threshold, after denoising, considering cell areas as ROIs. The autophagic flux using DAPRed/DALGreen was estimated using JaCoP with Manders coefficients, to determine the level of colocalization; auto threshold was applied after filtering. In Lysotracker DR-labelled iAstrocytes, the number of lysosomes was quantified per cell, using the “Analyze Particles” FIJI tool with constant threshold settings and after denoising. Lysosomal acidity based on Lysotracker DR labeling was measured as integrated density per cell using constant threshold settings. The integrated density of LysoSensor green and Magic Red fluorescence was measured per cell with constant threshold settings, considering each cell area as ROI. Lysosomal positioning was quantified manually; using “Analyze Particles”, with auto threshold, the nuclei of the cells were used as ROIs. To determine the perinuclear area (10 μm from the nucleus) for each ROI, ‘Enlarge’ was applied. The absence of LAMP2^+^ puncta outside this area, designated cells with perinuclear lysosomes and their percentage over total cells was calculated.

### Statistical analysis

All experiments were performed in at least 3 biological replicates of healthy and PD samples. Biological replicate is defined at the level of different iPSC clones that astrocytes or neurons were derived from (in most cases) or independent differentiations (stated). In most experiments non-isogenic lines were examined separately from the isogenic pair, used to verify the causality of the mutation, unless otherwise stated. For each readout, the exact details of analysis and statistics are given in figure legends. Statistical analysis was performed in GraphPad Prism (San Diego, CA, USA) and p value for significance was set to ≤ 0.05.

## Data availability

The proteomics data generated in this study are available via ProteomeXchange with identifier PXD057491.

## Acknowledgements

We would like to thank Dr Georgia Kouroupi for transferring iPSC expertise; Drs Nicoletta Selenti, Christalena Sofocleous, and J. Traeger-Synodinos (Laboratory of Medical Genetics, St. Sophia’s Children’s Hospital, Medical School, National and Kapodistrian University of Athens, Greece) for conducting the karyotypic analyses of the iPSC lines used in this study; Dr Andreas Beyer (Cluster of Excellence on Cellular Stress Responses in Aging-associated Diseases (CECAD), University of Cologne, Cologne, Germany) for supervising the statistical analyses of the proteomics data; Dr Katerina Segklia (Hellenic Pasteur Institute) for technical and project management support; Dr Panagiotis Handris (Hellenic Pasteur Institute) for scientific insight; Dr Rudolf Jaenisch (Whitehead Institute for Biomedical Research Cambridge, Massachusetts, USA) for providing the isogenic pair of iPSCs [WIBR-iPS-SNCA^A53T (PD3) and WIBR-iPS-SNCA^A53T-Corr (PD3^corr^)].

## Funding

This work was supported by the Hellenic Foundation for Research and Innovation (H.F.R.I.) under the “1st Call for H.F.R.I. Research Projects to support Faculty members and Researchers and the procurement of high-cost research equipment” (Project Number: HFRI-FM17C3-1019) [RM]; PIU – Pasteur International Unit for Neurodegenerative diseases established between the Hellenic Pasteur Institute and the Institut Pasteur in Paris [RM and CZ]; the General Secretariat for Research and Innovation under the action “National research network to elucidate the genetic basis of Alzheimer’s and Parkinson’s neurodegenerative diseases, detect reliable biomarkers, and develop innovative computational technologies and therapeutic strategies on the basis of precision medicine” ΤΑΑ TAEDR-0535850 - Brain Precision [European Union (NextGenerationEU); National Recovery and Resilience Plan Greece 2.0] [ET]; MS acknowledges support from the project “The Greek Research Infrastructure for Personalised Medicine (pMED-GR)” (MIS 5002802) which is implemented under the Action “Reinforcement of the Research and Innovation Infrastructure”, funded by the Operational Programme “Competitiveness, Entrepreneurship and Innovation” (NSRF 2014-2020) and co-financed by Greece and the European Union (European Regional Development Fund). SD was supported by a Bodossaki Foundation scholarship for visiting research scientists and a Federation of FENS (European Neuroscience Societies)]/ IBRO (International Brain Research Organization)] - Pan-Europe Regional Committee (PERC) Exchange Fellowship.

## Author contributions

CP performed most experiments involving iAstrocytes, including cell culture and characterization, imaging, image data analyses, Western blots, etc. OA carried out most experiments involving generation and characterization of iNeurons, including co-cultures, imaging, image data analyses, etc. AK conducted the Ca^2+^ imaging experiments and analyses, contributed to cell culture maintenance, differentiations, quality control, and Western blots. SD performed and analyzed experiments involving PFFs and live lysosomal tracking with help from FrP who participated in the experimental design and data interpretation. CP, OA, AK, and SD contributed to experimental design, data interpretation, figure and manuscript preparation. KC performed the preprocessing, statistical analyses and documentation of the proteomics data. MS performed the LC-MS/MS analysis to generate the proteomic data set. KD conducted the proteasome activity assay, supervised by EE. CZ provided resources, contributed to experimental design and data interpretation, and supervision of SD and FrP. RM conceived the study, provided funding, resources, scientific insight, and participated in experimental design and data interpretation. ET participated in conceptualization, experimental design and data interpretation, and provided funding and resources. FlP formulated the working hypothesis, designed, coordinated, and supervised experiments and interpreted the data; FlP wrote the original draft; RM reviewed, revised and edited the manuscript that was finalized with input from all authors.

## Competing interests

The authors declare no competing interests

## Notes

### Competing Interest Statement

The authors have declared no competing interest.

### Summary of Updates

There was an error in the numbering of authors' affiliations that has now been corrected.

