## Supplementary material for "Proteostasis dysregulation in p.A53T-α-Synuclein iPSC-derived astrocytes exacerbates neurodegeneration in a Parkinson’s disease model with Lewy-like pathology": SUPPL. TABLE 3

| Protein name | P-value | Folds Change |
|---|---|---|
| PDLIM3 | 0.000855057 | 2.039388243 |
| TUBA1C | 0.002686858 | 1.923808603 |
| SLC6A6 | 0.000147436 | 1.824682117 |
| TPM4 | 0.001638435 | 1.788507346 |
| CLMN | 0.001348142 | 1.729626764 |
| IGFBP7 | 0.000893899 | 1.623598587 |
| LTBP2 | 0.009540248 | 1.610679459 |
| NNMT | 0.00367998 | 1.562713067 |
| ALAS1 | 0.002527724 | 1.547438814 |
| CASTOR1 | 0.000130381 | 1.543613973 |
| CD82 | 0.006036219 | 1.434564421 |
| ITPKB | 0.001284676 | 1.408427918 |
| SDC2 | 0.000345457 | 1.404821879 |
| LGMN | 0.003054122 | 1.382919957 |
| TP53I11 | 0.007956608 | 1.335250746 |
| VLDLR | 0.007564163 | 1.263249807 |
| PFKFB2 | 0.000598988 | 1.190268635 |
| PRXL2A | 0.001262204 | 1.187789724 |
| SERPINB1 | 0.000183236 | 1.124583115 |
| GALC | 0.000187219 | 1.060836672 |
| F5 | 0.002213506 | 1.051832877 |
| ABCA3 | 0.003973395 | 1.029665595 |
| SLC26A2 | 0.001961267 | 1.010865022 |
| IDUA | 0.009721333 | 1.010487981 |
| LARP6 | 0.003014574 | 1.009321104 |
| CDH11 | 0.000220758 | 1.006260031 |
| BLVRB | 0.001580821 | 0.991946674 |
| COL6A2 | 0.008400027 | 0.975605072 |
| COL6A1 | 0.003145751 | 0.961039376 |
| MGST3 | 0.008380262 | 0.935473056 |
| TPCN1 | 0.000173353 | 0.93125564 |
| WASHC2C | 0.00667371 | 0.931026698 |
| FBN2 | 0.002909966 | 0.904413694 |
| ITPR2 | 0.001404121 | 0.902717707 |
| SNCA | 0.00185344 | 0.902388976 |
| DICER1 | 0.000913306 | 0.890961404 |
| SLC29A1 | 5.01E-06 | 0.886706732 |
| FTL | 2.56E-05 | 0.878727168 |
| SLC12A2 | 0.006716455 | 0.866466363 |
| PRKG1 | 0.008608009 | 0.853828213 |
| MAST4 | 0.002436467 | 0.838890564 |
| CORO7 | 0.000406192 | 0.83775342 |
| LGALS8 | 0.002311109 | 0.810951265 |
| CPPED1 | 0.0003028 | 0.804592177 |
| MCM7 | 0.004352493 | 0.802143696 |
| EML2 | 0.000319134 | 0.794447582 |
| WDFY2 | 0.006122155 | 0.783508418 |
| TRIM22 | 0.000139965 | 0.768752588 |
| MT-ND1 | 0.003344184 | 0.758972712 |
| TMEM245 | 0.004775973 | 0.747645727 |

| | | |
|---|---|---|
| QPRT | 0.005903283 | 0.735856452 |
| CEMIP2 | 0.000971982 | 0.735038811 |
| ACSL3 | 0.003205287 | 0.730421025 |
| ATP1B1 | 0.006376079 | 0.728525333 |
| NIPSNAP3A | 0.007506114 | 0.722326706 |
| TMEM65 | 0.009867096 | 0.710054442 |
| FAM210A | 0.001726626 | 0.703433177 |
| CDC42BPA | 0.001680968 | 0.682520745 |
| KIF16B | 0.002969055 | 0.67517687 |
| SLC39A11 | 0.000607213 | 0.672853845 |
| EXOC6 | 0.002132083 | 0.671324181 |
| DMXL1 | 0.000530506 | 0.666235731 |
| DHRSX | 0.006587645 | 0.658827865 |
| SDF2L1 | 0.001565808 | 0.658765166 |
| PRMT7 | 0.008136994 | 0.657668956 |
| DNAJC21 | 0.008706147 | 0.648971202 |
| HSD17B11 | 0.007650025 | 0.64327159 |
| CPT1A | 2.42E-05 | 0.624795732 |
| ALG11 | 0.003667031 | 0.624766139 |
| SACM1L | 0.000219818 | 0.606303931 |
| DHRS4 | 8.92E-05 | 0.592581045 |
| WNK1 | 0.000933701 | 0.592018778 |
| ARL8B | 0.000274335 | 0.581659669 |
| MMTAG2 | 0.00983699 | 0.577618518 |
| ARHGEF28 | 0.004916106 | 0.574539321 |
| RAB12 | 0.00477115 | 0.568482364 |
| RAB6A | 0.001075085 | 0.564896009 |
| ATL3 | 0.001036011 | 0.564555809 |
| NEMP1 | 0.000397607 | 0.557236825 |
| RHEB | 0.009194604 | 0.536101545 |
| FIG4 | 0.002353486 | 0.535137399 |
| GSN | 0.002521443 | 0.521731794 |
| RAB4B | 0.007017326 | 0.520039435 |
| TRIM26 | 0.000670123 | 0.519740613 |
| KIF3A | 0.000383056 | 0.506718758 |
| EOGT | 0.002259018 | 0.506218009 |
| SOGA1 | 0.006760061 | 0.504301461 |
| SORT1 | 0.003998166 | 0.503599664 |
| ANXA4 | 0.001495841 | 0.503095108 |
| UAP1L1 | 0.00073631 | 0.50151185 |
| PIEZO1 | 0.002217957 | 0.4989467 |
| ACOT7 | 0.007531966 | 0.497089032 |
| APOL2 | 0.000810323 | 0.483741204 |
| SERPINB6 | 0.00058994 | 0.472241698 |
| AP3D1 | 0.003594954 | 0.460858285 |
| PPIL4 | 0.001987745 | 0.453196941 |
| GLS | 0.001843921 | 0.439340305 |
| TGFBRAP1 | 0.005147419 | 0.435557593 |
| AGTRAP | 0.004062604 | 0.431514816 |
| AKAP13 | 0.004487771 | 0.431335025 |
| LAMTOR3 | 0.006633991 | 0.43083492 |

| | | |
|---|---|---|
| PDE6D | 0.001746749 | 0.429123501 |
| PFKM | 0.003935311 | 0.428096175 |
| LRP1 | 0.001925083 | 0.427573263 |
| RDH11 | 0.006843453 | 0.426572425 |
| CTBS | 0.005755557 | 0.425960868 |
| ATP1A1 | 0.004969051 | 0.421440474 |
| TMTC3 | 0.009747944 | 0.419833526 |
| LAMA4 | 0.000366257 | 0.41970648 |
| MTMR9 | 0.007262385 | 0.414396591 |
| RAB22A | 0.003402193 | 0.409012639 |
| GUSB | 0.002855767 | 0.40165747 |
| SOAT1 | 0.008537998 | 0.399857654 |
| ATP9A | 0.007828919 | 0.396472159 |
| ARFGEF2 | 0.000453156 | 0.395813145 |
| ABCD3 | 0.002035802 | 0.39055628 |
| ALS2 | 0.006116059 | 0.390148549 |
| ARRB1 | 0.008292436 | 0.384887239 |
| FBXW9 | 0.000243133 | 0.376874517 |
| SCAMP3 | 0.005846608 | 0.364505608 |
| PFKL | 0.009285847 | 0.359975194 |
| WDFY1 | 0.002585953 | 0.357016989 |
| NUMB | 0.004809155 | 0.356428803 |
| SETDB1 | 0.001130562 | 0.352671547 |
| DCTN6 | 0.00489016 | 0.349636845 |
| FLCN | 0.008042072 | 0.347942689 |
| EMC1 | 0.008126443 | 0.337766854 |
| DDX10 | 0.003313073 | 0.321661552 |
| NUDT5 | 0.005283732 | 0.320792449 |
| HADH | 0.008619507 | 0.319623717 |
| BMP2K | 0.003985434 | 0.309389544 |
| SLC27A4 | 0.004234537 | 0.291888836 |
| TARBP2 | 0.005094361 | 0.290903136 |
| TMEM63B | 0.003868198 | 0.288454148 |
| TBC1D2B | 0.008601531 | 0.284997404 |
| CFDP1 | 0.003605052 | 0.279416363 |
| NPC1 | 0.000245741 | 0.278804096 |
| RAB9A | 0.003446278 | 0.276376243 |
| CSNK1D | 0.007565706 | 0.256093183 |
| PPP2R2D | 0.001245666 | 0.254634211 |
| BCS1L | 0.002071394 | 0.249750785 |
| PKD2 | 0.009894613 | 0.247254518 |
| WDR13 | 0.007192516 | 0.241615976 |
| SORD | 0.006222504 | 0.239078181 |
| RPN1 | 0.007570326 | 0.219522005 |
| TFIP11 | 0.00690356 | 0.198771902 |
| KIF1B | 8.17E-05 | 0.170331247 |
| PANK4 | 0.009171087 | 0.146958216 |
| MTMR1 | 0.008250052 | -0.139384869 |
| FARSA | 0.000409044 | -0.17523966 |
| SAP18 | 0.008939458 | -0.176104112 |
| ABCE1 | 0.009585125 | -0.197438184 |

| | | |
|---|---|---|
| ACTR10 | 0.007201597 | -0.201027909 |
| RARS1 | 0.004363678 | -0.21808342 |
| ECPAS | 0.001513174 | -0.224385786 |
| RIPOR1 | 0.002663946 | -0.224545103 |
| POLDIP2 | 0.00576739 | -0.232352328 |
| EIF3H | 0.007980729 | -0.236396704 |
| SNUPN | 0.007001245 | -0.237022851 |
| DHX29 | 0.003710832 | -0.23851812 |
| PLEC | 0.001031974 | -0.246037441 |
| SIK2 | 0.006697934 | -0.253334577 |
| ADSL | 0.008938598 | -0.261807191 |
| TMEM43 | 0.000365575 | -0.270820038 |
| MKRN2 | 0.006578986 | -0.271842609 |
| ATP6V0D1 | 0.005372124 | -0.271872056 |
| TARS1 | 0.009000978 | -0.276203092 |
| CARS1 | 0.007924388 | -0.277462345 |
| METAP1 | 0.007402006 | -0.286433514 |
| MAT2B | 0.001768298 | -0.287003923 |
| MTHFD1 | 0.006467325 | -0.287386984 |
| MTAP | 0.003987762 | -0.296280817 |
| FAM120A | 0.000754097 | -0.297144028 |
| XRCC5 | 0.008922365 | -0.307916977 |
| RPL28 | 0.004235181 | -0.308958291 |
| TOMM40 | 0.002711592 | -0.317003617 |
| ATIC | 0.002629579 | -0.32950146 |
| EPB41 | 0.008894971 | -0.330447114 |
| RBM14 | 0.00710191 | -0.331939973 |
| ARHGEF7 | 0.008088491 | -0.334228369 |
| SELENON | 0.004930564 | -0.334237035 |
| THOP1 | 0.003751291 | -0.340682415 |
| PSMC6 | 0.003667312 | -0.343522751 |
| SLC44A2 | 0.006413595 | -0.343940381 |
| PYROXD2 | 0.008729181 | -0.34884584 |
| IGF2BP3 | 0.007638262 | -0.350457199 |
| KPNA3 | 0.00719416 | -0.351422991 |
| OTUB1 | 0.001597573 | -0.353070037 |
| MPST | 0.004083726 | -0.354796239 |
| PANK2 | 0.004754691 | -0.355954743 |
| MIOS | 0.002889075 | -0.360829424 |
| CNRIP1 | 0.002886856 | -0.361655056 |
| CNP | 0.003315202 | -0.364980247 |
| PKN1 | 0.009250753 | -0.36662637 |
| GPS1 | 0.006656069 | -0.36818794 |
| PFAS | 0.00479793 | -0.368556364 |
| RPL17 | 0.004875194 | -0.370268748 |
| SRP19 | 0.001125098 | -0.371626707 |
| GUF1 | 0.00147769 | -0.372654352 |
| DDX17 | 0.00677349 | -0.374790358 |
| SEPTIN8 | 9.01E-05 | -0.376338718 |
| CHPF2 | 0.001532799 | -0.377215156 |
| EIF2B2 | 0.007958837 | -0.379579934 |

| | | |
|---|---|---|
| PSMC2 | 0.006319097 | -0.386438659 |
| IFT88 | 0.001659455 | -0.390084162 |
| TSG101 | 0.006654343 | -0.390416953 |
| XPO5 | 0.009879199 | -0.391243664 |
| TEX10 | 0.001899137 | -0.391757354 |
| ATAD3A | 0.002856083 | -0.392041684 |
| PSMC1 | 0.001508096 | -0.394746569 |
| SNRPD2 | 0.006404511 | -0.399911865 |
| LRSAM1 | 0.00957846 | -0.401392085 |
| RPL37A | 0.005510304 | -0.401748475 |
| ZC3H7B | 0.000565634 | -0.405587109 |
| SRP72 | 0.003070785 | -0.405697273 |
| HNRNPL | 0.004478053 | -0.40598565 |
| SEC23IP | 0.003486472 | -0.408289227 |
| EXOC3 | 0.006619301 | -0.408416019 |
| PRPF4B | 0.008448546 | -0.408880273 |
| WASL | 0.007060296 | -0.412176592 |
| HNRNPH1 | 0.009901644 | -0.415127246 |
| RAD23B | 0.002575486 | -0.421672501 |
| LAMTOR2 | 0.005280887 | -0.422288463 |
| AARSD1 | 0.001493763 | -0.424468917 |
| TRABD | 0.003356122 | -0.424526697 |
| IMPACT | 0.00798477 | -0.42748661 |
| STAU2 | 0.002215353 | -0.428028374 |
| POLR1E | 0.00313727 | -0.428279645 |
| GLYR1 | 0.004616959 | -0.430206799 |
| ELMO2 | 0.006491528 | -0.431669829 |
| NUP98 | 0.004443079 | -0.434279577 |
| ADK | 0.009439997 | -0.434982781 |
| CASP6 | 0.004518984 | -0.435003007 |
| THOC5 | 0.006034035 | -0.435409021 |
| DNM1L | 0.00477142 | -0.43808206 |
| FIS1 | 0.001050277 | -0.43826101 |
| INTS10 | 0.008225675 | -0.447544498 |
| XRCC6 | 0.006516782 | -0.447989823 |
| MRM3 | 0.003295641 | -0.4480244 |
| EPM2AIP1 | 0.006784881 | -0.451232789 |
| ARHGAP21 | 0.003233275 | -0.451373627 |
| RPF2 | 0.00018706 | -0.452432017 |
| HNRNPR | 0.006633829 | -0.456174451 |
| FUBP1 | 0.008792133 | -0.457136395 |
| INTS6 | 0.005383587 | -0.458038189 |
| RCC1L | 0.000145205 | -0.458050603 |
| DHRS1 | 0.003958211 | -0.459547652 |
| PRPF38B | 0.004108053 | -0.460766264 |
| RBM4 | 0.003352052 | -0.461820749 |
| GTF2I | 0.001573173 | -0.462501622 |
| RBMS2 | 0.002494503 | -0.463015133 |
| HNRNPA2B1 | 0.001973464 | -0.463942849 |
| TLN1 | 0.00805561 | -0.46694519 |
| DDX56 | 0.002679421 | -0.46737088 |

| | | |
|---|---|---|
| PDCD6IP | 0.000759556 | -0.472746919 |
| EIF4G3 | 0.00245222 | -0.473279447 |
| OXCT1 | 0.007027009 | -0.474038888 |
| GCLC | 0.003421241 | -0.474210437 |
| KPNA6 | 0.000592377 | -0.47679387 |
| NEK9 | 0.001220257 | -0.480641532 |
| FAM120B | 0.000473257 | -0.481549132 |
| WARS1 | 0.00185868 | -0.483105706 |
| LPCAT1 | 0.006428114 | -0.489147788 |
| CHCHD6 | 0.006833943 | -0.491734073 |
| MSI2 | 0.000872853 | -0.492392688 |
| FTSJ1 | 0.00687269 | -0.493152719 |
| FAM98A | 0.004650264 | -0.493266288 |
| EZR | 0.000691012 | -0.494771067 |
| TRMT10C | 0.002219324 | -0.495346651 |
| RPP38 | 0.006717281 | -0.498406095 |
| POLDIP3 | 0.002767784 | -0.500016778 |
| LRPPRC | 0.005457737 | -0.500539903 |
| ACAD8 | 0.007906281 | -0.500621966 |
| MID1 | 0.003889322 | -0.501238732 |
| CIAPIN1 | 9.72E-05 | -0.502104278 |
| BECN1 | 0.00454357 | -0.50386338 |
| AIMP2 | 0.001499377 | -0.504981802 |
| TRMT5 | 0.003904388 | -0.506645636 |
| DNAAF5 | 0.003600531 | -0.50755889 |
| CD2AP | 0.0084082 | -0.50894385 |
| SRP68 | 0.008551133 | -0.511104552 |
| PPP1R21 | 0.001170696 | -0.511367388 |
| SNX1 | 0.004514205 | -0.513167186 |
| NAA15 | 0.000402478 | -0.515080827 |
| RPL4 | 0.001408335 | -0.516898924 |
| FNTA | 0.006810158 | -0.521316809 |
| RIN1 | 0.009536416 | -0.523816453 |
| PTPRS | 0.001481655 | -0.524923767 |
| RPL19 | 0.008908336 | -0.527700318 |
| SUMF2 | 0.008065025 | -0.528881755 |
| BPGM | 0.002767237 | -0.536029698 |
| PAFAH1B3 | 0.000532383 | -0.542567438 |
| SMNDC1 | 0.003457982 | -0.545508612 |
| MVD | 0.008639011 | -0.546871722 |
| PPP6R2 | 0.004319965 | -0.547934595 |
| HARS1 | 0.002901647 | -0.549069281 |
| SSH1 | 0.00729222 | -0.551520838 |
| SKP1 | 0.007241267 | -0.55646708 |
| PPIE | 0.003091468 | -0.557948562 |
| G3BP1 | 0.006410276 | -0.559898428 |
| OSBPL10 | 0.005940109 | -0.561670627 |
| CDC5L | 0.004250871 | -0.567861675 |
| STUB1 | 0.00843892 | -0.569703532 |
| KARS1 | 0.001267231 | -0.571431825 |
| RPL7 | 0.001705013 | -0.57145844 |

| | | |
|---|---|---|
| BCAT1 | 0.001316139 | -0.573082613 |
| MLH1 | 0.003310942 | -0.577749089 |
| ERP29 | 0.003164589 | -0.582091517 |
| NOL6 | 0.00364912 | -0.583282324 |
| DNAJC9 | 0.000805422 | -0.583594848 |
| PUM3 | 0.002321658 | -0.585137453 |
| MAPKAPK3 | 0.004973436 | -0.585280978 |
| PDXDC1 | 0.009521539 | -0.587634429 |
| EXOC7 | 0.00397842 | -0.590656442 |
| OLA1 | 0.000141395 | -0.595298889 |
| PML | 0.008175967 | -0.607741524 |
| FHOD1 | 0.002388294 | -0.6130921 |
| PRPF6 | 0.001658626 | -0.616951149 |
| ANKRD13A | 0.009754882 | -0.624874745 |
| RPS15 | 0.000306911 | -0.628054168 |
| RRP1B | 0.002046185 | -0.628611919 |
| EIF3E | 0.00641237 | -0.628691241 |
| MRRF | 0.000300324 | -0.629933616 |
| KEAP1 | 0.001128392 | -0.633217596 |
| DNAJC7 | 0.00032597 | -0.635296571 |
| SF3B1 | 0.00066467 | -0.638458962 |
| RPL10A | 0.000815499 | -0.639534402 |
| PSPC1 | 0.004399383 | -0.639655495 |
| DIAPH1 | 0.000189979 | -0.63991312 |
| GPC6 | 0.004125445 | -0.640196814 |
| TOMM70 | 0.004063412 | -0.640744221 |
| EHD1 | 0.000457828 | -0.645437241 |
| STAU1 | 0.001138057 | -0.646207273 |
| MRPL58 | 0.008546 | -0.648745098 |
| ANAPC5 | 0.004654134 | -0.65464512 |
| XRN2 | 0.00016269 | -0.657708799 |
| HSPD1 | 0.006567117 | -0.658646581 |
| ABHD12 | 0.004390231 | -0.659378716 |
| NIBAN2 | 0.000824692 | -0.659971447 |
| STARD13 | 1.83E-05 | -0.660628628 |
| CSTF3 | 0.000453292 | -0.661662481 |
| UNC119B | 0.003178459 | -0.663529791 |
| RPL13 | 0.008179119 | -0.66439059 |
| GTPBP10 | 0.004684512 | -0.664981413 |
| ACTN4 | 0.002224649 | -0.666490972 |
| DST | 0.002032725 | -0.666672506 |
| SMARCD2 | 0.001360527 | -0.677469248 |
| CENPV | 0.008802023 | -0.678372799 |
| CCAR2 | 0.001402909 | -0.683754409 |
| MAP2K3 | 0.006695093 | -0.683884242 |
| EIF4G2 | 0.002585351 | -0.684109506 |
| SELENBP1 | 0.003359767 | -0.693262303 |
| ITGA6 | 0.005525871 | -0.694528989 |
| MAD1L1 | 0.003947292 | -0.695344537 |
| UNC45A | 0.001129041 | -0.697878427 |
| C11orf98 | 0.005607912 | -0.697986462 |

| | | |
|---|---|---|
| CCDC124 | 0.005179803 | -0.701393944 |
| EWSR1 | 0.000726989 | -0.701567575 |
| SFPQ | 0.003906879 | -0.703883698 |
| THUMPD1 | 0.004610067 | -0.704901428 |
| PAFAH1B2 | 0.007181869 | -0.707075563 |
| ILF3 | 0.001924049 | -0.7072104 |
| CNOT2 | 0.000509345 | -0.707285926 |
| JUND | 0.003480781 | -0.712752311 |
| C8orf33 | 0.002060746 | -0.713912814 |
| NELFB | 0.009074692 | -0.715337438 |
| MAN1A1 | 0.009466668 | -0.715733035 |
| EHD4 | 0.006859968 | -0.716943023 |
| SIRT2 | 0.004664928 | -0.717033047 |
| HPRT1 | 0.00963905 | -0.718177307 |
| API5 | 0.001708955 | -0.718791213 |
| PIP4K2A | 0.001168791 | -0.721105357 |
| RFC2 | 0.000346631 | -0.725581568 |
| FMNL2 | 0.000470173 | -0.726396897 |
| SEPTIN10 | 0.001970975 | -0.732011963 |
| ALDH7A1 | 0.004430666 | -0.733599037 |
| TP53BP2 | 0.009054417 | -0.744452293 |
| TENM4 | 0.006243948 | -0.756771112 |
| SRSF9 | 0.001535099 | -0.759228162 |
| ANKMY2 | 0.008281957 | -0.759751095 |
| TAB2 | 0.00249792 | -0.759969713 |
| ZNF512 | 0.001129836 | -0.760653966 |
| TTC39C | 0.006494704 | -0.762600542 |
| GATAD2A | 0.003124437 | -0.7642526 |
| REXO4 | 0.001504991 | -0.765612256 |
| DNAJC8 | 0.00258939 | -0.777759378 |
| UTP6 | 0.000282289 | -0.779140313 |
| HABP4 | 0.002929212 | -0.780610938 |
| ANXA3 | 0.009611208 | -0.790490168 |
| MRPL1 | 0.001482115 | -0.805346077 |
| DMAP1 | 0.004052015 | -0.805388069 |
| ARHGAP18 | 0.003438693 | -0.806606077 |
| MICU1 | 0.001890023 | -0.80884828 |
| TANGO2 | 0.005568239 | -0.812876997 |
| VWA5A | 0.004031426 | -0.814206244 |
| BYSL | 0.000347368 | -0.819141113 |
| BAG2 | 0.003989849 | -0.83723539 |
| RAVER1 | 0.002437832 | -0.839478443 |
| RAI1 | 0.001980136 | -0.841264532 |
| INA | 0.002701552 | -0.842512656 |
| BZW1 | 0.001868702 | -0.846104161 |
| NANS | 0.000358645 | -0.85228276 |
| NOB1 | 0.006947855 | -0.853317898 |
| H1-10 | 0.000856572 | -0.860514123 |
| CRMP1 | 0.001054573 | -0.86878823 |
| PPP3CA | 0.001239772 | -0.875096757 |
| C7orf50 | 0.001716923 | -0.875453922 |

| | | |
|---|---|---|
| ERI1 | 0.0009677 | -0.877253513 |
| PPP2R5E | 0.003284287 | -0.878874725 |
| TXLNG | 0.0023492 | -0.880523211 |
| RAB3B | 0.005450526 | -0.892160087 |
| GDAP1 | 0.002358233 | -0.899430715 |
| PRSS23 | 0.003105636 | -0.908509585 |
| CSNK2B | 0.003247069 | -0.910526438 |
| EFHD2 | 0.001556223 | -0.913519429 |
| SNW1 | 0.003440509 | -0.914805303 |
| S100A10 | 0.002632959 | -0.928934851 |
| TBC1D2 | 0.005239032 | -0.93500024 |
| PLCD1 | 0.007912137 | -0.93587598 |
| UBE2L6 | 0.008689707 | -0.936671556 |
| PHGDH | 0.00254441 | -0.938666803 |
| SPATS2L | 0.006549648 | -0.946543204 |
| NUMA1 | 0.002540018 | -0.950317704 |
| SDC4 | 0.000282922 | -0.952196517 |
| MYLK | 0.000279148 | -0.96111374 |
| RPS25 | 0.00723098 | -0.96673294 |
| GULP1 | 0.000231769 | -0.997602686 |
| RBMX | 0.000873991 | -1.000666163 |
| SARM1 | 0.004136327 | -1.003629442 |
| RRS1 | 0.000717266 | -1.006288408 |
| PGRMC1 | 0.002589216 | -1.010018348 |
| NGDN | 0.002765511 | -1.02905199 |
| SMAD3 | 1.61E-05 | -1.048590586 |
| LRRC59 | 0.000284075 | -1.063660604 |
| RPL35 | 0.003378539 | -1.069585541 |
| MAGED1 | 0.000856716 | -1.099342816 |
| TM4SF1 | 0.000184388 | -1.110023134 |
| AHR | 0.000182176 | -1.118418857 |
| ANXA2 | 0.001923949 | -1.129603118 |
| DCBLD2 | 0.004817748 | -1.130492078 |
| LUC7L | 0.002177999 | -1.181945124 |
| SAMHD1 | 0.00085051 | -1.18236731 |
| KRR1 | 0.001417112 | -1.215440533 |
| BCAM | 0.000328187 | -1.369820587 |
| TNFAIP2 | 0.006944811 | -1.384224075 |
| RPL29 | 0.007457694 | -1.391376366 |
| H1-5 | 0.001038187 | -1.527209513 |
| H1-4 | 0.003158298 | -1.552187371 |
| TJP2 | 9.42E-06 | -1.614743058 |
| LOX | 0.000197512 | -1.632569508 |
| OLFML2A | 0.008128865 | -1.68057862 |
| HMGA1 | 0.005606698 | -1.903563236 |
| RBP1 | 0.000139832 | -2.081787032 |
| KRT19 | 0.000570549 | -3.713214564 |
