## Supplementary material for "Proteostasis dysregulation in p.A53T-α-Synuclein iPSC-derived astrocytes exacerbates neurodegeneration in a Parkinson’s disease model with Lewy-like pathology": SUPPL. TABLE 4

| NAME | MANUFACTURER | CATALOG NUMBER |
| --- | --- | --- |
| DMEM High Glucose w/ L-Glutamine w/ Sodium Pyruvate | Biosera, Nuaille, France | LM-D1111/500 |
| Hanks' Balanced Salts Solution w/o Ca w/o Mg w/o Na Bicarbonate w/o Phenol Red | Biosera, Nuaille, France | LM-S2034/500 |
| Fluo4-AM | Biotium, Fremont, CA, USA | 50018 |
| Hanks' Balanced Salts Solution w/o Ca w/o Mg w/ Sodium Bicarbonate w/o Phenol Red | Biowest, Nuaille, France | L0607 |
| DAPT (N-[2S-(3,5-difluorophenyl)acetyl]-L-alanyl-2-phenyl-glycine, 1,1-dimethylethyl ester) | Cayman Chemicals, Ann Arbor, MI, USA | 13197 |
| Bafilomycin A1 | Cayman Chemicals, Ann Arbor, MI, USA | 11038.5 |
| Corning® Matrigel® hESC-Qualified Matrix, LDEV-free | Corning, Corning, NY, USA | 354277 |
| C1q | Gentaur MyBiosource, San Diego, CA, USA | MBS147305 |
| BMP4 | Peprotech, Cranbury, NJ, USA | AF-120-05ET-B |
| CNTF | Peprotech, Cranbury, NJ, USA | 450-13-B |
| EGF | Peprotech, Cranbury, NJ, USA | GMP100-15 |
| FGF-Basic (146 a.a.) Human Recombinant (FGF-2) | Peprotech, Cranbury, NJ, USA | 100-18C-0100 |
| Human FGF-8b | Peprotech, Cranbury, NJ, USA | 167100-25-B |
| IL-1α | Peprotech, Cranbury, NJ, USA | 200-01A-B |
| Recombinant Human GDNF | Peprotech, Cranbury, NJ, USA | 450-10-50µG |
| Recombinant Human TGF Beta 3 | Peprotech, Cranbury, NJ, USA | 100-36E-A |
| TNF-α | Peprotech, Cranbury, NJ, USA | 300-01A |
| Recombinant Human BDNF Protein | R&D Systems, Minneapolis, MN, USA | 248-BD-025 |
| Recombinant Human Sonic Hedgehog/Shh (C24II) N-Terminus | R&D Systems, Minneapolis, MN, USA | 1845-SH-025/CF |
| Astrocyte Medium | ScienCell Research Laboratories, Carlsbad, CA, USA | 1801 |
| Dibutyryl cyclic-AMP sodium salt | Sigma Aldrich, St. Louis, MO, USA | D0627 |
| ATP (Adenosine 5′-triphosphate disodium salt hydrate) | Sigma Aldrich, St. Louis, MO, USA | A6419 |
| Laminin from Engelbreth-Holm-Swarm murine sarcoma basement membrane | Sigma Aldrich, St. Louis, MO, USA | L2020-1MG |
| L-Ascorbic acid | Sigma Aldrich, St. Louis, MO, USA | A4403 |
| Poly-L-ornithine hydrobromide | Sigma Aldrich, St. Louis, MO, USA | P3655 |
| Proteasome inhibitor I | Sigma Aldrich, St. Louis, MO, USA | 539160 |
| mTeSR™1 Complete Kit | StemCell Technologies, Vancouver, BC, Canada | 85857 |
| ReLeSR | StemCell Technologies, Vancouver, BC, Canada | 5872 |
| Stemolecule CHIR99021 | Stemgent, Cambridge, MA, USA | 38078 |
| Stemolecule LDN-193189 | Stemgent, Cambridge, MA, USA | 27120 |
| Stemolecule Purmorphamine | Stemgent, Cambridge, MA, USA | 39904 |
| Stemolecule SB 431542 | Stemgent, Cambridge, MA, USA | 25704-0010-10 |
| 2-mercaptoethanol (1000X) | Thermo Fisher Scientific, Waltham, MA, USA | 21985-023 |
| Advanced DMEM/F12 | Thermo Fisher Scientific, Waltham, MA, USA | 12634010 |
| B-27™ Plus Supplement (50X) | Thermo Fisher Scientific, Waltham, MA, USA | A3582801 |
| B-27™ Supplement (50X), minus vitamin A | Thermo Fisher Scientific, Waltham, MA, USA | 12587-010 |
| Fetal Bovine Serum (FBS) | Thermo Fisher Scientific, Waltham, MA, USA | 10270-106 |
| Geltrex LDEV FREE HESC QUAL | Thermo Fisher Scientific, Waltham, MA, USA | A1413302 |
| GlutaMAX (100X) | Thermo Fisher Scientific, Waltham, MA, USA | 35050-038 |
| HAM's F12 | Thermo Fisher Scientific, Waltham, MA, USA | 21765-029 |
| HEPES 1M | Thermo Fisher Scientific, Waltham, MA, USA | 15630-106 |
| KnockOut DMEM | Thermo Fisher Scientific, Waltham, MA, USA | 10829-018 |
| KnockOut™ Serum Replacement (KOSR) | Thermo Fisher Scientific, Waltham, MA, USA | 10828-010 |
| MEM Non-Essential Amino Acids Solution (100X) | Thermo Fisher Scientific, Waltham, MA, USA | 11140-035 |
| N-2 Supplement (100X) | Thermo Fisher Scientific, Waltham, MA, USA | 17502-048 |
| Neurobasal™ Medium | Thermo Fisher Scientific, Waltham, MA, USA | 21103-049 |
| Penicillin/Streptomycin | Thermo Fisher Scientific, Waltham, MA, USA | 15140-122 |
| Phosphate-Buffered Saline, pH 7.4 | Thermo Fisher Scientific, Waltham, MA, USA | 10010-015 |
| StemPro® Accutase | Thermo Fisher Scientific, Waltham, MA, USA | A1110501 |
| Y-27632 ROCK inhibitor | Tocris, Bristol, UK | 1254 |
