## Supplementary material for "Proteostasis dysregulation in p.A53T-α-Synuclein iPSC-derived astrocytes exacerbates neurodegeneration in a Parkinson’s disease model with Lewy-like pathology": SUPPL. TABLE 5

| Epitope | MANUFACTURER | CATALOG NUMBER | DILUTION |
|---|---|---|---|
| **Used in Immunofluorescence** | | | |
| ALDH1L1 | Proteintech, Rosemont, IL, USA | 17390-1-AP | 1/100 |
| Alpha Synuclein | BD Biosciences, Franklin Lakes, NJ, USA | 610787 | 1/500 |
| alpha Synuclein (phosphorylated Ser129) | FUJIFILM Wako Pure Chemical Corp, Osaka, Japan | 015-25191 | 1/1000 |
| alpha synuclein [MJFR-14-6-4-2] | Abcam, Cambridge, UK | ab209538 | 1/20,000 |
| Aquaporin 4 | Euroimmun, Lübeck, Germany | 1128-0101 | - |
| CD44 | Cell Signaling Technology, Danvers, MA, USA | 5640S | 1/1000 |
| CD49F | Biolegend, San Diego, CA, USA | 313602 | 1/200 |
| EAAT1 | Santa Cruz Biotechnology, Dallas, TX, USA | sc-515839 | 1/100 |
| FOXA2 (HNF-3β) | Santa Cruz Biotechnology, Dallas, TX, USA | sc-101060 | 1/100 |
| LAMP1 | Developmental Studies Hybridoma Bank, Iowa City, IA, USA | H4A3-s | 1/100 |
| LAMP2 | Developmental Studies Hybridoma Bank, Iowa City, IA, USA | H4A-s | 1/100 |
| LC3B | Abcam, Cambridge, UK | ab51520 | 1/200 |
| LMX1A | Millipore Sigma, Burlington, MA, USA | AB10533 | 1/2000 |
| MAP2 | Synaptic Systems, Goettingen, Germany | 188004 | 1/1000 |
| Nestin | Millipore Sigma, Burlington, MA, USA | ABD69 | 1/1000 |
| p62 | Abcam, Cambridge, UK | ab56416 | 1/250 |
| PAX6 | Developmental Studies Hybridoma Bank, Iowa City, IA, USA | AB_528427 | 1/50 |
| S100B | Abcam, Cambridge, UK | Ab41548 | 1/1000 |
| Tau (phospho T212) | Abcam, Cambridge, UK | ab4842 | 1/250 |
| Tyrosine Hydroxylase | Millipore Sigma, Burlington, MA, USA | AB152 | 1/500 |
| Ubiquitin | DAKO Agilent, Santa Clara, CA, USA | Z0458 | 1/500 |
| Vimentin | Synaptic Systems, Goettingen, Germany | 172006 | 1/500 |
| β3-Tubulin | Cell Signaling Technology, Danvers, MA, USA | 5568 | 1/1000 |
| **Used in Western Blot** | | | |
| ACTB | Proteintech, Rosemont, IL, USA | 66009-1-lg | 1/1000 |
| Alpha Synuclein | Santa Cruz Biotechnology, Dallas, TX, USA | sc-12767 | 1/1000 |
| alpha Synuclein (phosphorylated Ser129) | Abcam, Cambridge, UK | ab51253 | 1/1000 |
| GAPDH | Santa Cruz Biotechnology, Dallas, TX, USA | sc-365062 | 1/10000 |
| ITPKB | Proteintech, Rosemont, IL, USA | 12816-1-AP | 1/1000 |
| LAMP1 | Developmental Studies Hybridoma Bank, Iowa City, IA, USA | H4A3-s | 1/5000 |
| LC3B | Abcam, Cambridge, UK | ab51520 | 1/3000 |
| phospho-ULK1 (Ser757) | Cell Signaling Technology, Danvers, MA, USA | 14202 | 1/10000 |
| Ubiquitin | Cell Signaling Technology, Danvers, MA, USA | 3936 | 1/1000 |
| ULK1 | Cell Signaling Technology, Danvers, MA, USA | 8054 | 1/10000 |
