## Supplementary material for "Proteostasis dysregulation in p.A53T-α-Synuclein iPSC-derived astrocytes exacerbates neurodegeneration in a Parkinson’s disease model with Lewy-like pathology": SUPPL. TABLE 6

| GENE | FORWARD PRIMER | REVERSE PRIMER |
|---|---|---|
| CORIN | CATATCTCCATCGCCTCAGTTG | GGCAGGAGTCCATGACTGT |
| DAT | ACTTTCTCCTGTCCGTCATTG | CATGTAGAAAAGTGGCATCCC |
| EN1 | CGTGGCTTACTCCCCATTTA | TCTCGCTGTCTCTCCCTCTC |
| FOXA1 | GGGCAGGGTGGCTCCAGGAT | TGCTGACCGGGACGGAGGAG |
| FOXA2 | CCGTTCTCCATCAACAACCT | GGGGTAGTGCATCACCTGTT |
| FOXG1 | TGGCCCATGTCGCCCTTCCT | GCCGACGTGGTGCCGTTGTA |
| GBX2 | GTTCCCGCCGTCGCTGATGAT | GCCGGTGTAGACGAAATGGCCG |
| GDF7 | GACGCTGCTCAACTCCATGGCA | TTGGCGGCGTCGATGTAGAGGA |
| GIRK2 | TGTTCAGTCTTGCTCCGTTTC | TCATGGATTCTGTCAGCTTGG |
| HOXA4 | ACGCTCTGTTTGTCTGAGCGCC | AGAGGCCGAGGCCGAATTGGA |
| IRX3 | GGCTTGCGCCCCGTAGAAATGT | AGGAGCCAGGTCAGGTCCGAAC |
| LHX1 | AGGTGAAACACTTTGCTCCG | CTCCAGGGAAGGCAAACTCT |
| LHX2 | GGGCGACCACTTCGGCATGAA | CGTCGGCATGGTTGAAGTGTGC |
| LMX1A | CGCATCGTTTCTTCTCCTCT | CAGACAGACTTGGGGCTCAC |
| LMX1B | CTTAACCAGCCTCAGCGACT | TCAGGAGGCGAAGTAGGAAC |
| NTN1 | GCATGCAGGTTGCAGTTACA | GCTGCAAGCCCTTCCACTA |
| NURR1 | TCGACATTTCTGCCTTCTCCTG | GGTTCCTTGAGCCCGTGTCT |
| OTX2 | ACAAGTGGCCAATTCACTCC | GAGGTGGACAAGGGATCTGA |
| PAX5 | CCCCATTGTGACAGGCCGTGAC | TCAGCGTCGGTGCTGAGTAGCT |
| PAX6 | TGGTATTCTCTCCCCCTCCT | TAAGGATGTTGAACGGGCAG |
| PAX7 | CTTCAGTGGGAGGTCAGGTT | CAAACACAGCATCGACGG |
| S100A10 | GGCTACTTAACAAAGGAGGACC | GAGGCCCGCAATTAGGGAAA |
| SHH | CCAATTACAACCCCGACATC | AGTTTCACTCCTGGCCACTG |
| SIM1 | AAAGGGGGCCAAATCCCGGC | TCCGCCCCACTGGCTGTCAT |
| SIX3 | ACCGGCCTCACTCCCACACA | CGCTCGGTCCAATGGCCTGG |
| SNCA | TGTAGGCTCCAAAACCAAGG | CCTCCAACATTTGTCACTTGC |
| TH | TGTCTGAGGAGCCTGAGATTCG | GCTTGTCCTTGGCGTCACTG |
| VGAT | CCGAGTGGTGAACGTAGCG | GTGGCGATAATGGACCAGGAC |
| VGLUT1 | CGACGACAGCCTTTTGTGGT | GCCGTAGACGTAGAAAACAGAG |
| VMAT2 | CCCAGTGAAGACAAAGACCTC | GCAGAATCCCGCAAATATGG |
| WNT1 | GAGCCACGAGTTTGGATGTT | TGCAGGGAGAAAGGAGAGAA |
