## SUPPLEMENTAL MATERIAL for "Proteostasis dysregulation in p.A53T-α-Synuclein iPSC-derived astrocytes exacerbates neurodegeneration in a Parkinson’s disease model with Lewy-like pathology"

#### Title

[3]Pasteur International Unit for Neurodegenerative Diseases

[4]Institute for Bioinnovation, Biomedical Sciences Research Center "Alexander Fleming", Vari, 16672, Greece

[5]Membrane Traffic and Pathogenesis Unit, Department of Cell Biology and Infection, CNRS UMR 3691, Université de Paris, Institut Pasteur, Paris, 75015, France

[5]Department of Chemistry, School of Sciences, National and Kapodistrian University of Athens, Panepistimioupolis Zografou, 15772, Greece

[6]Department of Molecular Medicine and Medical Biotechnology, University of Naples Federico II, Naples, Italy

[#] Contributed equally
[$] Contributed equally
[‡] Co-senior authors
* Corresponding author

### Supplementary Figures

#### Suppl. Fig. 1

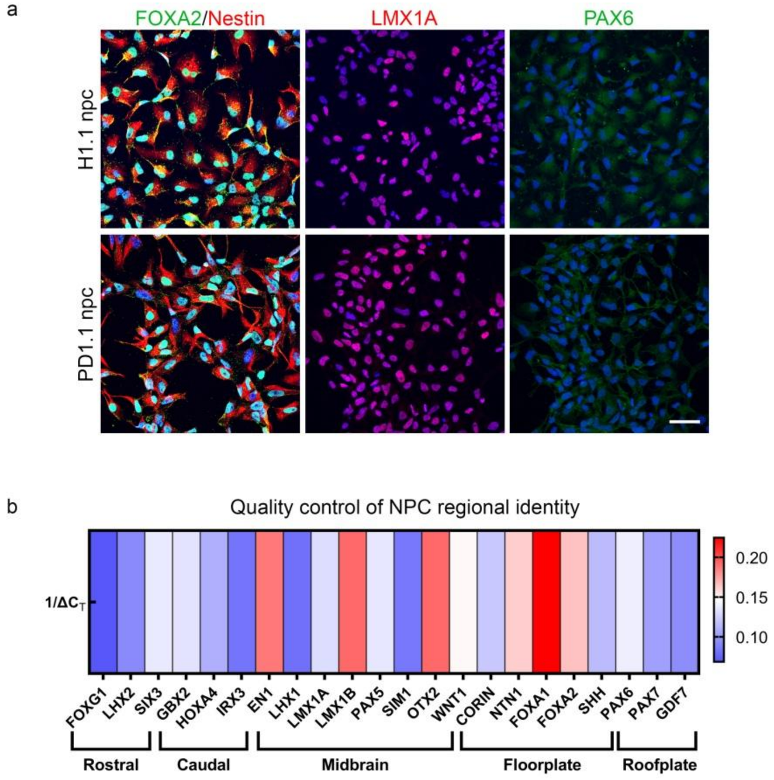


Supplementary Figure 1| Quality control of ventral midbrain-patterned NPCs

**a** Representative confocal images of healthy and PD iPSC-derived NPCs positively immunostained for midbrain/floorplate markers: FOXA2/Nestin (NPC marker), LMX1A, and roofplate marker PAX-6 (negative) with DAPI. Scale bar, 40 μm. **b** Quality control of midbrain-patterned NPCs assessed by RT-qPCR. A heatmap shows the calculated 1/ΔCT values for genes associated with rostral, caudal, midbrain, floorplate, and roofplate fates, as previously described[1]. Acceptable cell lines exhibit relatively higher expression of midbrain and floorplate-specific genes.

*DAPI, 4',6-diamidino-2-phenylindole; FOXA2, Forkhead box A2; LMX1A, LIM homeobox transcription factor 1 alpha; NPC, neural precursor cell; PAX-6, Paired box protein Pax-6; PD, Parkinson's disease.*

Suppl. Fig. 2

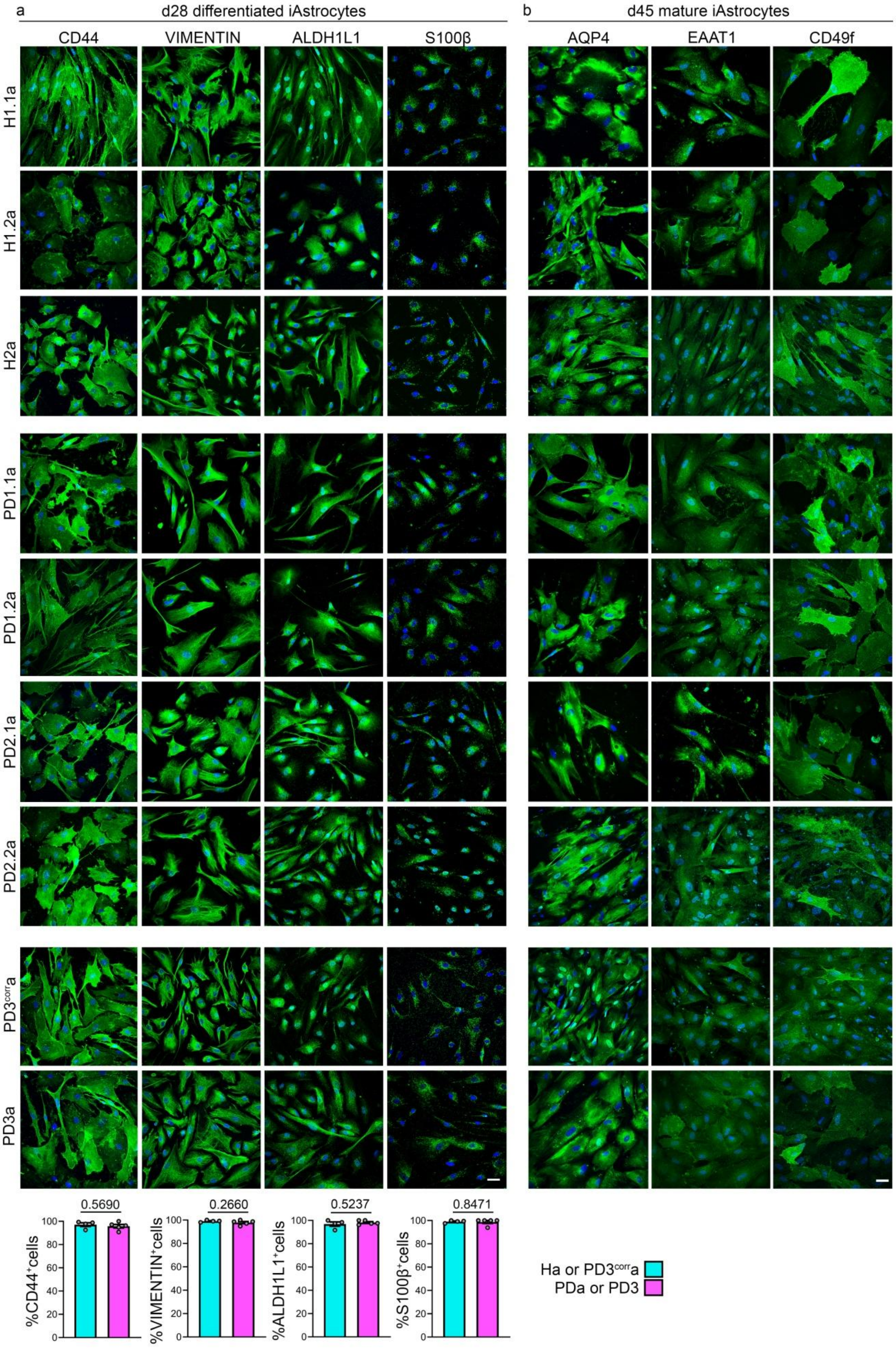
a
d28 differentiated iAstrocytes
CD44
VIMENTIN
ALDH1L1
S100β
b
d45 mature iAstrocytes
AQP4
EAAT1
CD49f
H1.1a
H1.2a
H2a
PD1.1a
PD1.2a
PD2.1a
PD2.2a
PD3corra
PD3a
0.5690
0.2660
0.5237
0.8471
%CD44+cells
%VIMENTIN+cells
%ALDH1L1+cells
%S100β+cells
Ha or PD3corra
PDa or PD3

Supplementary Figure 2| Expression of typical astrocytic markers in differentiated iPSC-derived astrocytes.

**a** Representative confocal images of iPSC-derived astrocytes after 28 days of differentiation, showing expression of typical astrocytic markers CD44, Vimentin, ALDH1L1, and S100β. Bar plots show the quantification of the percentage of cells expressing each marker relative to the total cell population (DAPI). **b** Representative confocal images of iPSC-derived astrocytes after 28 days of differentiation, followed by an additional 15 days of maturation, showing expression of more mature astrocytic markers: AQP4, EAAT1, and CD49f. All Ha and PDa lines (non-isogenic and isogenic) used in this study are presented in both panels. Scale bars: 40 μm.

Data are presented as mean ± SEM; Unpaired two-tailed t-test was used for comparisons; Sample sizes: n = 4 Ha and n = 5 PDa lines, including the isogenic pair.

*AQP4, Aquaporin-4; CD44, Cluster of Differentiation 44; CD49f, Integrin alpha-6; DAPI, 4′,6-diamidino-2-phenylindole; EAAT1, Excitatory amino acid transporter 1; Ha, healthy astrocytes; PDa, p.A53T-αSyn astrocytes; S100β, S100 calcium-binding protein B; Vimentin, intermediate filament protein.*

Suppl. Fig. 3

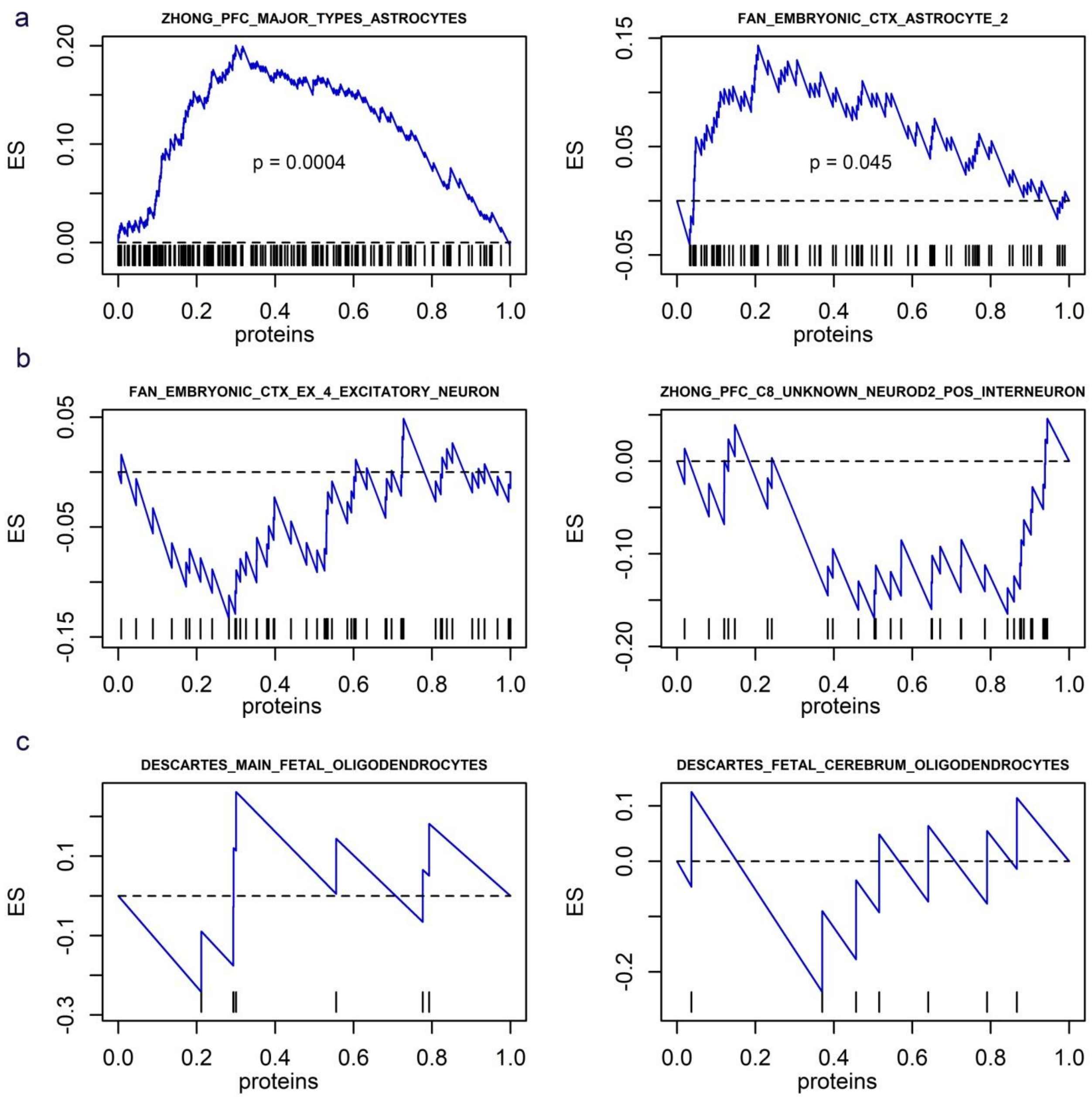


Supplementary Figure 3| Proteomic analysis for astrocytic characterization.

Mountain plots from Gene Set Enrichment Analysis (GSEA) of the average proteomic profile. **a** The plots demonstrate a significant overlap between highly expressed proteins in our data and two astrocyte-related gene sets. **b** No significant overlap is observed with brain neuron-related gene sets, nor **c** with oligodendrocyte-related gene sets. The brain neuron-related and oligodendrocyte-related gene sets shown were selected at random, as no such gene sets in the tested database were found to be significant.

Suppl. Fig. 4

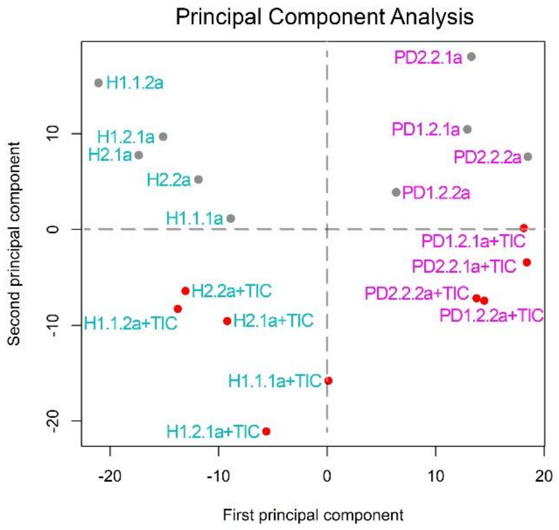


Supplementary Figure 4| Proteomic analysis demonstrating astrocytic response to TIC treatment

Principal Component Analysis (PCA) plot for the full dataset (Ha and PDa, untreated and TIC treated) after batch and cell cycle correction (see methods). Grey dots correspond to untreated samples; Red dots correspond to TIC-treated samples. Ha samples are designated with cyan and PDa samples with magenda fonts. Clustering of samples based on the presence of the mutation (first principal component) and on TIC-treatment (second principal component) is apparent.

*Ha, healthy astrocytes; PDa, p.A53T-αSyn astrocytes; TIC, TNFα, IL-1α and C1q treatment.*

Suppl. Fig. 5

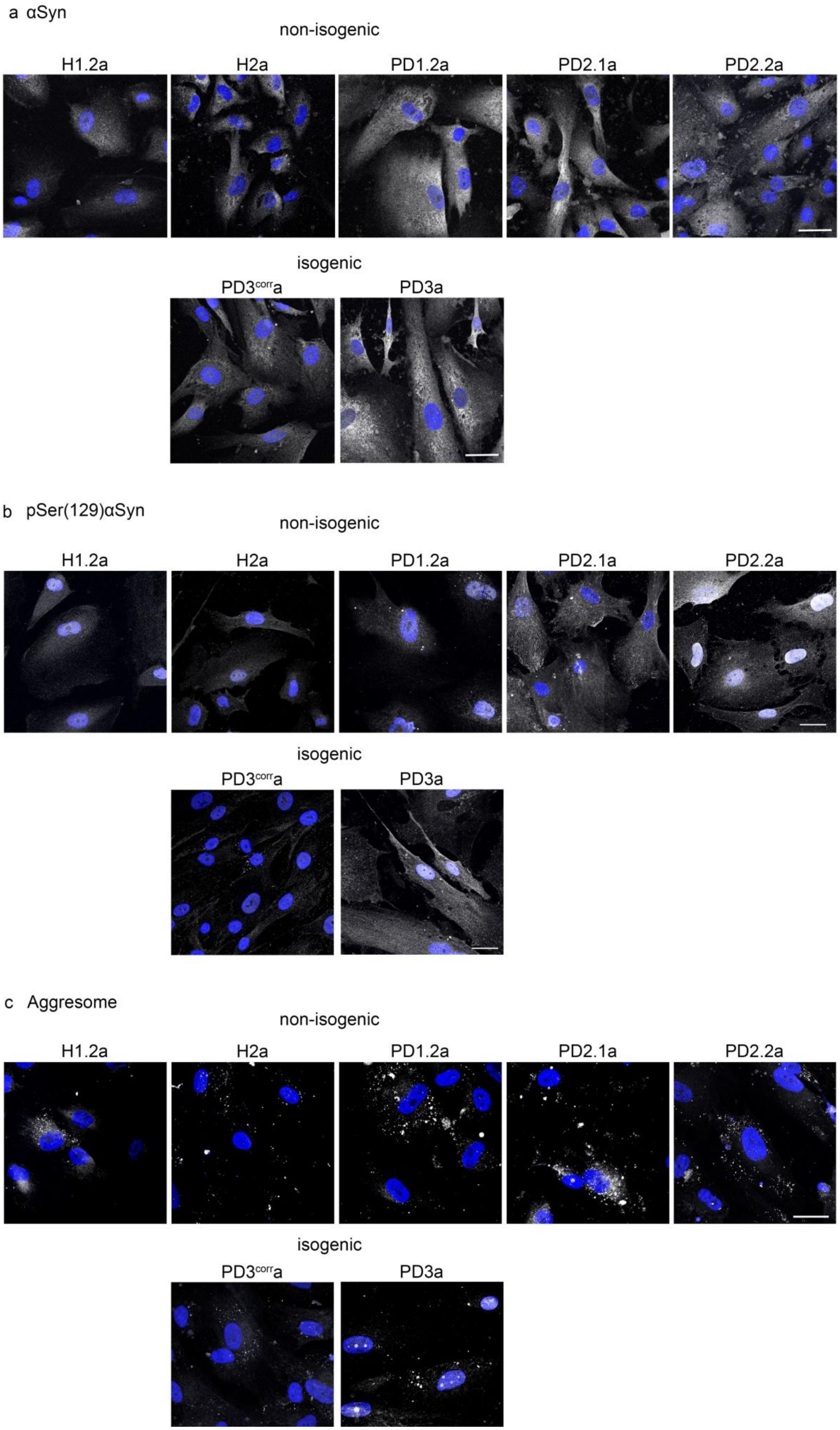
a αSyn
non-isogenic
H1.2a
H2a
PD1.2a
PD2.1a
PD2.2a
isogenic
PD3corra
PD3a
b pSer(129)αSyn
non-isogenic
H1.2a
H2a
PD1.2a
PD2.1a
PD2.2a
isogenic
PD3corra
PD3a
c Aggresome
non-isogenic
H1.2a
H2a
PD1.2a
PD2.1a
PD2.2a
isogenic
PD3corra
PD3a

Supplementary Figure 5| Upregulation of αSyn and increased p(Ser129)αSyn and protein aggregation in p.A53T-αSyn astrocytes (supplemental to Figure 2).

Representative confocal images of non-isogenic and isogenic ($PD3^{corr}a$ and PD3a) lines immunostained for (**a)** αSyn and (**b**) p(Ser129)αSyn with DAPI nuclear counterstain. Scale bars: 30 μm. **c** Representative confocal images of non-isogenic and isogenic ($PD3^{corr}a$ and PD3a) lines after Aggresome staining with DAPI. Scale bar: 30 μm.

*αSyn, alpha-synuclein; p(Ser129)αSyn, phosphorylated alpha-synuclein at serine 129; GAPDH, glyceraldehyde 3-phosphate dehydrogenase; Ha, healthy astrocytes; PDa, p.A53T-αSyn astrocytes;* $PD3^{corr}a$*, corrected isogenic astrocytes; PD3a, p.A53T-αSyn astrocytes.*

Suppl. Fig. 6

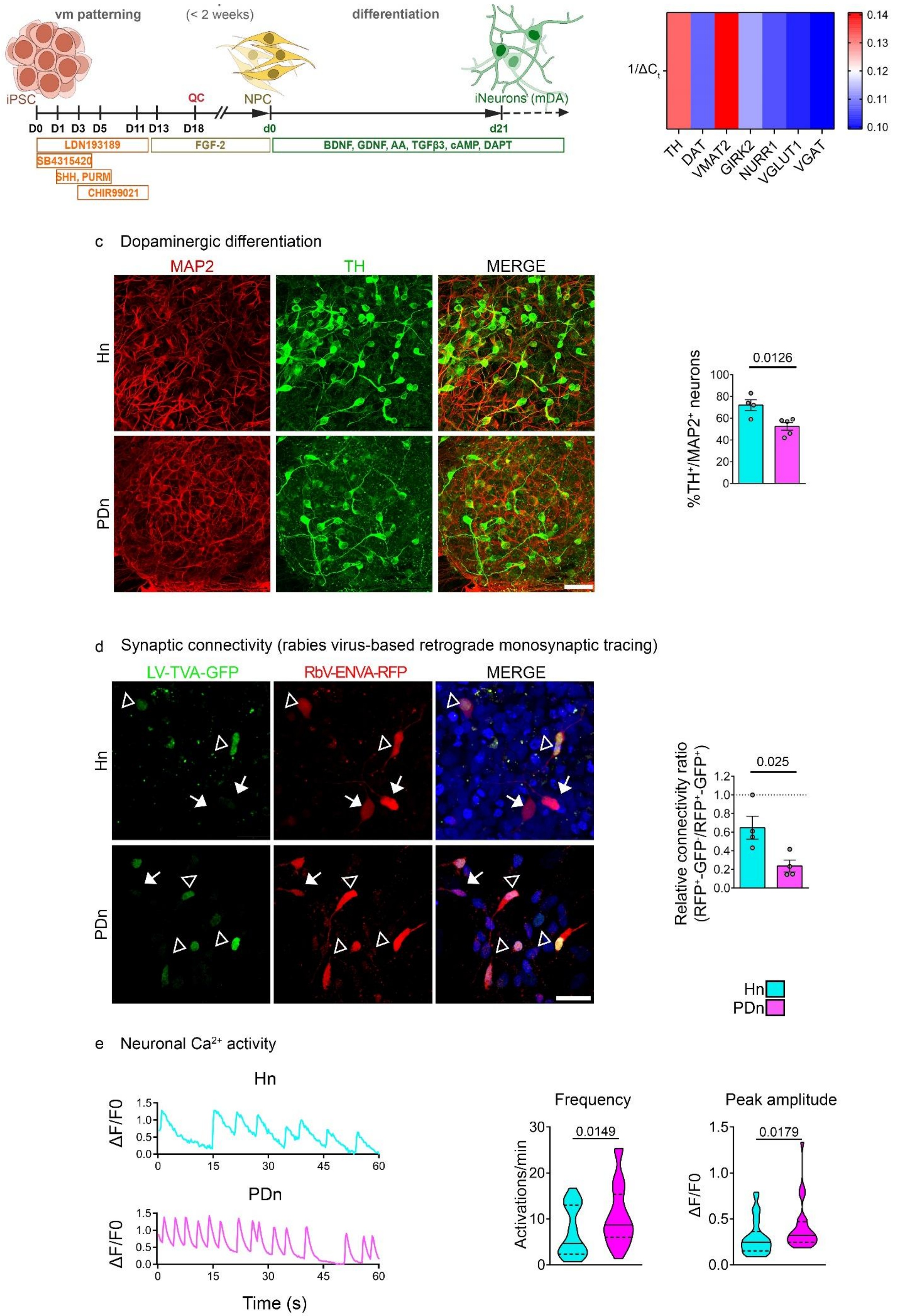
vm patterning
(< 2 weeks)
differentiation
iPSC
QC
NPC
iNeurons (mDA)
D0 D1 D3 D5 D11 D13 D18
d0
d21
LDN193189
SB4315420
SHH, PURM
CHIR99021
FGF-2
BDNF, GDNF, AA, TGFβ3, cAMP, DAPT
1/ΔCt
TH DAT VMAT2 GIRK2 NURR1 VGLUT1 VGAT
0.14 0.13 0.12 0.11 0.10
c Dopaminergic differentiation
MAP2
TH
MERGE
Hn
PDn
%TH+/MAP2+ neurons
0.0126
d Synaptic connectivity (rabies virus-based retrograde monosynaptic tracing)
LV-TVA-GFP
RbV-ENVA-RFP
MERGE
Relative connectivity ratio (RFP+-GFP-/RFP+-GFP+)
0.025
Hn
PDn
e Neuronal Ca2+ activity
ΔF/F0
Time (s)
Frequency
Activations/min
0.0149
Peak amplitude
0.0179

Supplementary Figure 6| Molecular and functional characterization of iPSC-derived dopaminergic neurons.

**a** Schematic illustration of the iPSC differentiation protocol toward ventral midbrain dopaminergic neurons. **b** Heatmap depicting relative mRNA levels of key neuronal markers (TH, DAT, VMAT2, GIRK2, NURR1, VGLUT1, VGAT) detected by RT-qPCR in iPSC-derived neuronal monocultures. Data are averages from 3 independent differentiations of both healthy and PD iPSC lines. **c** Representative confocal images showing immunostaining for MAP2 (neuronal marker) and TH (dopaminergic neuron marker) in Hn and PDn neuronal cultures at d21. Scale bar, 30 μm. The accompanying bar plot quantifies the dopaminergic neuron ($TH^+$) percentage relative to total neurons ($MAP2^+$). **d** Representative confocal images of rabies-virus based retrograde monosynaptic tracing in day-40 of neuronal differentiation healthy (Hn, including $PD3^{corr}$n) and p.A53T-αSyn (PDn) neuronal cultures. Target (presynaptic) neurons are RbV-ENVA-$RFP^+$/LV-TVA-$GFP^-$ (solid arrows) and starter (postsynaptic) neurons are RbV-ENVA-$RFP^+$/LV-TVA-$GFP^+$ (empty arrowheads). Scale bar, 30 μm. The bar plot represents the quantification of the relative connectivity ratio. **e** Representative Fluo4AM traces showing spontaneous $Ca^{2+}$ activity of one Hn, and one PDn at day-35 of neuronal differentiation. Quantification of frequency and peak amplitude of $Ca^{2+}$ oscillations.

Data are presented as mean ± SEM (c, d) or median with interquartile range (e); Unpaired two-tailed t-test was used for comparisons in c, d. Mann-Whitney test was used to compare frequency data in e; Sample sizes: c, n = 4 (3 Hn and $PD3^{corr}$n) and 5 (4 PDn and PD3n); d, n = 4 in both Hn (incl. $PD3^{corr}$n) and PDn, all from independent experiments [Note: Hn samples have been reused in Fig. 1c, in a different context]; e, *n* = 37 Hn and 31 PDn from one non-isogenic pair.

*AUC, area under the curve; DAT, dopamine transporter; Fluo4AM, Fluo-4 acetoxymethyl ester; GAPDH, glyceraldehyde 3-phosphate dehydrogenase; GIRK2, G protein-coupled inward rectifier potassium channel 2; iPSC, induced pluripotent stem cell; LV, lentivirus; MAP2, microtubule-associated protein 2; NDIV, neuronal day in vitro; NURR1, nuclear receptor related 1 protein; p(Ser129)αSyn, phosphorylated alpha-synuclein at serine 129; RbV, rabies virus; SEM, standard error of the mean; TH, tyrosine hydroxylase; VGAT, vesicular GABA transporter; VGLUT1, vesicular glutamate transporter 1; VMAT2, vesicular monoamine transporter 2; Hn, healthy neurons; PDn, p.A53T-αSyn neurons.*

Suppl. Fig. 7

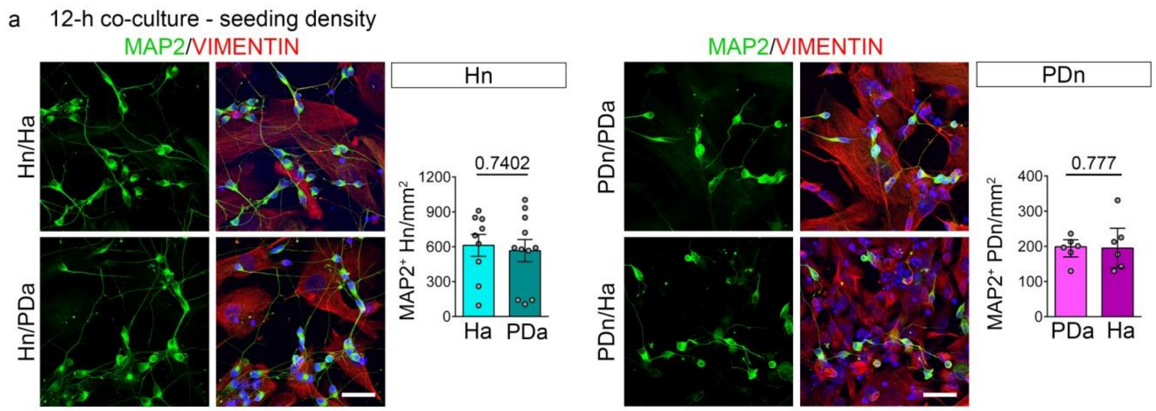


Supplementary Figure 7| Seeding density of iNeurons and iAstrocytes in co-culture.

**a** Representative confocal images of 12-h co-cultures of Hn, on Ha or PDa, and PDn on PDa or Ha, immunostained for the neuronal marker MAP2 and the astrocytic marker Vimentin, with DAPI counterstain. Scale bar = 30 μm. Bar plots show quantification of the neuronal seeding density, i.e., the number of $MAP2^+$ neurons per area, across all combinations.

Data are presented as mean ± SEM (a). For comparisons, unpaired two-tailed t-test was used. Sample sizes: a, n = 9 (Hn/Ha) and 11 (Hn/PDa) FOV from two separate experiments; n = 6 FOV from one co-culture of PDn with PDa or Ha.

*DAPI, 4',6-diamidino-2-phenylindole; FOV, field of view; Ha, healthy astrocytes; Hn, healthy neurons; MAP2, microtubule-associated protein 2; NDIV, neuronal day in vitro; PDa, p.A53T-αSyn astrocytes; PDn, p.A53T-αSyn neurons.*

Suppl. Fig. 8

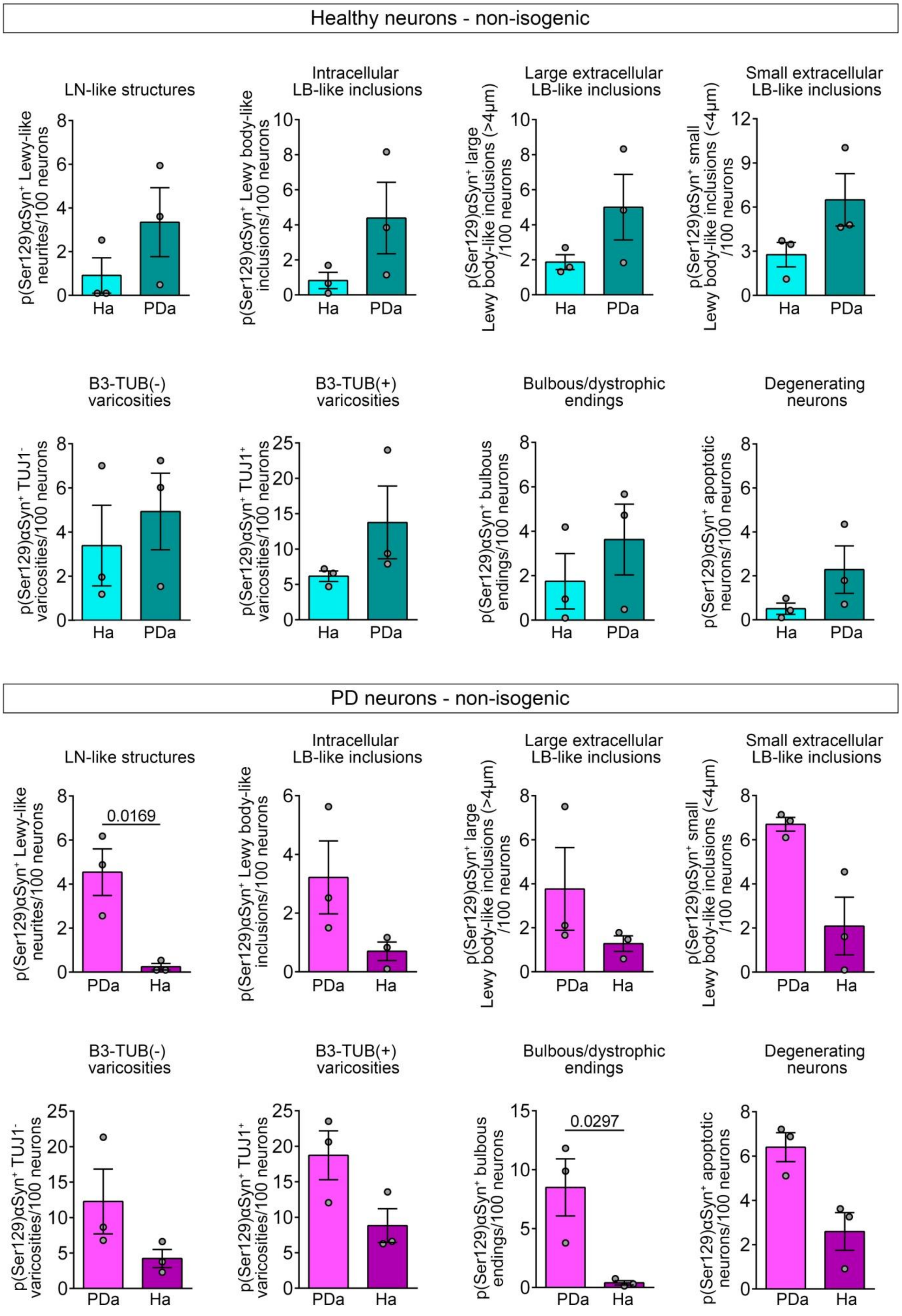
Healthy neurons - non-isogenic
LN-like structures
Intracellular LB-like inclusions
Large extracellular LB-like inclusions
Small extracellular LB-like inclusions
p(Ser129)αSyn⁺ Lewy-like neurites/100 neurons
p(Ser129)αSyn⁺ Lewy body-like inclusions/100 neurons
p(Ser129)αSyn⁺ large Lewy body-like inclusions (>4μm) /100 neurons
p(Ser129)αSyn⁺ small Lewy body-like inclusions (<4μm) /100 neurons
Ha
PDa
B3-TUB(-) varicosities
B3-TUB(+) varicosities
Bulbous/dystrophic endings
Degenerating neurons
p(Ser129)αSyn⁺ TUJ1⁻ varicosities/100 neurons
p(Ser129)αSyn⁺ TUJ1⁺ varicosities/100 neurons
p(Ser129)αSyn⁺ bulbous endings/100 neurons
p(Ser129)αSyn⁺ apoptotic neurons/100 neurons
PD neurons - non-isogenic
0.0169
0.0297

Supplementary Figure 8| Quantification of PD-relevant neuropathological features in co-cultures (non-isogenic, supplemental to Figure 5).

Bar plots showing each separate quantification of eight neuropathology-related features identified in 14-day co-cultures of Hn on Ha or PDa, and of PDn on PDa or Ha, after immunostaining for pSer(129)αSyn and B3-tubulin. The quantified pathological features include: Lewy neurite (LN)-like structures, Intracellular Lewy body (LB)-like inclusions, large extracellular LB-like inclusions, small extracellular LB-like inclusions, B3-tubulin$^-$ axonal varicosities, B3-tubulin$^+$ axonal varicosities, bulbous/dystrophic axonal endings, and degenerating neurons. Data are presented as mean ± SEM; n = 3 Hn and 3 PDn non-isogenic lines, from separate experiments. Ratio-paired two-tailed t-test was used for group comparisons. *Hn, Healthy neurons; Ha, Healthy astrocytes; PDa, p.A53T-αSyn astrocytes; PDn, p.A53T-αSyn neurons; LN, Lewy neurite; LB, Lewy body; SEM, standard error of the mean.*

Suppl. Fig. 9

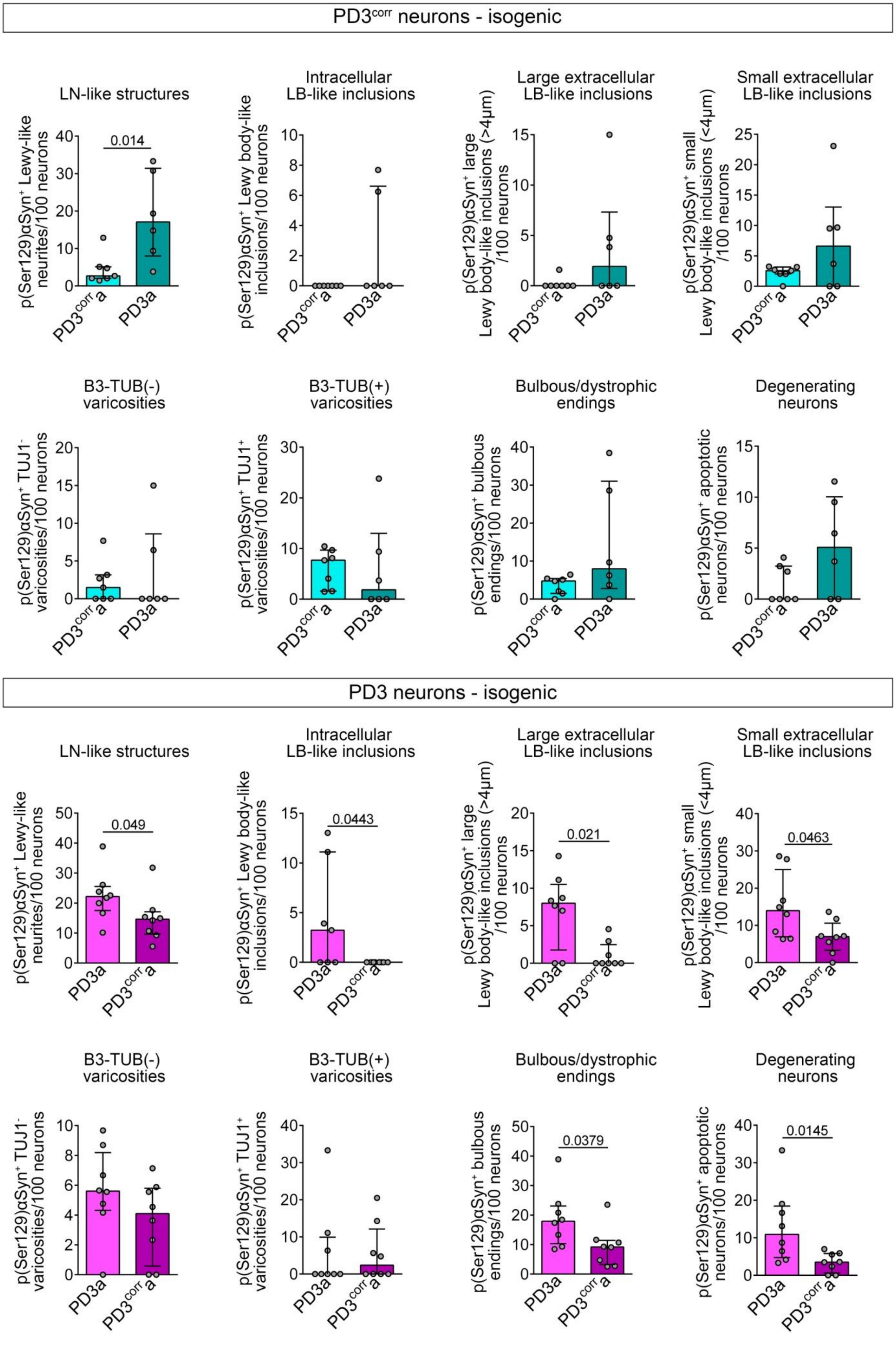
PD3corr neurons - isogenic
LN-like structures
p(Ser129)αSyn+ Lewy-like neurites/100 neurons
0.014
Intracellular LB-like inclusions
p(Ser129)αSyn+ Lewy body-like inclusions/100 neurons
Large extracellular LB-like inclusions
p(Ser129)αSyn+ large Lewy body-like inclusions (>4μm) /100 neurons
Small extracellular LB-like inclusions
p(Ser129)αSyn+ small Lewy body-like inclusions (<4μm) /100 neurons
B3-TUB(-) varicosities
p(Ser129)αSyn+ TUJ1- varicosities/100 neurons
B3-TUB(+) varicosities
p(Ser129)αSyn+ TUJ1+ varicosities/100 neurons
Bulbous/dystrophic endings
p(Ser129)αSyn+ bulbous endings/100 neurons
Degenerating neurons
p(Ser129)αSyn+ apoptotic neurons/100 neurons
PD3corr a
PD3a
PD3 neurons - isogenic
LN-like structures
0.049
Intracellular LB-like inclusions
0.0443
Large extracellular LB-like inclusions
0.021
Small extracellular LB-like inclusions
0.0463
B3-TUB(-) varicosities
B3-TUB(+) varicosities
Bulbous/dystrophic endings
0.0379
Degenerating neurons
0.0145

Supplementary Figure 9| Quantification of PD-relevant neuropathological features in co-cultures (isogenic pair - supplemental to Figure 5).

Bar plots showing the separate quantification of eight neuropathology-related features identified in 14-day co-cultures of the isogenic pair across all combinations, i.e., PD3$^{corr}$n, on PD3$^{corr}$a or PD3a, and PD3n on PD3a or PD3$^{corr}$a, after immunostaining for pSer(129)$\alpha$Syn and B3-tubulin. The quantified pathological features include: Lewy neurite (LN)-like structures, Intracellular Lewy body (LB)-like inclusions, large extracellular LB-like inclusions, small extracellular LB-like inclusions, B3-tubulin$^{-}$ axonal varicosities, B3-tubulin$^{+}$ axonal varicosities, bulbous/dystrophic axonal endings, and degenerating neurons. Data are presented as mean ± SEM; n = 7 FOV (PD3$^{corr}$n/PD3$^{corr}$a); 6 FOV (PD3$^{corr}$n/ PD3a); 8 FOV (PD3n/PD3a); 8 FOV (PD3n/PD3$^{corr}$a). Unpaired two-tailed t-test was used for comparisons.

*LN, Lewy neurite; LB, Lewy body; FOV, field of view; PD3a, p.A53T-$\alpha$Syn astrocytes; PD3$^{corr}$a, isogenic control astrocytes; PD3$^{corr}$n, corrected isogenic neurons; PD3n, p.A53T-$\alpha$Syn neurons.*

#### Supplementary Tables

Supplementary Table 1| Summary of patient-derived cell lines used in this study

| | Donor status | iPSC line coding | Age at biopsy | aSyn mutation | Sex | Source/reference |
|---|---|---|---|---|---|---|
| Non-isogenic | Healthy (non-PD control) | H1.1 | 41 | Wild Type SNCA | M | Kouroupi et al., 2017 |
| | | H1.2 | | | | |
| | Healthy (non-PD control) | H2 | 45 | Wild Type SNCA | M | New York Stem Cell Foundation |
| | Parkinson's Disease Patient | PD1.1 | 49 | G209A SNCA | M | Kouroupi et al., 2017 |
| | | PD1.2 | | | | |
| | Parkinson's Disease Patient | PD2.1 | 40 | G209A SNCA | M | Kouroupi et al., 2017 |
| | | PD2.2 | | | | |
| Isogenic pair | Parkinson's Disease Patient | PD3 | 49 | G209A SNCA | F | Soldner et al., 2011 |
| | Parkinson's Disease Patient (gene-corrected control) | $PD3^{corr}$ | 49 | G209A SNCA corrected | F | Soldner et al., 2011 |

Supplementary Table 2| iAstrocyte Samples included in proteomic analysis

| Donor status | aSyn mutation | Sex | iPSC line coding | Independent differentiations |
|---|---|---|---|---|
| Healthy (non-PD) | Wild Type SNCA | M | H1.1 | H1.1.1 |
| | | | | H1.1.2 |
| | | | H1.2 | H1.2.1 |
| | | | | H1.2.2 |
| Healthy (non-PD) | Wild Type SNCA | M | H2 | H2.1 |
| | | | | H2.2 |
| PD | G209A SNCA | M | PD1.2 | PD1.2.1 |
| | | | | PD1.2.2 |
| PD | G209A SNCA | M | PD2.2 | PD2.2.1 |
| | | | | PD2.2.2 |
